# Population codes enable learning from few examples by shaping inductive bias

**DOI:** 10.1101/2021.03.30.437743

**Authors:** Blake Bordelon, Cengiz Pehlevan

## Abstract

Learning from a limited number of experiences requires suitable inductive biases. To identify how inductive biases are implemented in and shaped by neural codes, we analyze sample-efficient learning of arbitrary stimulus-response maps from arbitrary neural codes with biologically-plausible readouts. We develop an analytical theory that predicts the generalization error of the readout as a function of the number of observed examples. Our theory illustrates in a mathematically precise way how the structure of population codes shapes inductive bias, and how a match between the code and the task is crucial for sample-efficient learning. We observe that many different codes can support the same inductive bias. By analyzing recordings from the mouse primary visual cortex, we demonstrate that biological codes have lower total activity than other codes with identical bias. Using these mouse primary visual cortex responses, we demonstrate the existence of an efficiency bias towards low frequency orientation discrimination tasks for grating stimuli and low spatial frequency reconstruction tasks for natural images. We reproduce the discrimination bias in a simple model of primary visual cortex, and further show how invariances in the code to certain stimulus variations alter learning performance. We extend our methods to time-dependent neural codes and predict the sample efficiency of readouts from recurrent networks. Finally, we discuss implications of our theory in the context of recent developments in neuroscience and artificial intelligence. Overall, our study provides a concrete method for elucidating inductive biases of the brain and promotes sample-efficient learning as a general normative coding principle.

## 1 Introduction

The ability to learn quickly is crucial for survival in a complex and an everchanging environment, and the brain effectively supports this capability. Often, only a few experiences are sufficient to learn a task, whether acquiring a new word [1] or recognizing a new face [2]. Despite the importance and ubiquity of sample efficient learning, our understanding of the brain’s information encoding strategies that support this faculty remains poor [3, 4, 5].

In particular, when learning and generalizing from past experiences, and especially from few experiences, the brain relies on implicit assumptions it carries about the world, or its inductive biases [6, 5]. Reliance on inductive bias is not a choice: inferring a general rule from finite observations is an ill-posed problem which requires prior assumptions since many hypotheses can explain the same observed experiences [7]. Consider learning a rule that maps photoreceptor responses to a prediction of whether an observed object is a threat or is neutral. Given a limited number of visual experiences of objects and their threat status, many threat-detection rules are consistent with these experiences. By choosing one of these threat-detection rules, the nervous system reveals an inductive bias. Without the right biases that suit the task at hand, successful generalization is impossible [6, 5]. In order to understand why we can quickly learn to perform certain tasks accurately but not others, we must understand the brain’s inductive biases [3, 4, 5].

In this paper, we study sample efficient learning and inductive biases in a general neural circuit model which comprises of a population of sensory neurons and a readout neuron learning a stimulus-response map with a biologically-plausible learning rule (Fig 1A). For this circuit and learning rule, inductive bias arises from the nature of the neural code for sensory stimuli, specifically its similarity structure. While different population codes can encode the same stimulus variables and allow learning of the same output with perfect performance given infinitely many samples, learning performance can depend dramatically on the code when restricted to a small number of samples, where the reliance on and the effect of inductive bias are strong (Fig 1B,C,D). Given the same sensory examples and their associated response values, the readout neuron may make drastically different predictions depending on the inductive bias set by the nature of the code, leading to successful or failing generalizations (Fig 1C,D). We say that a code and a learning rule, together, have a good inductive bias for a task if the task can be learned from a small number of examples.

**Figure 1:**
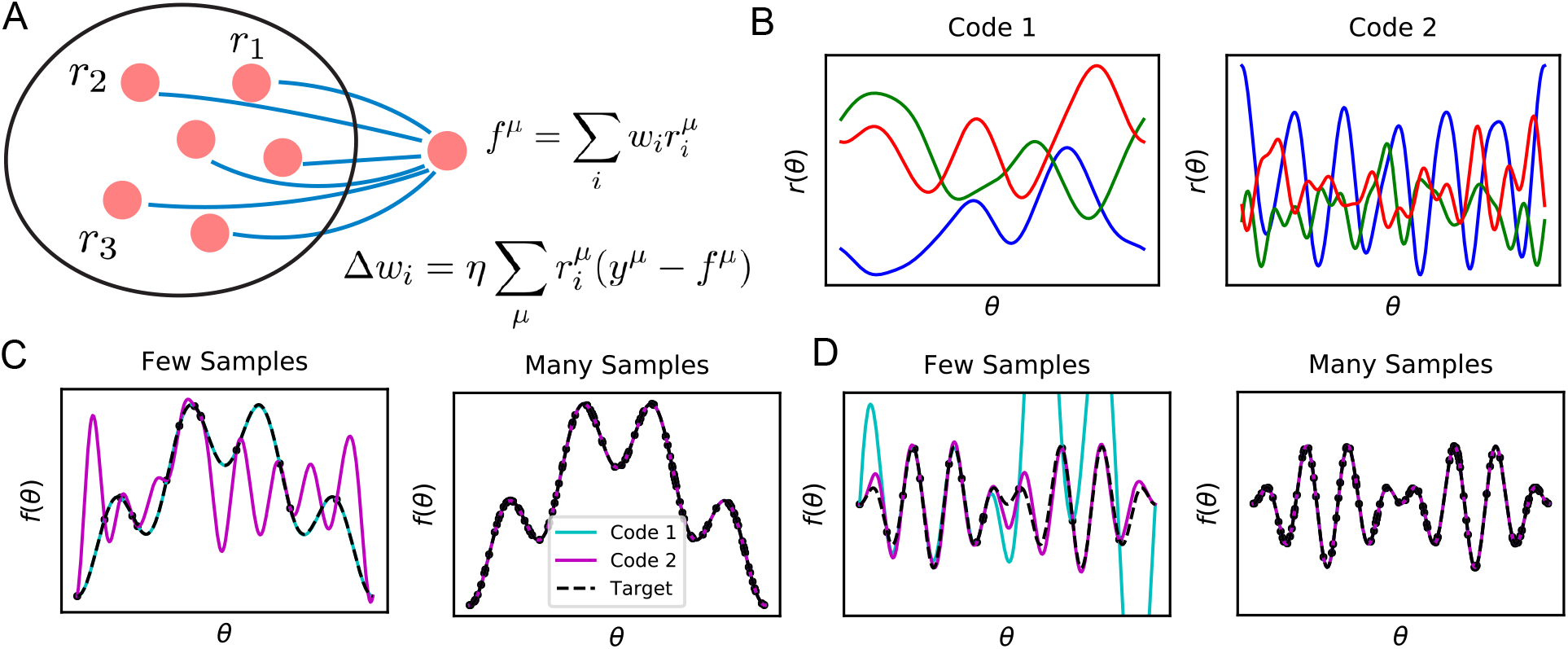
Learning tasks through linear readouts exploit representations of the population code to approximate a target response. **A** The readout weights from the population to a downstream neuron, shown in blue, are updated to fit target values *y,* using the local, biologically plausible delta rule. **B** Examples of tuning curves for two different population codes: Smooth tuning curves (Code 1) and rapidly varying tuning curves (Code 2). **C** (Left) A target function with low frequency content is approximated through the learning rule shown in **A** using these two codes. The readout from Code 1 (turquoise) fits the target function (black) almost perfectly with only *P* =12 training examples, while readout from Code 2 (purple) does not accurately approximate the target function. (Right) However, when the number of training examples is sufficiently large (*P* = 120), the target function is estimated perfectly by both codes, indicating that both codes are equally expressive. **D** The same experiment is performed on a task with higher frequency content. (Left) Code 1 fails to perform well with *P* = 12 samples indicating mismatch between inductive bias and the task can prevent sample efficient learning while Code 2 accurately fits the target. (Right) Again, provided enough data *P* = 120, both models can accurately estimate the target function. Details of these simulations are given in Methods 4.1.

In order to understand how population codes shape inductive bias and allow fast learning of certain tasks over others with a biologically plausible learning rule, we develop an analytical theory of the readout neuron’s learning performance as a function of the number of sampled examples, or sample size. We find that the readout’s performance is completely determined by the code’s kernel, a function which takes in pairs of population response vectors and outputs a representational similarity defined by the inner product of these vectors. We demonstrate that the spectral properties of the kernel introduce an inductive bias toward explaining sampled data with simple stimulus-response maps and determine compatibility of the population code with learning task, and hence the sample-efficiency of learning. We observe that many codes could support the same kernel function, however, by analyzing data from mouse primary visual cortex (V1) [8, 9, 10, 11], we find that the biological code is metabolically more efficient than others. Further, mouse V1 responses support sample-efficient learning of low frequency orientation discrimination and low spatial frequency reconstruction tasks over high frequency ones. We demonstrate the discrimination bias in a simple model of V1 and show how response nonlinearity, sparsity, and relative proportion of simple and complex cells influence the code’s bias and performance on learning tasks, including ones that involve invariances. Finally, we extend our theory to temporal population codes. As an example, we study codes generated by recurrent neural networks learning a delayed response task.

Overall, our results demonstrate that for a fixed learning rule, the neural sensory representation imposes an inductive bias over the space of learning tasks, allowing some tasks to be learned by a downstream neuron more sample-efficiently than others. Our work provides a concrete method for elucidating inductive biases of populations of neurons and suggest sample-efficient learning as a novel functional role for population codes.

## 2 Results

We consider a population of *N* neurons whose responses, {*r*_1_(***θ***), *r*_2_(***θ***),…, *r_N_*(***θ***)}, vary with the input stimuli, which is parameterized by a vector variable **θ**, such as the orientation and the phase of a grating (Figure 1A). These responses define the population code. A readout neuron learns its weights **w** to approximate a stimulus-response map, or a target function *y*(***θ***), such as one that classifies stimuli as apetitive (*y* = 1) or aversive (*y* = −1), or a more smooth one that attaches intermediate values of valence. We emphasize that in our model only the readout neuron performs learning, and the population code is assumed to be static. Our theory is general in its assumptions about the structure of the population code and the stimulus-response map considered (Methods 4.2), and can apply to many scenarios.

The readout neuron learns from *P* stimulus-response examples with the goal of generalizing to previously unseen ones. Example stimuli ***θ**^μ^*, (*μ* = 1,…,*P*) are sampled from a probability distribution describing stimulus statistics *p*(***θ***). This distribution can be natural or artificially created, for example, for a laboratory experiment (Appendix, App. 1). From the set of learning examples, 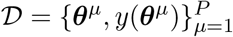, the readout weights are learned with the local, biologically-plausible delta-rule, Δ*w_j_* = *η*∑*r_μ_ r_j_*(***θ**^μ^*)(*y*(***θ**^μ^*) – **r**(***θ**^μ^*) · **w**), where *η* is a learning rate (Figure 1A). This learning process, when weights are initialized as **w**_0_ = 0, converges to a unique set of weights 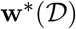 (App. 3A). Generalization error with these weights is given by

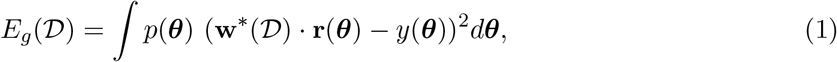

which quantifies the expected error of the trained readout over the entire stimulus distribution *p*(***θ***). This quantity will depend on the population code **r**(***θ***), the target function y(***θ***) and the set of training examples 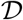. Our theoretical analysis of this model provides insights into how populations of neurons encode information and allow sample-efficient learning.

### Kernel structure of population codes controls learning performance

First, we note that the generalization performance of the learned readout on a given task depends entirely on the inner product kernel, defined by 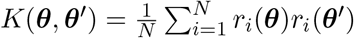, which quantifies the similarity of population responses to two different stimuli ***θ*** and ***θ***′. This is because the learning procedure converges to a unique solution 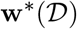 for the training set 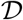 [12, 13] and the readout neuron’s learned output has the form

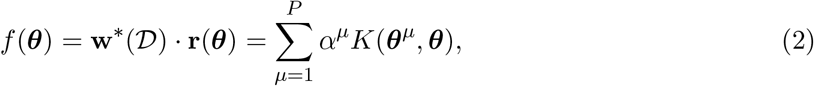

where the coefficient vector ***α*** = **K**^+^**y**, where ^+^ denotes Moore-Penrose inverse (App. 3A), and the matrix **K** has entries *K_μν_* = *K*(***θ**^μ^, **θ**^ν^*). In these expressions the population code only appears through the kernel *K*, showing that the kernel alone controls the learned response pattern. This result applied also to nonlinear readouts (App. 3G), showing that the kernel can control the learned solution in a variety of cases.

### Biological codes are metabolically more efficient and more selective than other codes with identical kernels

The fact that learning performance depends only on the kernel introduces a large degeneracy in the set of codes which achieve identical desired performance on learning tasks. This is because the kernel is invariant with respect to left-rotations of the population code. An orthogonal transformation **Q** applied to a population code **r**(***θ***) generates a new code 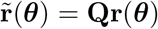 with an identical kernel. Such orthogonal transformations are the most general linear transformations which preserve the kernel for all possible pairs of vectors **x**, **y** ∈ ℝ^*N*^. For rank deficient neural codes, other linear transformations can preserve the kernel, however, for any linear transformation of the code 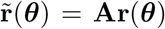 which preserves the kernel, there exists an orthogonal matrix **Q** such that 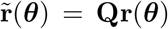 (App. 2A). It therefore suffices to consider orthogonal matrices **Q**. Codes **r**(***θ***) and 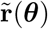 will have identical readout performance on all possible learning tasks. We illustrate this degeneracy in Figure 2 using a publicly available dataset which consists of activity recorded from ~ 20,000 neurons from the primary visual cortex of a mouse while shown static gratings [8, 9]. An original code **r**(***θ***) is rotated to generate 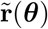 (Figure 2A) which have the same kernels (Figure 2B) and the same performance on a learning task (Figure 2C).

**Figure 2:**
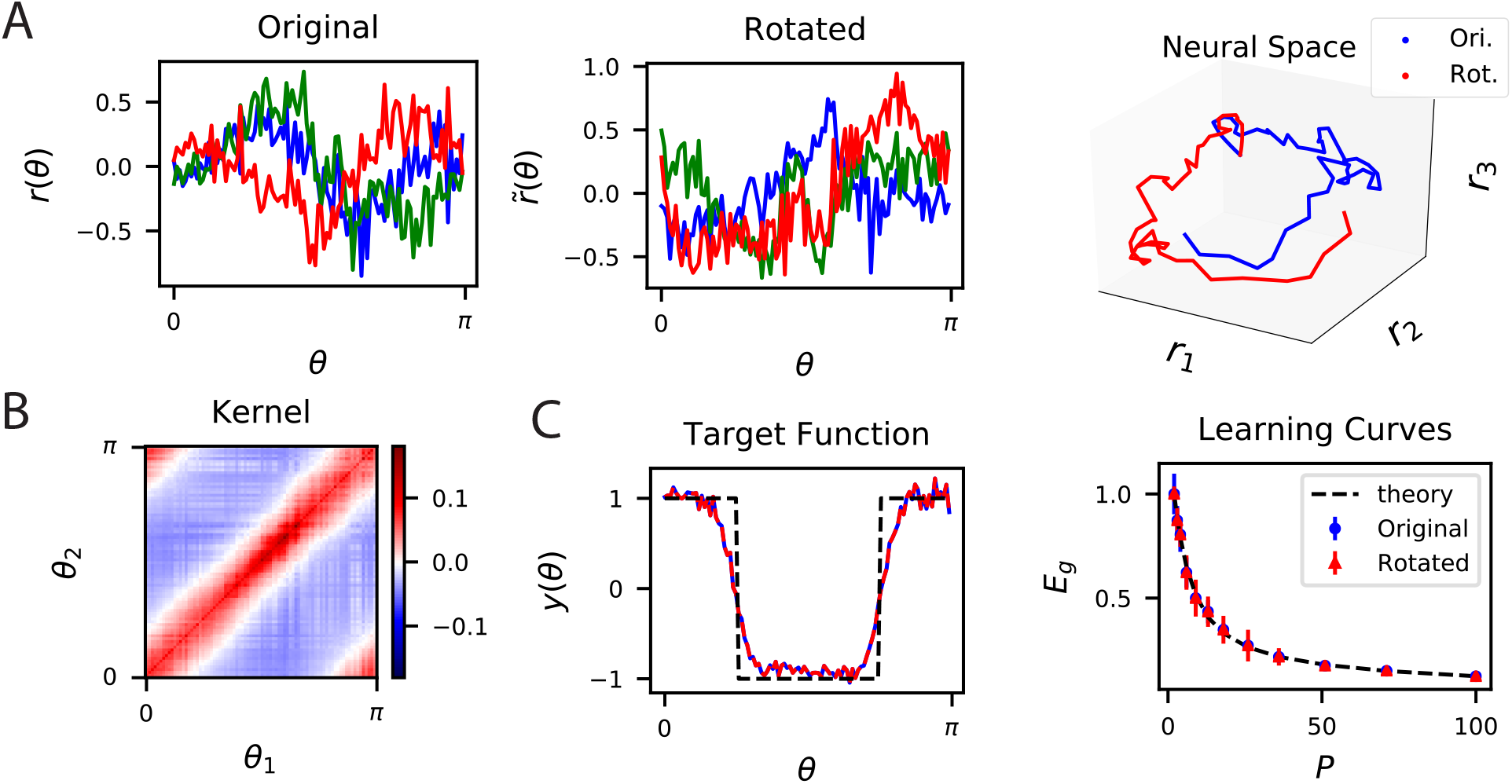
The inner product kernel controls the generalization performance of readouts. **A** Tuning curves *r*(*θ*) for three example recorded Mouse V1 neurons to varying static grating stimuli oriented at angle *θ* [8, 9] (Left) are compared with a randomly rotated version (Middle) 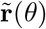 of the same population code. (Right) These two codes, original (Ori.) and rotated (Rot.) can be visualized as parametric trajectories in neural space. **B** The inner product kernel matrix has elements *K*(*θ*_1_, *θ*_2_). The original V1 code and its rotated counterpart have identical kernels. **C** In a learning task involving uniformly sampled angles, readouts from the two codes perform identically, resulting in identical approximations of the target function (shown on the left as blue and red curves) and consequently identical generalization performance as a function of training set size P (shown on right with blue and red points). The theory curve will be described in the main text.

Although, the performance of linear readouts may be invariant to such rotations, metabolic efficiency may favor certain codes over others [14, 15, 16, 17, 18], reducing degeneracy in the space of codes with identical kernels. To formalize this idea, we define ***δ*** to be the vector of spontaneous firing rates of a population of neurons, and **s**^*μ*^ = **r**(***θ**^μ^*) + ***δ*** be the spiking rate vector in response to a stimulus ***θ**^μ^*. The modulation with respect to the spontaneous activity, **r**(***θ***^*μ*^), gives the population code and defines the kernel, 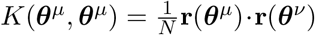. To avoid confusion with **r**(***θ***^*μ*^), we will refer to **s**^*μ*^ as total spiking activity. We propose that population codes prefer smaller spiking activity subject to a fixed kernel. In other words, because the kernel is invariant to any change of the spontaneous firing rates and left rotations of **r**(***θ***), the orientation and shift of the population code **r**(***θ***) should be chosen such that the resulting total spike count 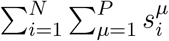 is small.

We tested whether biological codes exhibit lower total spiking activity than others exhibiting the same kernel on mouse V1 recordings, using deconvolved calcium activity as a proxy for spiking events [8, 9, 19] (Methods 4.4; Figure 3). To compare the experimental total spiking activity to other codes with identical kernels, we computed random rotations of the neural responses around spontaneous activity, 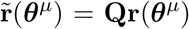, and added the 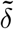 that minimizes total spiking activity and maintains its nonnegativity (Methods 4.4.2). We refer to such an operation as RROS (random rotation and optimal shift), and a code generated by an RROS operation as an RROS code The matrix **Q** is constructed by performing QR decomposition of a random Gaussian matrix, which guarantees sampling from the Haar measure of *O*(*N*) [20]. In other words, we compare the true code to the most metabolically efficient realizations of its random rotations. This procedure may result in an increased or decreased total spike count in the code, and is illustrated in a synthetic dataset in Figure 3A. We conducted this procedure on subsets of various sizes of mouse V1 neuron populations, as our proposal should hold for any subset of neurons (Methods 4.4.2), and found that the true V1 code is much more metabolically efficient than randomly rotated versions of the code (Figure 3B and C). This finding holds for both responses to static gratings and to natural images as we show in Figure 3B and C respectively.

**Figure 3:**
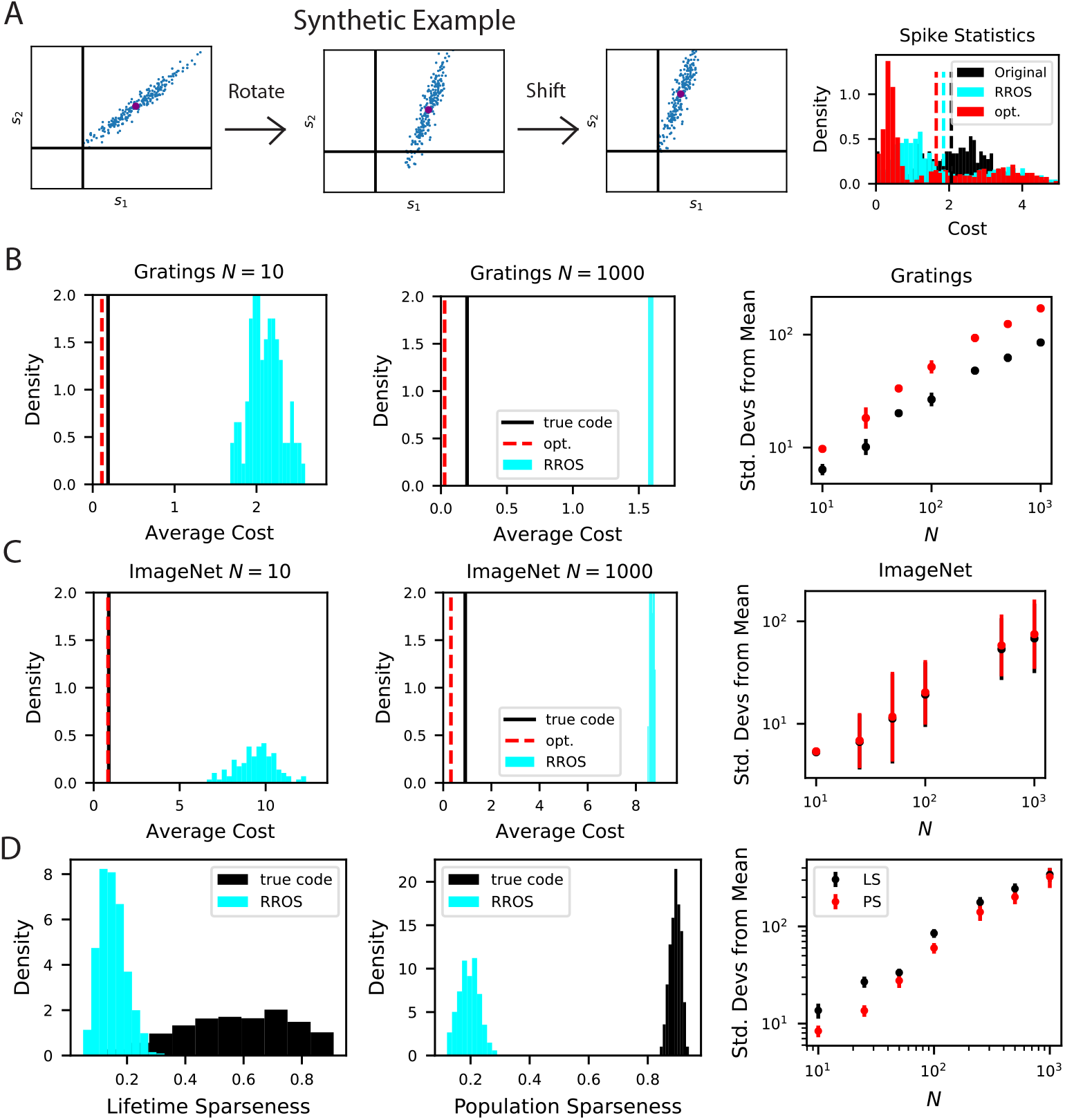
The biological code is more metabolically efficient than random codes with same inductive biases. **A** We illustrate our procedure in a synthetic example. A non-negative population code (left) can be randomly rotated about its spontaneous firing rate (middle), illustrated as a purple dot, and optimally shifted to a new non-negative population code (right). If the kernel is measured about the spontaneous firing rate, these transformations leave the inductive bias of the code invariant but can change the total spiking activity of the neural responses. We refer to such an operation as random rotation + optimal shift (RROS). We also perform gradient descent over rotations and shifts, generating an optimized code (opt). **B** Performing RROS on *N* neuron subsamples of experimental Mouse V1 recordings [8, 9], shows that the true code has much lower average cost 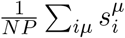 compared to random rotations of the code. The set of possible RROS transformations (Methods 4.4.2,4.4.3) generates a distribution over average cost, which has higher mean than the true code. We also optimize metabolic cost over the space of RROS transformations, which resulted in the red dashed lines. We plot the distance (in units of standard deviations) between the cost of the true and optimal codes and the cost of randomly rotated codes for different neuron subsample sizes *N*. **C** The same experiment performed on Mouse V1 responses to ImageNet images from 10 relevant classes [11, 10]. **D** The *lifetime* (LS) and *population sparseness* (PS) levels (Methods 4.4.4) are higher for the Mouse V1 code than for a RROS code. The distance between average LS and PS of true code and RROS codes increases with *N*.

To further explore metabolic efficiency, we posed an optimization problem which identifies the most efficient code with the same kernel as the biological V1 code. This problem searches over rotation matrices **Q** and finds the **Q** matrix and off-set vector ***δ*** which gives the lowest cost 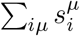 (Methods 4.4.3)(Figure 3). Though the local optimum identified with the algorithm is lower in cost than the biological code, both the optimal and biological codes are significantly displaced from the distribution of random codes with same kernel. Our findings do not change when data is preprocessed with an alternative strategy, an upper bound on neural responses is imposed on rotated codes, or subsets of stimuli are considered (Figure App.1). We further verified these results on electrophysiological recordings of mouse visual cortex from the Allen Institute Brain Observatory [21], (Figure App.2). Overall, the large disparity in total spiking activity between the true and randomly generated codes with identical kernels suggests that metabolic constraints may favor the biological code over others that realize the same kernel.

The disparity between the true biological code and the RROS code is not only manifested in terms of total activity level, but also in terms of single neuron and single stimulus sparseness measures, specifically lifetime and population sparseness distributions (Methods 4.4.4) [22, 23, 24, 25]. In Figure 3D, we compare the lifetime and population sparseness distributions of the true biological code with a RROS version of the same code, revealing biological neurons have significantly higher lifetime sparseness. In App. Section 2B, we provide analytical arguments which suggest that tuning curves of optimally sparse non-negative codes with full-rank kernels will have selective tuning.

### Code-task alignment governs generalization

We next examine how the population code affects generalization performance of the readout. We calculated analytical expressions of the average generalization error in a task defined by the target response *y*(***θ***) after observing *P* stimuli using methods from statistical physics (Methods 4.2). Because the relevant quantity in learning performance is the kernel, we leveraged results from our previous work studying generalization in kernel regression [26, 27], and approximated the generalization error averaged over all possible realizations of the training dataset of composed of *P* stimuli, 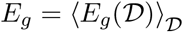. As *P* increases, the variance in *E_g_* due to the composition of the dataset falls, and our expressions become descriptive of also the typical case. Our final analytical result is given in Equation (11) in Methods 4.2. We provide details of our calculations in Methods 4.2 and App. Section 3, and focus on their implications here.

One of our main observations is that given a population code **r**(***θ***), the singular value decomposition of the code gives the appropriate basis to analyze the inductive biases of the readouts (Figure 4A). The tuning curves for individual neurons *r_i_*(***θ***) form an *N*-by-*M* matrix **R**, where *M*, possibly infinite, is the number of all possible stimuli. We discuss the SVD for continuous stimulus spaces in App. Section 1A. The left-singular vectors (or principal axes) and singular values of this matrix have been used in neuroscience for describing lower dimensional structure in the neural activity and estimating its dimensionality, see e.g. [28, 29, 30, 31, 32, 33, 11, 8, 34, 35, 36]. We found that the function approximation properties of the code are controlled by the singular values, or rather their squares {λ_*k*_} which give variances along principal axes, indexed in decreasing order, and the corresponding right singular vectors {*ψ_k_* (***θ***)}, which are also the kernel eigenfunctions (Methods 4.2 and App. Section 1). This follows from the fact that learned response (Eq. (2)) is only a function of the kernel *K*, and the eigenvalues λ_*k*_ and orthonormal eigenfunctions *ψ_k_*(***θ***) collectively define the code’s inner-product kernel *K*(**θ, θ**′) through an eigendecomposition 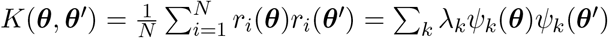 [37] (Methods 4.2 and App. 3).

**Figure 4:**
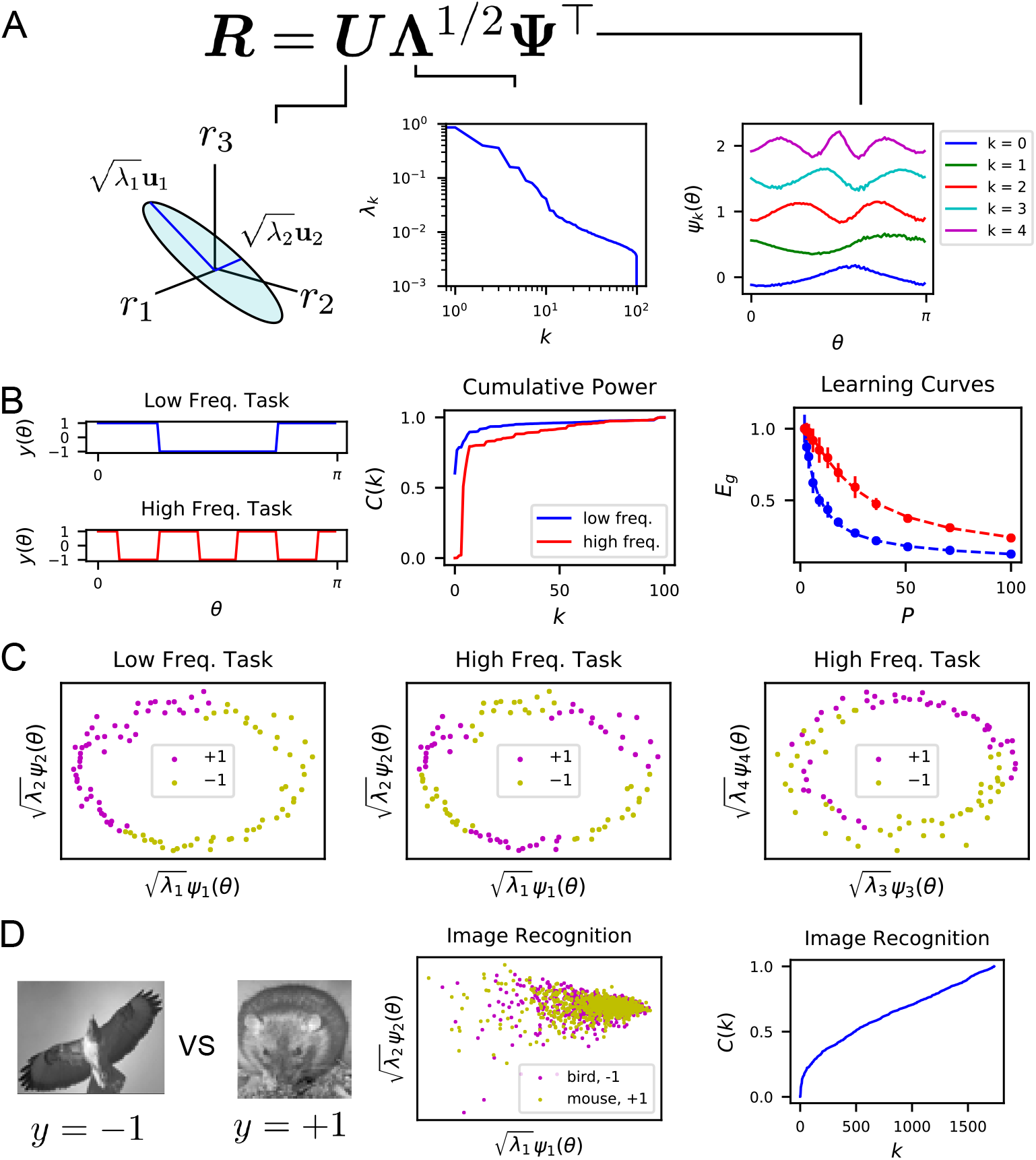
The singular value decomposition of the population code reveals the structure and inductive bias of the code. **A** Singular value decomposition of the response matrix **R** gives left singular vectors **u**_*k*_ (principal axes), kernel eigenvalues *λ_k_*, and kernel eigenfunctions *ψ_k_*(***θ***). The ordering of eigenvalues provides an ordering of which modes *ψ_k_* can be learned by the code from few training examples. The eigenfunctions were offset by 0.5 for visibility. **B** (Left) Two different learning tasks *y*(*θ*), a low frequency (blue) and high frequency (red) function, are shown. (Middle) The cumulative power distribution rises more rapidly for the low frequency task than the high frequency, indicating better alignment with top kernel eigenfunctions and consequently more sample-efficient learning as shown in the learning curves (right). Dashed lines show theoretical generalization error while dots and solid vertical lines are experimental average and standard deviation over 30 repeats. **C** The feature space representations of the low (left) and high (middle and right) frequeny tasks. Each point represents the embedding of a stimulus response vector along the *k*-th principal axis 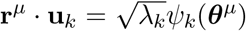. The binary target value {±1} is indicated with the color of the point. The easy (left), low frequency task is well separated along the top two dimensions, while the hard, high frequency task is not linearly separable in two (middle) or even with four feature dimensions (right). **D** On an image discrimination task (recognizing birds vs mice), V1 has an entangled representation which does not allow good performance of linear readouts. This is evidenced by the top principal components (middle) and the slowly rising *C*(*k*) curve (right).

Our analysis shows the existence of a bias in the readout towards learning certain target responses faster than others. The kernel eigenfunctions form a complete basis for square integrable functions, allowing the expansion of the target response *y*(***θ***) ∑_*k*_ *v_k_ψ_k_*(***θ***) and the learned readout response 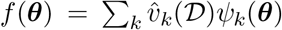 in this basis. We found that the readout’s generalization is better if the target function *y*(***θ***) is aligned with the top eigenfunctions *ψ_k_*, equivalent to 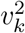 decaying rapidly with *k* (App. Section 3B,C). We formalize this notion by the following metric. Mathematically, generalization error 〈*E_g_*〉 can be decomposed into normalized estimation errors *E_k_* for the coefficients of these eigenfunctions 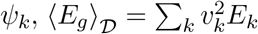, where 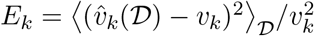. We found that the ordering of the eigenvalues λ_*k*_ controls the rates at which these mode errors *E_k_* decrease as *P* increases (App. Section 3C, [26]): λ_*k*_ > λ_*ℓ*_ ⇒ *E_k_* < *E_ℓ_*. Hence, larger eigenvalues mean lower generalization error for those normalized mode errors *E*k. We term this phenomenon the *spectral bias* of the readout. Based on this observation, we propose *code-task alignment* as a principle for good generalization. To quantify code-task alignment, we use a metric which was introduced in [27] to measure the compatibility of a kernel with a learning task. This is the cumulative power distribution *C*(*k*) which measures the total power in of the target function in the top *k* eigenmodes, normalized by the total power [27]:

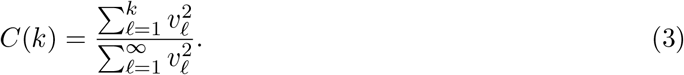

Stimulus-response maps that have high alignment with the population code’s kernel will have quickly rising cumulative power distributions *C*(*k*), since a large proportion of power is placed in the top modes. Target responses with high *C*(*k*) can be learned with fewer training samples than target responses with low *C*(*k*) since the mode errors *E_k_* are ordered for all *P* (App. Section 3C).

### Probing learning biases in neural data

Our theory can be used to probe the learning biases of neural populations. Here we provide various examples of this using publicly available calcium imaging recordings from mouse primary visual cortex (V1). Our examples illustrate how our approach can be used to analyze neural population data.

We first analyzed population responses to static grating stimuli oriented at an angle *θ* [8, 9]. We found that the kernel eigenfunctions have sinusoidal shape with differing frequency. The ordering of the eigenvalues and eigenfunctions in Figure 4A indicates a frequency bias: lower frequency functions of *θ* are easier to estimate at small sample sizes.

We tested this idea by constructing two different orientation discrimination tasks shown in Figures 4B,C, where we assign static grating orientations to positive or negative valence with different frequency square wave functions of *θ*. We trained the readout using a subset of the experimentally measured neural responses, and measured the readout’s generalization performance. We found that the cumulative power distribution for the low frequency task has a more rapidly rising *C*(*k*) (Figure 4B). Using our theory of generalization, we predicted learning curves for these two tasks, which express the generalization error as a function of the number of sampled stimuli *P*. The error for the low frequency task is lower at all sample sizes than the hard task. The theoretical predictions and numerical experiments show perfect agreement (Figure 4B). More intuition can be gained by visualizing by projection of the neural response along the top principal axes (Figure 4C). For the low frequency task, the two target values are well separated along the top two axes. However, the high frequency task is not well separated along even the top four axes (Figure 4C).

Using the same ideas, we can use our theory to get insight into tasks which the V1 population code is ill-suited to learn. For the task of identifying mice and birds [11, 10] the linear rise in cumulative power indicates that there is roughly equal power along all kernel eigenfunctions, indicative of a representation poorly aligned to this task (Figure 4D).

To illustrate how our approach can be used for different learning problems, we evaluate the ability of linear readouts to reconstruct natural images from neural responses to those images (Figure 5). The ability to reconstruct sensory stimuli from a neural code is an influential normative principle for primary visual cortex [17]. Here we ask which aspects of the presented natural scene stimuli are easiest to learn to reconstruct. Since mouse V1 neurons tend to be selective towards low spatial frequency bands [38, 39, 40], we consider reconstruction of band-pass filtered images with spatial frequency constrained to 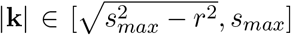 for *r* = 0.2 (in units of pixels^-1^) and plot the cumulative power *C*(*k*) associated with each choice of the upper limit *s_max_* (Figure 5C,D). The frequency cutoffs were chosen in this way to preserve the volume in Fourier space to *V_k_* = *πr*^2^, which quantifies the dimension of the function space. We see that the lower frequency band-limited images are easier to reconstruct, as evidenced by their cumulative power *C*(*k*) and learning curves *E_g_* (Figure 5D,E). This reflects the fact that the population preferentially encodes low spatial frequency content in the image (Figure 5F). Experiments with additional values of r are provided in the Figure App.2 with additional details found in the App. Section 4.

**Figure 5:**
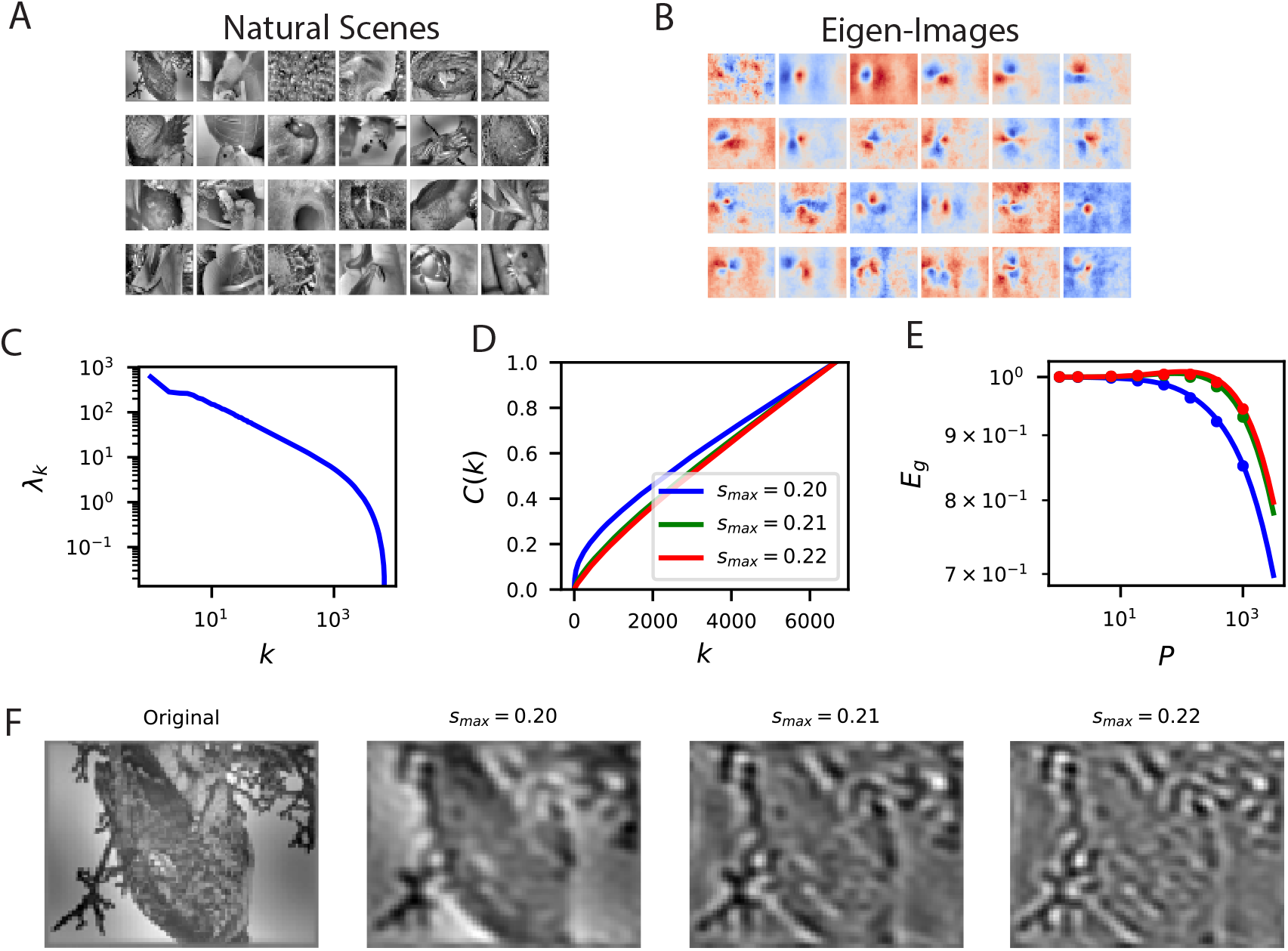
Reconstructing filtered natural images from V1 responses reveals preference for low spatial frequencies. **A** Natural scene stimuli ***θ*** were presented to mice and V1 cells were recorded. **B** The images weighted by the top eigenfunctions **v**_*k*_ = 〈*ϕ_k_*(***θ***)***θ***〉_***θ***_. These “eigenimages” collectively define the difficulty of reconstructing images through readout. **C** The kernel spectrum of the V1 code for natural images. **D** The cumulative power curves for reconstruction of band-pass filtered images. Filters preserve spatial frequencies in the range 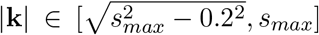, chosen to preserve volume in Fourier space as *s_max_* is varied. **E** The learning curves obey the ordering of the cumulative power curves. The images filtered with the lowest band-pass cutoff are easiest to reconstruct from the neural responses. **F** Examples of a band-pass filtered image with different preserved frequency bands.

### Neural mechanisms of low frequency bias and code-task alignment in a simple model of V1

How do population level inductive biases arise from properties of single neurons? To illustrate that a variety of mechanisms may be involved in a complex manner, we study a simple model of V1 to elucidate neural mechanisms that lead to the low frequency bias at the population level. In particular, we focus on neural nonlinearities and selectivity profiles.

We model responses of V1 neurons as photoreceptor inputs passed through Gabor filters and a subsequent nonlinearity, *g*(*z*), modeling a population of orientation selective simple cells (Figure 6A) (see App. 5). In this model, the kernel for static gratings with orientation *θ* ∈ [0, *π*] is of the form *K*(*θ, θ*′) = *κ*(|*θ* – *θ*′|), and, as a consequence, the eigenfunctions of the kernel in this setting are Fourier modes. The eigenvalues, and hence the strength of the spectral bias, are determined by the nonlinearity.

**Figure 6:**
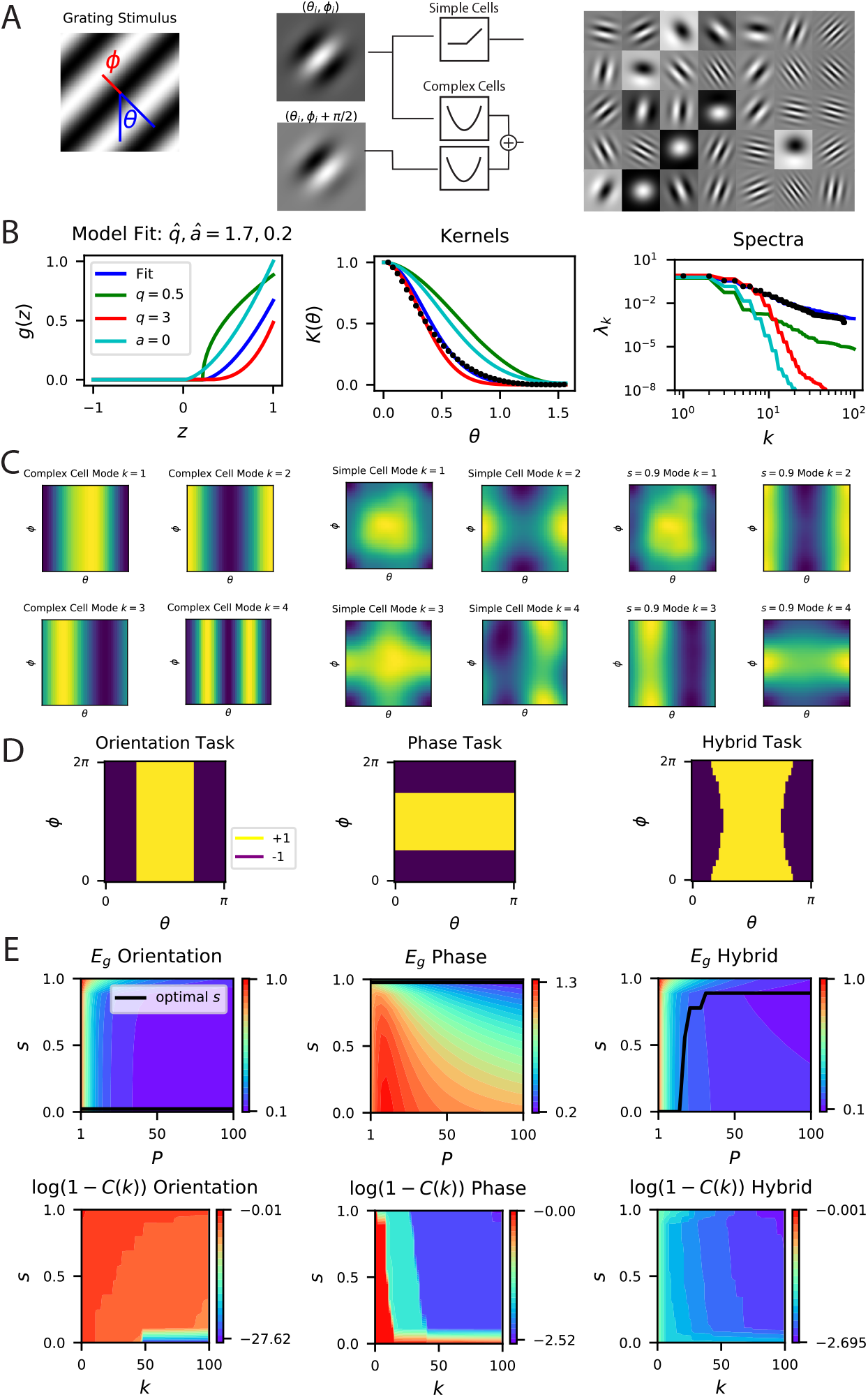
A model of V1 as a bank of Gabor filters recapitulates experimental inductive bias. **A** Gabor filtered inputs are mapped through nonlinearity. A grating stimulus (left) with orientation *θ* and phase *ϕ* is mapped through a circuit of simple and complex cells (middle). Some examples of randomly sampled Gabor filters (right) generate preferred orientation tuning of neurons in the population. **B** A threshold-powerlaw nonlinearity *g_q,a_*(*z*) = max{0,*z* – *a*}^*q*^ is fit to the mouse V1 kernel (black dots). Kernels and spectra for alternative choices of *q, a* are shown (color code defined in left panel). **C** We plot eigenfunctions *ψ_k_* (modes) for mixtures of sN simple cells and (1 – *s*)*N* complex cells. A pure complex cell population has all eigenfunctions independent of phase *ϕ*. A pure simple cell population *s* = 1 or mixture codes 0 < *s* < 1 depend on both orientation phase in a nontrivial way. **D** Three tasks are visualized, where color indicates the binary target value ±1. The left task only depends on orientation stimulus variable *θ*, the middle only depends on phase *ϕ*, the hybrid task (right) depends on both. **E** (top) Generalization error and cumulative power distributions for the three tasks as a function of the simple-complex cell mixture parameter *s*. In Figure App.3 we provide more comparisons of our theory and numerical experiments.

Motivated by findings in the primary visual cortex [41, 42, 43, 44], we studied the spectral bias induced by rectified power-law nonlinearities of the form *g*(*z*) = max{0, *z* – *a*}^*q*^. From theory side, such a power-law activation function arises in a spiking neuron when firing is driven by input fluctuations [41, 42]. Further, this activation is observed in intracellular recordings over the dynamic range of neurons in primary visual cortex [44]. For example, in cats, the power, *q,* ranges from 2.7 to 3.9 [43]. We fit parameters of our model to the Mouse V1 kernel and compared to other parameter sets in Figure 6B. Our best fit value of *q* = 1.7 is lower but on par with the estimates from the cat.

Computation of the kernel and its eigenvalues (App. 5A for linear *g*, App. 5B for threshold-power law *g*) indicates a low frequency bias: the eigenvalues for low frequency modes are higher than those for high frequency modes, indicating a strong inductive bias to learn functions of low frequency in the orientation. Decreasing sparsity (lower *a*) leads to a faster decrease in the spectrum (but similar asymptotic scaling at the tail, see App. Section 5) and a stronger bias towards lower frequency functions (Figure 6B; more comparisons in Figure App.3). The effect of the power of nonlinearity *q* is more nuanced: increasing power may increase spectra at lower frequencies, but may also lead to a faster decay at the tail (Figure 6B; more comparisons in Figure App.3). In general, an exponent *q* implies a power-law asymptotic spectral decay λ_*k*_ ~ *k*^-2*q*-2^ as *k* → ∞ (App. Section 5B). The behavior at low frequencies may have significant impact for learning with few samples. We discuss this in more detail in the next section. Overall, our findings show that the spectral bias of a population code can be determined in non-trivial ways by its biophysical parameters, including neural thresholds and nonlinearities.

To further illustrate the importance of code-task alignment, we next study how invariances in the code to stimulus variations may affect the learning performance. We introduce complex cells in addition to simple cells in our model with proportion *s* ∈ [0,1] of simple cells (App. Section 5C; Figure 6A), and allow phase, *ϕ,* variations in static gratings. We use the energy model [45, 46] to capture the phase invariant complex cell responses (App. 5B, 5C). We reason that in tasks that do not depend on phase information, complex cells should improve sample efficiency.

In this model, the kernel for the V1 population is a convex combination of the kernels for the simple and complex cell populations *K*_*V*1_(*θ, θ*′, *ϕ, ϕ*′) = *sK_s_*(*θ, θ*′, *ϕ, ϕ*′) + (1 – *s*)*K_c_*(*θ, θ*′) where *K_s_* is the kernel for a pure simple cell population that depends on both orientation and phase, and *K_c_* is the kernel of a pure complex cell population that is invariant to phase (App. 5C). Figure 6C shows top kernel eigenfunctions for various values of *s* elucidating inductive bias of the readout.

Figures 6D and 6E show generalization performance on tasks with varying levels of dependence on phase and orientation. On pure orientation discrimination tasks, increasing the proportion of complex cells by decreasing *s* improves generalization. Increasing the sensitivity to the nuisance phase variable, *ϕ*, only degrades performance. The cumulative power distribution is also maximized at *s* = 0. However, on a task which only depends on the phase, a pure complex cell population cannot generalize, since variation in the target function due to changes in phase cannot be explained in the codes’ responses. In this setting, a pure simple cell population attains optimal performance. The cumulative power distribution is maximized at *s* = 1. Lastly, in a nontrivial hybrid task which requires utilization of both variables *θ, ϕ*, an optimal mixture *s* exists for each sample budget *P* which minimizes the generalization error. The cumulative power distribution is maximized at different s values depending on *k,* the component of the target function. This is consistent with an optimal heterogenous mix, because components of the target are learned successively with increasing sample size. In reality, V1 must code for a variety of possible tasks and we can expect a nontrivial optimal simple cell fraction *s*. We conclude that the degree of invariance required for the set of natural tasks, and the number of samples determine the optimal simple cell, complex cell mix.

### Small and large sample size behaviors of generalization

Recently, Stringer *et al.* [11] argued that the input-output differentiability of the code may be enabling better generalization, which is in turn governed by the asymptotic rate of spectral decay. Our results provide a more nuanced view of the relation between generalization and kernel spectra. First, generalization with low sample sizes crucially depend on the top eigenvalues and eigenfunctions of the code’s kernel, not the tail. Second, generalization requires alignment of the code with the task of interest. Non-differentiable codes can generalize well if there is such an alignment. To illustrate these points, here, we provide examples where asymptotic conditions on the kernel spectrum are insufficient to describe generalization performance for small sample sizes (Figure 7 and App.5), and where non-differentiable kernels generalize better than differentiable kernels (Figure App.6).

**Figure 7:**
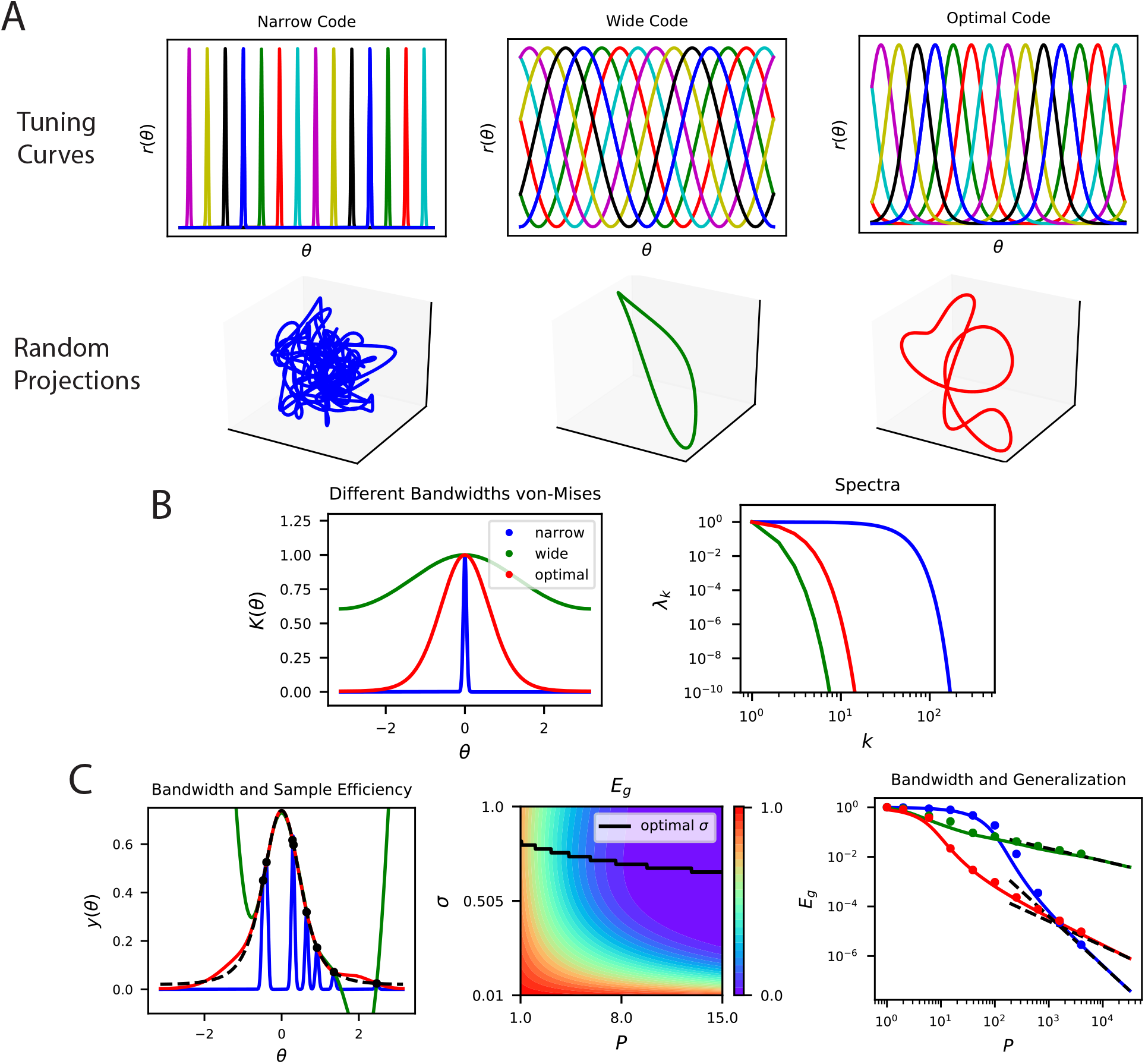
The top eigensystem of a code determines its low-*P* generalization error. **A** A periodic variable is coded by a population of neurons with tuning curves of different widths (top). Narrow, wide and optimal refers to the example in C. These codes are all smooth (infinitely differentiable) but have very different feature space representations of the stimulus variable *θ*, as random projections reveal (below). **B** (left) The population codes in the above figure induce von Mises kernels *K*(*θ*) ∝ *e*^cos(*θ*)/*σ*2^ with different bandwidths *σ*. (right) Eigenvalues of the three kernels. **C** (left) As an example learning task, we consider estimating a “bump” target function. The optimal kernel (red, chosen as optimal bandwidth for *P* = 10) achieves a better generalization error than either the wide (green) or narrow (blue) kernels. (middle) A contour plot shows generalization error for varying bandwidth *σ* and sample size *P*. (right) The large *P* generalization error scales in a power law. Solid lines are theory, dots are simulations averaged over 15 repeats, dashed lines are asymptotic power law scalings described in main text. Same color code as B and C-left.

Our example demonstrates how a code allowing good generalization for large sample sizes can be disadvantageous for small sizes. In Figure 7A, we plot three different populations of neurons with smooth (infinitely differentiable) tuning curves that tile a periodic stimulus variable, such as the direction of a moving grating. The tuning width, *σ*, of the tuning curves strongly influences the structure of these codes: narrower widths have more high frequency content as we illustrate in a random 3D projection of the population code for *θ* ∈ [0,2*π*] (Figure 7A). Visualization of the corresponding (von Mises) kernels and their spectra are provided in Figure 7B. The width of the tuning curves control bandwidths of the kernel spectra Figure 7B, with narrower curves having an later decay in the spectrum and higher high frequency eigenvalues. These codes can have dramatically different generalization performance, which we illustrate with a simple “bump” target response (Figure 7C). In this example, for illustration purposes, we let the network learn with a delta-rule with a weight decay, leading to a regularized kernel regression solution (App. 3A.1). For a sample size of *P* = 10, we observe that codes with too wide or too narrow tuning curves (and kernels) do not perform well, and there is a well-performing code with an optimal tuning curve width *σ*, which is compatible with the width of the target bump, *σ_T_*. We found that optimal *σ* is different for each *P* (Figure 7C). In the large-P regime, the ordering of the performance of the three codes are reversed (Figure 7C). In this regime generalization error scales in a power law 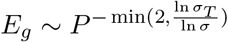 (App. 3D) and the narrow code, which performed worst for *P* ~ 10, performs the best. This example demonstrates that asymptotic conditions on the tail of the spectra are insufficient to understand generalization in the small sample size limit. The bulk of the kernel’s spectrum needs to match the spectral structure of the task to generalize efficiently in the low-sample size regime. However, for large sample sizes, the tail of the eigenvalue spectrum becomes important. We repeat the same exercise and draw the same conclusions for a non-differentiable kernel (Laplace) (Figure App.5) showing that these results are not an artifact of the infinite differentiability of von Mises kernels. We further provide examples where non-differentiable kernels generalizing better than differentiable kernels in Figure App.5.

### Time-Dependent Neural Codes

Our framework can directly be extended to learning of arbitrary time-varying functions of time-varying inputs from an arbitrary spatiotemporal population code (Methods 4.3, App. 6). In this setting, the population code **r**({***θ***(*t*)}, *t*) is a function of an input stimulus sequence ***θ***(*t*) and possibly its entire history, and time *t*. A downstream linear readout *f*({***θ***},*t*) = **w** · **r**({***θ***},*t*) learns a target sequence *y*({***θ***},*t*) from a total of 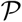 examples that can come at any time during any sequence. Learning is again achieved through the delta-rule and the learned function can be expressed as a linear combination of the kernel evaluated at the 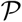 examples. The kernel in this case is a more complicated object that computes inner products of neural population vectors at different times *t,t′* for different input sequences {***θ***}, 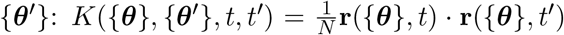 [47, 48, 49]. Our theory carries over from the static case with appropriate modifications (Methods 4.3). Kernels whose top eigenfunctions have high alignment with the target time-varying response *y*({***θ***}, *t*) will achieve the best average case generalization performance.

As a concrete example, we focus on readout from a temporal population code generated by a recurrent neural network in a task motivated by a delayed reach task [50] (Figure 8A,B). We consider a randomly connected network with current dynamics 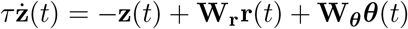 where the rates are related to input currents through a tanh nonlinearity **r**(*t*) = tanh(**z**(*t*)). The recurrent weights are drawn from a normal distribution 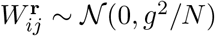 and the input encoding weights from 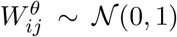 (Methods 4.3). The gain parameter *g* was set to 1.5 to generate rich dynamics [51]. In this task, the network is presented for a short time an input cue sequence coding an angular variable which is drawn randomly from a distribution (Figure 8C). The recurrent neural network must remember this angle and reproduce an output sequence which is a simple step function whose height depends on the angle which begins after a time delay from the cessation of input stimulus and lasts for a short time (Figure 8D).

**Figure 8:**
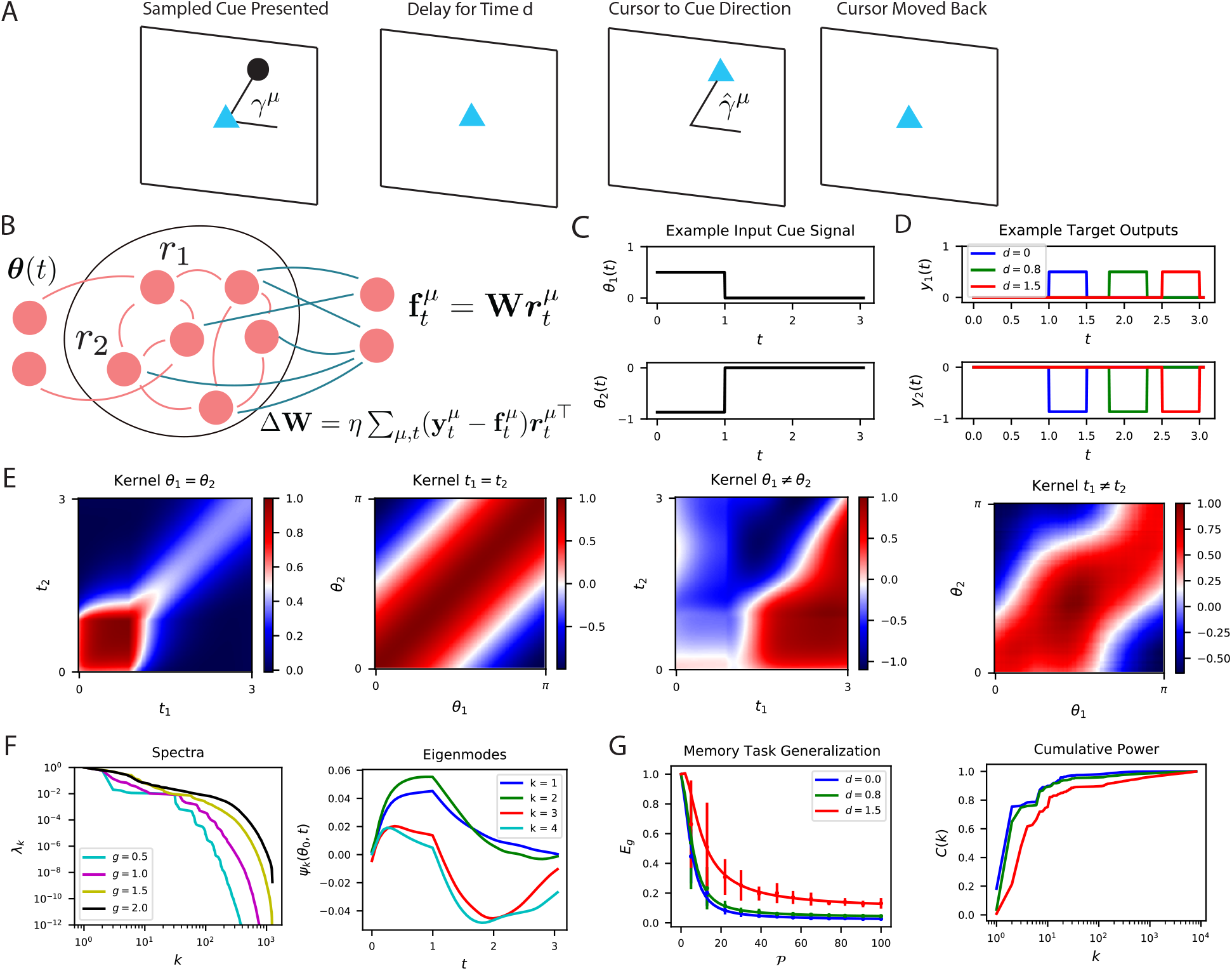
The performance of time-dependent codes when learning dynamical systems can be understood through spectral bias. **A** We study the performance of time dependent codes on a delayed response task which requires memory retrieval. A cue (black dot) is presented at an angle *γ^μ^*. After a delay time *d*, the cursor position (blue triangle) must be moved to the remembered cue position and then subsequently moved back to the origin after a short time. **B** The readout weights (blue) of a time dependent code can be learned through a modified delta rule. **C** Input is presented to the network as a time series which terminates at *t* = 1. The sequences are generated by drawing an angle *γ^μ^* ~ Uniform[0, 2*π*] and using two step functions as input time-series that code for the cosine and the sine of the angle (Methods 4.3, App. 6). We show an example of the one of the variables in a input sequence. **D** The target functions for the memory retrieval task are step functions delayed by a time *d*. **E** The kernel *K_μ,μ′,t,t′_* compares the code for two sequences at two distinct time points. We show the time dependent kernel for identical sequences (left) and the stimulus dependent kernel for equal time points (middle left) as well as for non-equal stimuli (middle right) and non-equal time (right). **F** The kernel can be diagonalized, and the eigenvalues λ_*k*_ determine the spectral bias of the reservoir computer (left). We see that higher gain *g* networks have higher dimensional representations. The “eigensystems” *ψ_k_* (*θ^μ^, t*) are functions of time and cue angle. We plot only *μ* = 0 components of top systems *k* = 1, 2, 3, 4 (right). **G** The readout is trained to approximate a target function *y^μ^*(*t*), which requires memory of the presented cue angle. (left) The theoretical (solid) and experimental (vertical errorbar, 100 trials) generalization error *E_g_* are plotted for the three delays d against training sample size 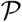. (right) The ordering of *E_g_* matches the ordering of the *C*(*k*) curves as expected.

The kernel induced by the spatiotemporal code is shown in Figure 8E. The high dimensional nature of the activity in the recurrent network introduces complex and rich spatiotemporal similarity structure. Figure 8F shows the kernel’s eigensystem, which consists of stimulus dependent time-series *ψ_k_*({***θ***}; *t*) for each eigenvalue *λ_k_*. An interesting link can be made with this eigensystem and linear low-dimensional manifold dynamics observed in several cortical areas [28, 29, 31, 52, 33, 53, 36, 32, 54, 30]. The kernel eigenfunctions also define the latent variables obtained through a singular value decomposition of the neural activity 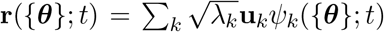 [31].

With enough samples, the readout neuron can learn to output the desired angle with high fidelity (Figure 8G). Unsurprisingly, tasks involving long time delays are more difficult and exhibit lower cumulative power curves. Consequently, the generalization error for small delay tasks drops much more quickly with increasing 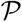.

## 3 Discussion

Elucidating inductive biases of the brain is fundamentally important for understanding natural intelligence [3, 4, 5, 55]. These biases are coded into the brain by the dynamics of its neurons, the architecture of its networks, its representations and plasticity rules. Finding ways to extract the inductive biases from neuroscience datasets requires a deeper theoretical understanding of how all these factors shape the biases, and is an open problem. In this work, we attempted to take a step towards filling this gap by focusing on how the structure of static neural population codes shape inductive biases for learning of a linear readout neuron under a biologically plausible learning rule. If the readout neuron’s output is correlated with behavior, and that correlation is known, then our theory could possibly be modified to predict what behavioral tasks can be learned faster.

Under the delta rule, the generalization performance of the readout is entirely dependent on the code’s inner product kernel; the kernel is a determinant of inductive bias. In its finite dimensional form, the kernel is an example of a representational similarity matrix and is a commonly used tool to study neural representations [56, 57, 58, 59, 60, 61]. Our work elucidates a concrete link between this experimentally measurable mathematical object, and sample-efficient learning.

We derived an analytical expression for the generalization error as a function of sample-size under very general conditions, for an arbitrary stimulus distribution, arbitrary population code and an arbitrary target stimulus-response map. We used our findings in both theoretical and experimental analysis of primary visual cortex, and temporal codes in a delayed reach task. This generality of our theory is a particular strength.

Our analysis elucidated two principles that define the inductive bias. The first one is spectral bias: kernel eigenfunctions with large eigenvalues can be estimated using a smaller number of samples. The second principle is the code-task alignment: Target functions with most of their power in top kernel eigenfunctions can be estimated efficiently and are compatible with a code. The cumulative power distribution, *C*(*k*) [27], provides a measure of this alignment. These findings define a notion of “simplicity” bias in learning from examples, and provides a solution to the question of what stimulus-response maps are easier to learn. A similar simplicity bias has been also observed in training deep neural networks [62, 63, 64]. Due to a correspondence between trained neural networks in the infinite-width limit and kernel machines [65], results on the spectral bias of kernel machines may shed light onto these findings [26, 27].

We used our finding in both theoretical and experimental analysis of mouse primary visual cortex. We demonstrated a bias of neural populations towards low frequency orientation discrimination and low spatial frequency reconstruction tasks. The latter finding is consistent with the finding that mouse visual cortex neurons are selective for low spatial frequency [38, 39, 40]. Our modeling showed that these biases can be modified by the neural thresholds and nonlinearities in nontrivial ways. The theoretical model of the visual cortex as a mixture of simple and complex cells demonstrated how invariances, specifically the phase invariance of the complex cells, in the population code can facilitate learning some tasks involving phase invariant responses at the expense of performance on others. Role of invariances in learning with kernel methods have recently been investigated in machine learning literature [66, 67].

A recent proposal considered the possibility that the brain acts as an overparameterized interpolator [68]. Suitable inductive biases are crucial to prevent overfitting and generalize well in such a regime [69]. Our theory can explain these inductive biases since, when the kernel is full-rank, which typically is the case when there are more neurons in the population than the number of learning examples, the delta rule converges to an interpolator of the learning examples. Modern deep learning architectures also operate in an overparameterized regime, but generalize well [70, 69], and an inductive bias towards simple functions has been proposed as an explanation [26, 27, 64, 71].

Our work suggests sample efficiency as a general coding principle for neural populations, relating neural representations to the kinds of problems they are well suited to solve. These codes may be shaped through evolution or themselves be learned through experience [55]. Prior related work demonstrated the dependence of sample-efficient learning of a two-angle estimation task on the width of the individual neural tuning curves [72] and additive function approximation properties of sparsely connected random networks [73].

A sample efficiency approach to population coding differs from the classical efficient coding theories [16, 14, 15, 74, 75, 76, 17, 77], which postulate that populations of neurons optimize information content of their code subject to metabolic constraints or noise. While these theories emphasize different aspect of the code’s information content (such as reduced redundancy, predictive power, or sparsity), they do not address sample efficiency demands on learning. Further, recent studies demonstrated hallmarks of redundancy and correlation in population responses [54, 30, 78, 36, 79, 80, 11], violating a generic prediction of efficient coding theories that responses of different neurons should be uncorrelated across input stimuli in high signal-to-noise regimes to reduce redundancy in the code and maximize information content [14, 15, 74, 75, 81, 82]. In our theory, the structured correlations of neural responses correspond to the decay in the spectrum of the kernel, and play a key role in biasing learned readouts towards simple functions.

In recent related studies, the asymptotic decay rate of the kernel’s eigenspectrum was argued to be important for generalization [11] and robustness [83]. Decay rate in the mouse V1 was found to be consistent with a high dimensional (power law) but smooth (differentiable) code, and smoothness was argued to be an enabler of generalization [11]. We show that sample-efficient learning requires more than smoothness conditions in the form of asymptotic decay rates on the kernel’s spectrum. The interplay between the stimulus distribution, target response and the code gives rise to sample efficient learning. Because of spectral bias, the top eigenvalues govern the small sample size behavior. The tail of the spectrum becomes important for large sample sizes.

Though the kernel is degenerate with respect to rotations of the code in the neural activity space, we demonstrated that the true V1 code has much lower average activity than random codes with the same kernel, suggesting that evolution and learning may be selecting neural codes with low average spike rates which preserve sample-efficiency demands for downstream learning tasks. We predict that metabolic efficiency may be a determinant in the orientation and placement of the ubiquitously observed low-dimensional coding manifolds [53, 80] in neural activity space in other parts of the brain. The demand of metabolic efficiency is consistent with prior sparse coding theories [84, 17, 18, 85], however, our theory emphasizes sample-efficient learning as the primary normative objective for the code.

Our work constitutes a first step towards understanding inductive biases in neuronal circuits. To achieve this, we focused on a linear, delta-rule readout of a static population code. More work is need to study other factors that affect inductive bias. Importantly, sensory neuron tuning curves can adapt during perceptual learning tasks [86, 87, 88, 89] with the strength of adaptation dependent on brain area [90, 91, 92, 93]. This motivates analysis of learning in multi-layer networks. Our work focused on the effect of signal correlations to coding and inductive bias [94, 95]. Future analysis could study how signal and noise correlations interact to shape inductive bias and determine generalization. Other factors that need to be taken into account are readout nonlinearities, alternative learning rules, initialization schemes, and network architectures.

## 4 Methods

### 4.1 Generating example codes (Figure 1)

The two codes in Figure 1 were constructed to produce two different kernels for *θ* ∈ *S*^1^:

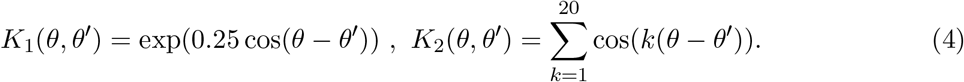

An infinite number of codes could generate either of these kernels. After diagonalizing the kernel into its eigenfunctions on a grid of 120 points, 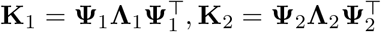, we used a random rotation matrix **Q** ∈ *O*(120) to generate a valid code

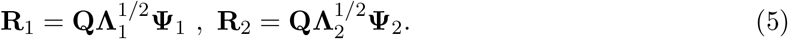

This construction guarantees that 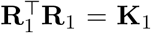 and 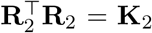. We plot the tuning curves for the first three neurons. The target function in the first experiment is *y* = cos(*θ*) – 0.6cos(4*θ*), while the second experiment used *y* = cos(6*θ*) – cos(8*θ*).

### 4.2 Theory of Generalization

Recent work has established analytic results that predict the average case generalization error for kernel regression

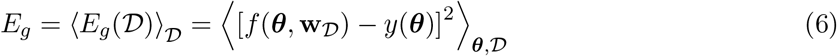

where 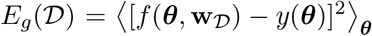 is the generalization error for a certain sample 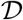 of size *P* and 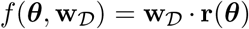 is the kernel regression solution for 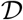 (App. 3A) [26, 27]. The typical or average case error *E_g_* is obtained by averaging over all possible datasets of size P. This average case generalization error is determined solely by the decomposition of the target function *y*(**x**) along the eigenbasis of the kernel and the eigenspectrum of the kernel. This continuous diagonalization again takes the form (App. 1A) [96]

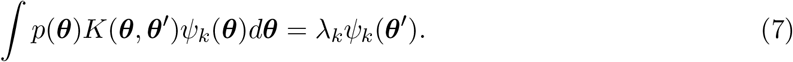

Our theory is also applicable to discrete stimuli if *p*(***θ***) is a Dirac measure as we describe in (App. 1B). Since the eigenfunctions form a complete set of square integrable functions [96], we expand both the target function *y*(***θ***) and the learned function 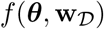 in this basis *y*(***θ***) = ∑_*k*_ *v_k_ψ_k_*(***θ***), 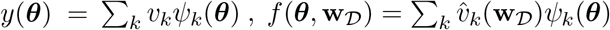. Due to the orthonormality of the kernel eigenfunctions {*ψ_k_*}, the generalization error for any set of coefficients 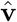 is

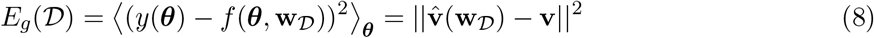

We now introduce the equivalent training error, or empirical loss, written directly in terms of eigenfunction coefficients 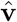, which depends on the disorder in the dataset 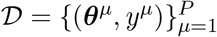

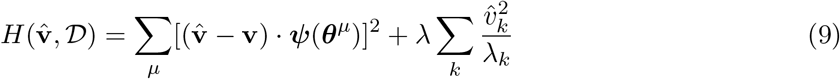

It is straightforward to verify that the *H*-minimizing coefficients are 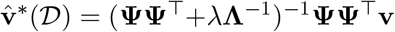, giving the learned function 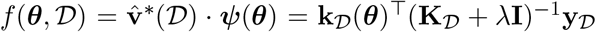, whose λ → 0 limit coincides with kernel interpolation. This allows us to characterize generalization without reference to learned readout weights **w**. The generalization error for this optimal function is

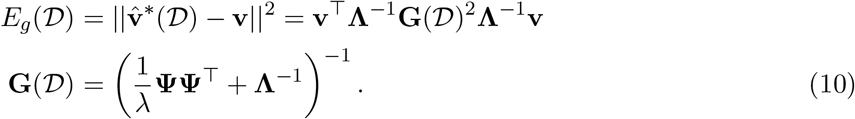

We note that the dependence on the randomly sampled dataset 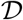 only appears through the matrix 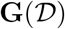. Thus to compute the *typical* generalization error we need to average over this matrix 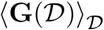. There are multiple strategies to perform such an average and we will study one here based on a partial differential equation which was introduced in [97, 98] and studied further in [26]. We describe in detail one method for performing such an average in App. 3B. After this computation, we find that the generalization error can be approximated at large *P* as

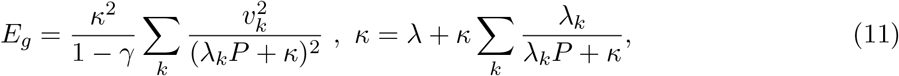

where 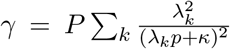, giving the desired result. We note that (11) defines the function *κ* implicitly in terms of the sample size *P*. Taking λ → 0 gives the generalization error of the minimum norm interpolant, which desribes the generalization error of the solution. This result was recently reproduced using the replica method from statistical mechanics in an asymptotic limit where the number of neurons and samples are large [26, 27]. Other recent works have verified our theoretical expressions on a variety of kernels and datasets [99, 100].

### 4.3 RNN Experiment

For the simulations in Figure 8 we integrated a rate based recurrent network model with *N* = 6000 neurons, time constant *τ* = 0.05 and gain *g* = 1.5. Each of the *P* = 80 randomly chosen angles *γ^μ^* generates a trajectory over *T* = 100 equally spaced points in *t* ∈ [0, 3]. The two dimensional input sequence is simply ***θ***(*t*) = *H*(*t*)*H*(1 – *t*) [cos(*γ^μ^*), sin(*γ^μ^*)]^⊤^ ∈ ℝ^2^. Target function for a delay *d* is **y**(*θ^μ^,t*) = *H*(1.5 + *d* – *t*)*H*(*t* – *d* – 1)[cos(*γ^μ^*), sin(*γ^μ^*)]^⊤^ which is nonzero for times *t* ∈ [1 + *d*, 1.5 + *d*]. In each simulation, the activity in the network is initialized to **u**(0) = **0**. The kernel gram matrix **K** ∈ ℝ^*PT*×*PT*^ is computed by taking inner products of the time varying code at for different inputs *γ^μ^* and at different times. Learning curves represent the generalization error obtained by randomly sampling 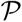 time points from the *PT* total time points generated in the simulation process and training readout weights **w** to convergence with gradient descent.

### 4.4 Data Analysis

#### 4.4.1 Data source and processing

Mouse V1 neuron responses to orientation gratings were obtained from a publicly available dataset [8, 9]. Two-photon calcium microscopy fluorescence traces were deconvolved into spike trains and spikes were counted for each stimulus, as described in [8]. The presented grating angles were distributed uniformly over [0, 2*π*] radians. Data pre-processing, which included z-scoring against the mean and standard deviation of null stimulus responses, utilized the provided code for this experiment, which also publicly available at https://github.com/MouseLand/stringer-et-al-2019. This preprocessing technique was used in all Figures in the paper. To reduce corruption of the estimated kernel from neural noise (trial-to-trial variability), we first trial average responses, binning the grating stimuli oriented at different angles *θ* into a collection of 100 bins over the interval from [0, 2*π*] and averaging over all of the available responses from each bin. Since grating angles were sampled uniformly, there is a roughly even distribution of about 45 responses in each bin. After trial averaging, SVD was performed on the response matrix **R**, generating the eigenspectrum and kernel eigenfunctions as illustrated in Figure 4. Figures 2, 3, 4, all used this data anytime responses to grating stimuli were mentioned.

In Figures 3C 4D and 5, the responses of mouse V1 neurons to ImageNet images [101] were obtained from a different publicly available dataset [10, 11]. The images were taken from 15 different classes from the Imagenet dataset with ethological relevance to mice (birds, cats, flowers, hamsters, holes, insects, mice, mushrooms, nests, pellets, snakes, wildcats, other animals, other natural, other man made). In the experiment in Figure 4D we use all images from the mice and birds category for which responses were recorded. The preprocessing code and image category information were obtained from the publicly available code base at https://github.com/MouseLand/stringer-pachitariu-et-al-2018b. Again, spike counts were obtained from deconvolved and z-scored calcium fluorescence traces. In the reconstruction experiment shown in Figure 5 we use the entire set of images for which neural responses were recorded.

#### 4.4.2 Generating RROS codes

In Figure 3, the randomly rotated codes are generated by sampling a matrix **Q** from the Haar measure on the set of *N*-by-*N* orthogonal matrices [20], and chosing a ***δ*** by solving the following optimization problem:

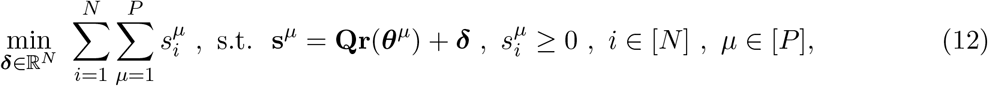

which minimizes the total spike count subject to the kernel and nonnegativity of firing rates. The solution to this problem is given by 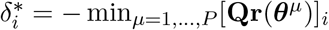.

#### 4.4.3 Comparing Sparsity of Population Codes

To explore the metabolic cost among the set of codes with the same inductive biases, we estimate the distribution of average spike counts of codes with the same inner product kernel as the biological code. These codes are generated in the form **s**^*μ*^ = **Qr**^*μ*^ + ***δ*** where ***δ*** solves the optimization problem

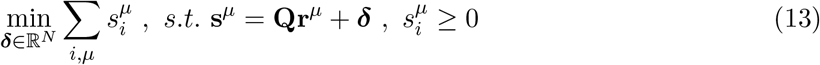

To quantify the distribution of such codes, we randomly sample **Q** from the Haar-measure on *O*(*N*) and compute the optimal ***δ*** as described above. This generates the aqua colored distribution in Figure 3 B and C.

We also attempt to characterize the most efficient code with the same inner product kernel

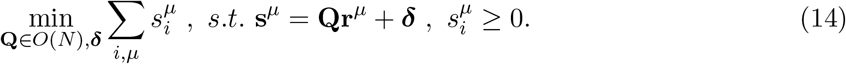

Since this optimization problem is non-convex in **Q**, there is no theoretical guarantee that minima are unique. Nonetheless, we attempt to optimize the code by starting **Q** at the identity matrix and conduct gradient descent in the tangent space *so*(*N*). Such updates take the form

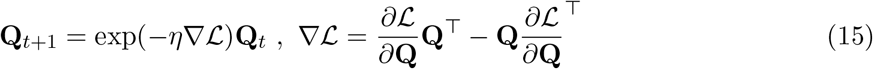

where exp(·) is the matrix exponential. To make the loss function differentiable, we incorporate the non-negativity constraint with a soft-minimum:

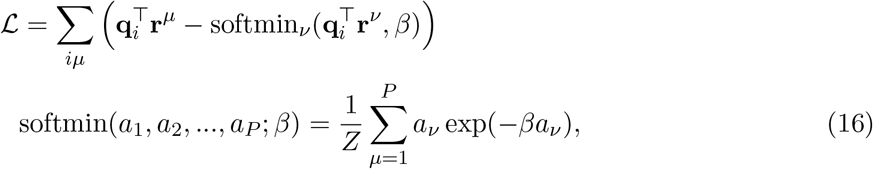

where *Z* = ∑_*ν*_ exp(−*βα_ν_*) is a normalizing constant and **Q** = [**q**_1_, …**q**_*N*_]. In the *β* → ∞ limit, this cost function converges to the exact optimization problem with non-negativity constraint. Finite *β*, however, allows learning with gradient descent. Gradients are computed with automatic differentiation in JAX [102]. This optimization routine is run until convergence and the optimal value is plotted as dashed red lines labeled “opt.” in Figure 3.

We show that our result is robust to different pre-processing techniques and to imposing bounds on neural firing rates in the Figure App.1. To demonstrate that our result is not an artifact of z-scoring the deconvolved signals against the spontaneous baseline activity level, we also conduct the random rotation experiment on the raw deconvolved signals. In addition, we show that imposing realistic constraints on the upper bound of the each neuron’s responses does not change our findings. We used a subset of *N* = 100 neurons and computed random rotations. However, we only accepted a code as valid if it’s maximum value was less than some upper bound *u_b_*. Subsets of N = 100 neurons in the biological code achieve maxima in the range between 3.2 and 4.7. We performed this experiment for *Ub* ∈ {3,4, 5} so that the artificial codes would have maxima that lie in the same range as the biological code.

#### 4.4.4 Lifetime and Population Sparseness

We compute two more refined measures of sparseness in a population code. For each neuron i we compute the lifetime sparseness *LS_i_* (also known as selectivity) and for each stimulus ***θ*** we compute the population sparseness *PS_θ_* which are defined as the following two ratios [22, 23, 24, 25]

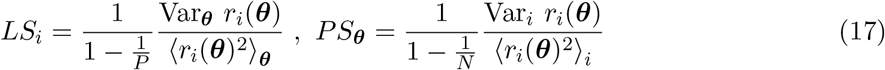

The normalization factors ensure that these quantities lie in the interval between (0,1). Intuitively, lifetime sparseness quantifies the variability of each neuron’s responses over the full set of stimuli, whereas population sparseness quantifies the variability of responses in the code for a given stimulus ***θ***.

#### 4.4.5 Fitting a Gabor model to mouse V1 kernel

Under the assumption of translation symmetry in the kernel *K*(*θ, θ*′), we averaged the elements of the over rows of the empirical mouse V1 kernel [9]

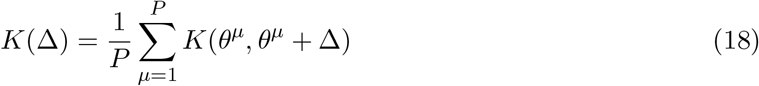

where angular addition is taken mod *π*. This generates the black dots in Figure 6 B. We aimed to fit a threshold-power law nonlinearity of the form *g_q,a_*(*z*) = max{0, *z* – *a*}^*q*^ to the kernel. Based on the Gabor model discussed above, we parameterized tuning curves as

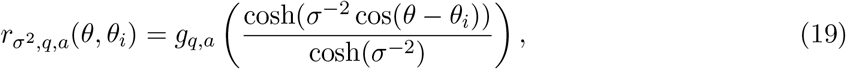

where *θ_i_* is the preferred angle of the i-th neuron’s tuning curve. Rather than attempting to perform a fit of *σ*^2^, 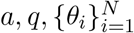 of this form to the responses of each of the ~ 20-k neurons, we instead simply attempt to fit to the population kernel by optimizing over (*σ*^2^, *a, q*) under the assumption of uniform tiling of *θ_i_*. However, we noticed that two of these variables *σ*^2^, *a* are constrained by the sparsity level of the code. If each neuron, on average, fires for only a fraction *f* of the uniformly sampled angles *θ*, then the following relationship holds between *σ*^2^ and *a*

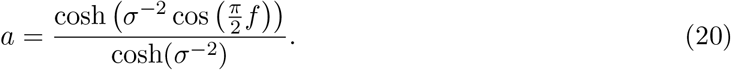

Calculation of the coding level *f* for the recorded responses allowed us to infer *a* from *β* during optimization. This reduced the free parameter set to (*σ*^2^, *q*). We then solve the following optimization problem

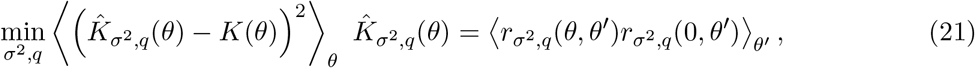

where integration over *θ_i_* is performed numerically. Using the Scipy Trust-Region constrained optimizer, we found (*q, σ*^-2^, *a*) = (1.7, 5.0, 0.2) which we use as the fit parameters in Figure 6.

## Lead contact

Requests for information should be directed to the lead contact, Cengiz Pehlevan.

## Data and code aVailability

Mouse V1 neuron responses to orientation gratings and preprocessing code were obtained from a publicly available dataset: https://github.com/MouseLand/stringer-et-al-2019, [8, 9].

Responses to ImageNet images and preprocessing code were obtained from another publicly available dataset, https://github.com/MouseLand/stringer-pachitariu-et-al-2018b [10, 11].

The code generated by the authors for this paper is also available https://github.com/Pehlevan-Group/sample_efficient_pop_codes

## Acknowledgements

We would like to thank Jacob Zavatone-Veth and Abdulkadir Canatar for useful comments and discussions about this manuscript. BB acknowledges the support of the NSF-Simons Center for Mathematical and Statistical Analysis of Biology at Harvard (award #1764269) and the Harvard Q-Bio Initiative. CP and BB would also like to acknowledge support from the Harvard Data Science Initiative.

## Appendix: Population codes enable learning from few examples by shaping inductive bias

### 1 Neural Code Decomposition

#### 1.1 Singular Value Decomposition of Continuous Population Responses

SVD of population responses is usually evaluated with respect to a discrete and finite set of stimuli. In the main paper, we implicitly assumed that a generalization of SVD to a continuum of stimuli. In this section we provide an explicit construction of this generalized SVD using techniques from functional analysis. Our construction is an example of the quasimatrix SVD defined in [1] and justifies our use of SVD in the main text.

For our construction, we note that Mercer’s theorem guarantees the existence of an eigendecomposition of any inner product kernel *K*(***θ, θ***′) in terms of a complete orthonormal set of functions 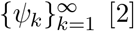 [2]. In particular, there exist a non-negative (but possibly zero) summable eigenvalues 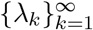 and a corresponding set of orthonormal eigenfunctions such that

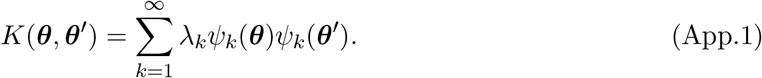

For a stimulus distribution *p*(***θ***), the set of functions 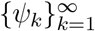 are orthonormal and form a complete basis for square integrable functions *L*_2_ which means

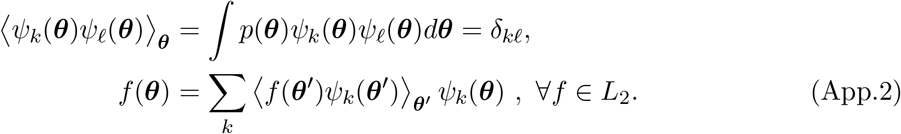

Next, we use this basis to construct the SVD. Each of the tuning curves *r_i_* ∈ *L*_2_ (assumed to be square integrable) can be expressed in this basis with the top *N* of the functions in the set 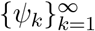

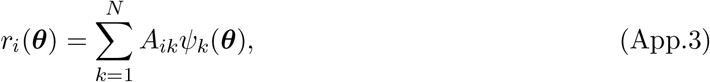

where we introduced a matrix **A** ∈ ℝ^*N*×*N*^ of expansion coefficients. Note that rank(**A**) ≤ *N*. We compute the singular value decomposition of the finite matrix **A**

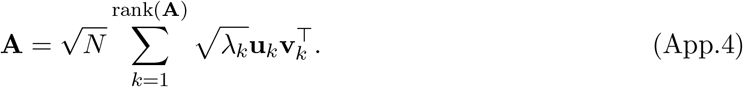

We note that the signal correlation matrix for this population code can be computed in closed form

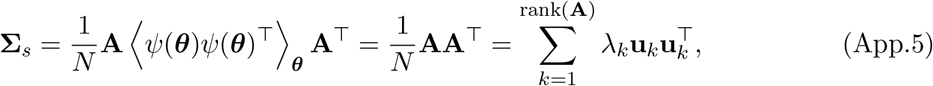

due to the orthonormality of {*ψ_k_*}. Thus the principal axes ***u***_*k*_ of the neural correlations are the left singular vectors of **A**. We may similarly express the inner product kernel in terms of the eigenfunctions

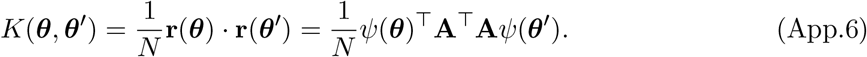

The kernel eigenvalue problem demands [2]

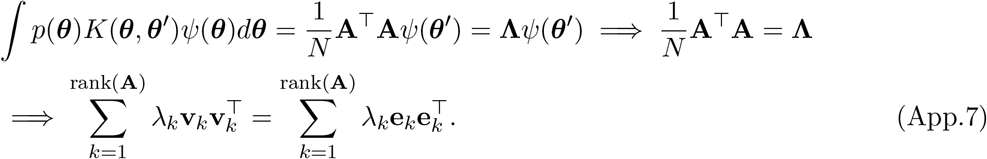

The **v**_*k*_ vectors must be identical to ±**e**_*k*_, the Cartesian unit vectors, if the eigenvalues are nondegenerate. From this exercise, we find that the SVD for **A** has the form 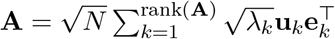. With this choice, the population code admits a singular value decomposition

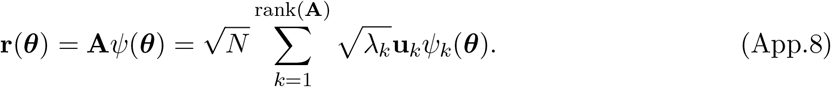

This singular value decomposition demonstrates the connection between neural manifold structure (principal axes **u**_*k*_) and function approximation (kernel eigenfunctions *ψ_k_*). This singular value decomposition can be verified by computing the inner product kernel and the correlation matrix, utilizing the orthonormality of {**u**_*k*_} and {*ψ_k_*}. This exercise has important consequences for the space of learnable functions, which is at most rank(**A**) dimensional since linear readouts lie in span 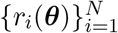.

#### 1.2 Discrete Stimulus Spaces: Finding Eigenfunctions with Matrix Eigende-composition

In our discussion so far, our notation suggested that ***θ*** take a continuum of values. Here we want to point that our theory still applies if ***θ*** take a discrete set of values. In this case, we can think of a Dirac measure 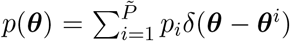, where *i* indexes all the 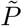 values ***θ*** can take. With this choice

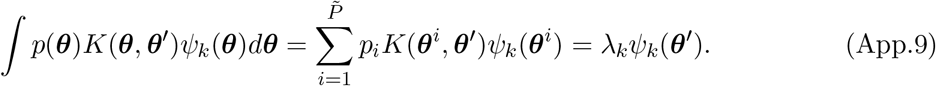

Demanding this equality for ***θ′*** = ***θ***^*i*^, 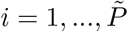 generates a matrix eigenvalue problem

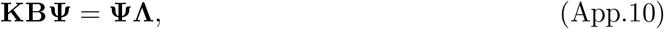

where **B**_*ij*_ = *δ_ij_p_i_*. The eigenfunctions over the stimuli are identified as the columns of **Ψ** while the eigenvalues are the diagonal elements of ***Λ***_*kℝ*_ = *λ_k_ δ_kℝ_*.

##### Experimental considerations

In an experimental setting, a finite number of stimuli are presented and the SVD is calculated over this finite set regardless of the support of *p*(***θ***). This raises the question of the interpretation of this SVD and its relation to the inductive bias theory we presented. Here we provide two interpretations.

In the first interpretation, we think of the empirical SVD as providing an estimate of the SVD over the full distribution *p*(***θ***). To formalize this notion, we can introduce a Monte-Carlo estimate of the integral eigenvalue problem

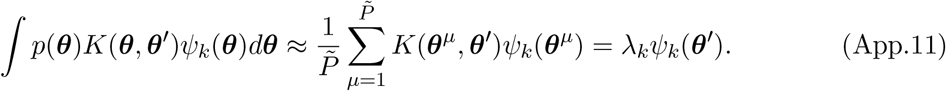

For this interpretation to work, the experimenter must sample the stimuli from *p*(***θ***), which could be the natural stimulus distribution. Measuring responses to a larger number of stimuli gives a more accurate approximation of the integral above, which will provide a better estimate of generalization performance on the true distribution *p*(***θ***).

In the second interpretation, we construct an empirical measure on 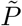 experimental stimulus values 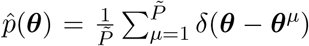, and consider learning and generalization over this distribution. This allows the application of our theory to an experimental setting where 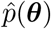 is designed by an experimenter. For example, the experimenter could procure a complicated set of 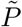 videos, to which an associated function *y*(*θ*) must be learned. After showing these videos to the animal and measuring neural responses, the experimenter could compute, with our theory, generalization error for a uniform distribution over this full set of 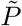 videos. Our theory would predict generalization over this distribution after providing supervisory feedback for only a strict subset of 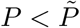 videos. Under this interpretation, the relationship between the integral eigenvalue problem and matrix eigenvalue problem is exact rather than approximate

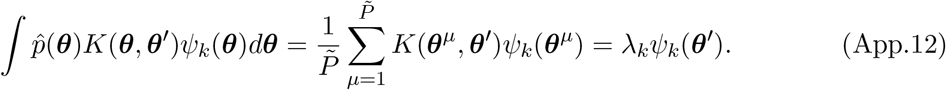

Demanding either of (App.11) or (App.12) equalities for **θ**′ = **θ**^*ν*^, *v* = 1, …, *P* generates a matrix eigenvalue problem

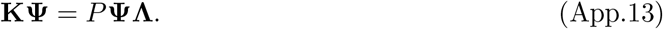

The eigenfunctions restricted to {***θ***^*μ*^} are identified as the columns of **Ψ** while the eigenvalues are the diagonal elements of Λ_*kℓ*_ = λ_*k*_ *δ_kℓ_*. For the case where *N* and *P* are finite, the spectrum obtained through eigendecomposition of the kernel **K** is the same as would be obtained through the finite *N* signal correlation matrix **Σ**_*s*_, since they are inner and outer products of trial averaged population response matrices **R**.

#### 1.3 Translation Invariant Kernels

For the special case where the data distribution 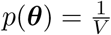 is uniform over volume *V* and the kernel is translation invariant *K*(**θ, *θ′***) = *κ*(***θ – θ′***), the kernel can be diagonalized in the basis of plane waves

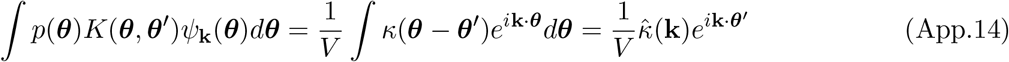

The eigenvalues are the Fourier components of the Kernel 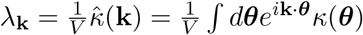 while the eigenfunctions are plane waves *ψ*_**k**_(***θ***) = *e*^*i*k.***θ***^. The set of admissible momenta 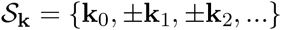 are determined by the boundary conditions. The diagonalized representation of the kernel is therefore

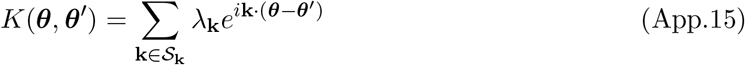

For example, if the space is the torus 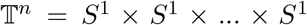, then the space of admissable momenta are the points on the integer lattice 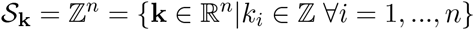. Reality and symmetry of the kernel demand that Im(λ_**k**_) = 0 and λ_**k**_ = λ**k** ≥ 0. Most of the models in this paper consider *θ* ~ Unif (*S*^1^), where the kernel has the following Fourier/Mercer decomposition

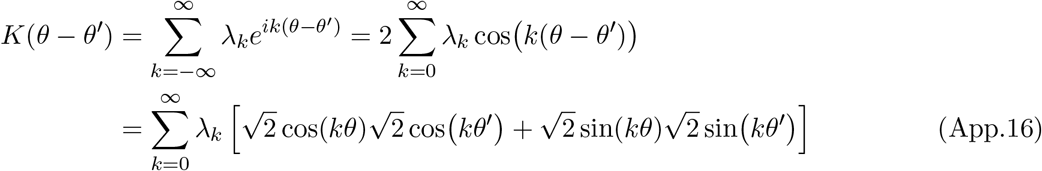

where we invoked the simple trigonometric identity cos(*a* – *b*) = cos(*a*)cos(*b*) + sin(*a*) sin(*b*). By recognizing that 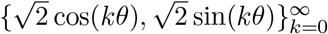 form a complete orthonormal set of functions with respect to Unif (*S*^1^), we have identified this as the collection of kernel eigenfunctions.

#### 1.4 Invariant Kernels Possess Invariant Eigenfunctions

Suppose the kernel *K*(***θ, θ***’) is invariant to some set of transformations 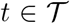, by which we mean that

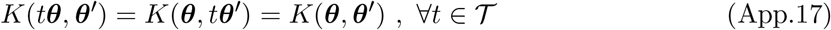

We will now show that any eigenfunction of such a kernel with nonzero eigenvalue must be an invariant function. Let *ψ_k_*(***θ***) be an eigenfunction with eigenvalue λ_*k*_ > 0, then

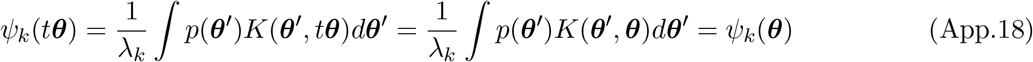

This establishes that all functions which depend on 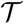 transformations must necessarily lie in the null-space of *K*.

### 2 Alternative Neural Codes with Same Kernel

#### 2.1 Orthogonal Transformations are Sufficient for Linear Kernel-Preserving Transformations

We will now show that for any linear transformation 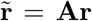 which preserves the inner product kernel *K*(***θ, θ***′), there exists an orthogonal matrix **Q** such that 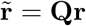.

##### Proof.

Let 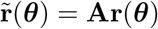 for all stimuli ***θ***. To preserve the kernel, we must have

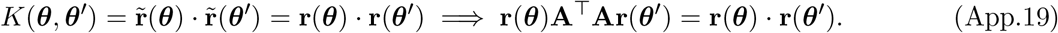

Taking projections against each of the orthonormal eigenfunctions *ψ_ℓ_*(*θ*) (see Appendix 1A), we define vectors **u**_*k*_ as 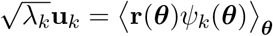, allowing us to express the SVD of the population code 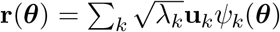. These vectors {**u**_*k*_} are orthonormal **u**_*k*_ · **u**_*ℓ*_ = *δ_kℓ_* since, by the definition of the kernel eigenfunctions *ψ_k_*,

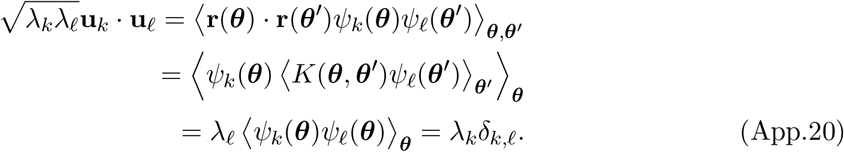

Since **r**(***θ***) and 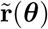 have the same inner product kernel, they must posess the same kernel eigenfunctions *ψ_k_* and kernel eigenvalues λ_*k*_, which are identified through the eigenvalue problem ∫*p*(***θ***)*K*(***θ, θ***′)*ψ_k_*(***θ***)*d****θ*** = λ*_k_ψ_k_*(***θ***). We therefore have the following two singular value decompositions for **r** and 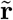

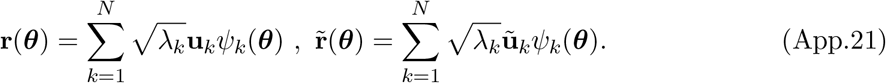

where 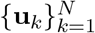 and 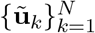 are both complete sets of orthonormal vectors (the sums above run over possible zero eigenvalues). Taking the equation 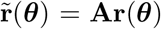, we multiply both sides of the equation by *ψ_k_*(***θ***) and average over ***θ*** giving

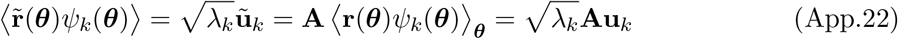

For an eigenmode *k* with positive eigenvalue λ_*k*_ > 0, this implies 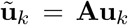, while there is no corresponding constraint for the null modes with λ_*k*_ = 0. However, the action of **A** on the nullspace of the code has no influence on 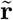 so there is no loss in generality to restrict consideration to transformations **A** which satisfy 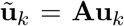 for all *k* ∈ [*N*] (rather than just the λ_*k*_ > 0 modes). This choice gives 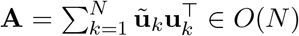. Thus, the space of codes 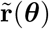 with equivalent kernels to **r**(***θ***) · **r**(***θ***′) generated through linear transformations is equivalent to all possible orthogonal transformations of the original code {**Qr**(***θ***): **Q** ∈ *O*(*N*)}.

#### 2.2 Necessary Conditions for Optimally Sparse Codes

Next we argue why optimally sparse codes should be lifetime and population selective. We consider the following optimization problem: find a non-negative neural responses **S** ∈ ℝ^*N*×*P*^ and baseline vector ***δ*** ∈ ℝ^*N*^ so that baseline subtracted responses **R** = **S** – ***δ*1**^⊤^ realize a desired inner product kernel **K** ∈ ℝ^*P*×*P*^ and have minimal total firing. This is equivalent to finding the most metabolically efficient code among the space of codes with equivalent inductive bias. Mathematically, we formulate this problem as

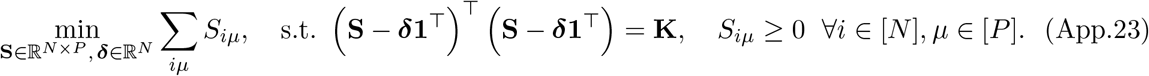

To enforce the constraints for the definition of the kernel and the non-negativity of the responses, we introduce the following Lagrangian

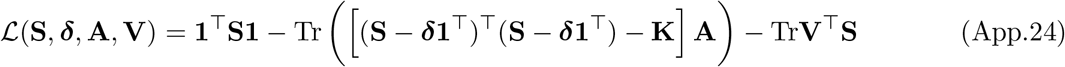

where **1** is the vector containing all ones, the Lagrange multiplier matrix **A** enforces the definition of the kernel and the KKT multiplier matrix **V** enforces the non-negativity constraints for each element of **S**. The KKT conditions require that any local optimum of the objective would have to satisfy the following equations [3]

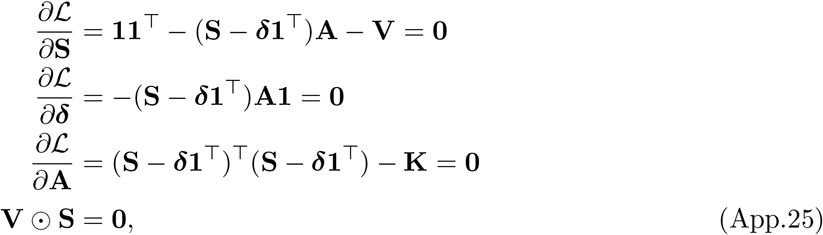

where ⊙ denotes the element-wise Hadamard product. Using the complementary slackness condition **S** ⊙ **V** = **0**, and the first optimality condition 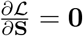, we have

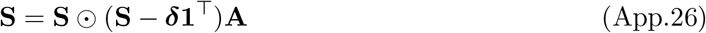

Therefore, for any neuron-stimulus pair (*i,μ*), either *S_iμ_* = 0 or ∑_*ν*_∈[*P*]*(S_iν_* – *δ_i_*)*A_νμ_* = 1. Further, under the condition that **K** is full rank, we conclude that for any stimulus *μ*, ∑_*ν*_∈[*P*] *A_μν_* = 0 from the equation 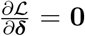. Let 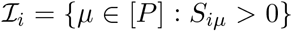 represent the set of stimuli for which neuron *i* fires. We will call this the *receptive field set* for neuron *i*. Let **B**_(*i*)_ ∈ ℝ^*P*×*P*^ have entries

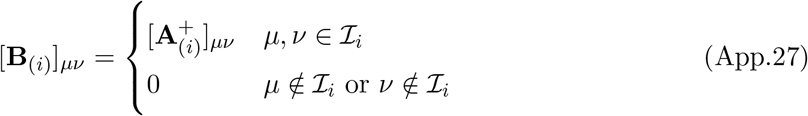

where the matrix **A**_(*i*)_ is the 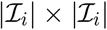 minor of **A** obtained by taking all rows and columns with indices 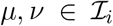, and **A**^+^ denotes pseudo-inverse of **A**. Then the *i*-th neuron’s tuning curve is a function of the index set 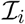 the baseline *δ_i_* and the neuron-independent *P* × *P* matrix 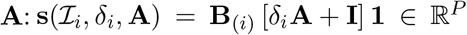. The non-negativity constraint for neuron *i*’s tuning curve implies that 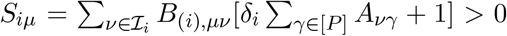 for all 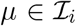. To satisfy the definition of the kernel, we have the following constraint on the matrix **A**, the index sets 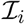 and baselines *δ_i_*

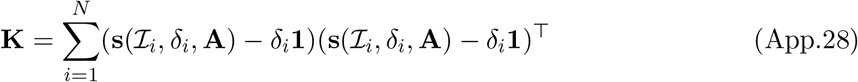

This equation implictly defines the index sets 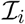 the baselines *δ_i_* and the KKT matrix **A**. We see that, in order to fit an arbitrary kernel, the receptive field sets 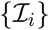 and baselines *δ_i_* for each neuron must be sufficiently diverse since otherwise only a low rank kernel matrix can be achieved from the optimally sparse code. As a concrete example, suppose that 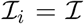 so that **B**_(*i*)_ = **B** and *δ_i_* = *δ* for all *i*. For example, this could occur if each neuron fired for every possible stimulus. In this case, the kernel would be rank one: 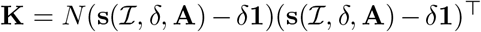. In order to achieve a higher rank code there must be sufficient diversity of the receptive fields 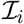. Thus the only way for optimally sparse codes to realize high rank kernels **K** is to have neurons to have different receptive field sets 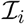. The necessary optimality conditions thus reveal a preference for sparse neural tuning curves to have high *lifetime sparseness*; to achieve diverse index sets 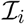, any given neuron will fire only for a unique subset of the possible stimuli.

### 3 Theory of Generalization

#### 3.1 Convergence of the delta-rule

Gradient descent training of readout weights **w** on a finite sample of size *P* converges to the kernel regression solution [4, 5, 6]. Let 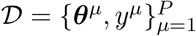 be the dataset with samples ***θ***^*μ*^ and target values *y^μ^*. We introduce a shorthand **r**^*μ*^ = **r**(***θ***^*μ*^) for convenience. The empirical loss we aim to minimize is a sum of the squared losses of each data point in the training set

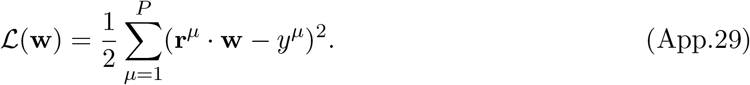

Performing gradient descent updates

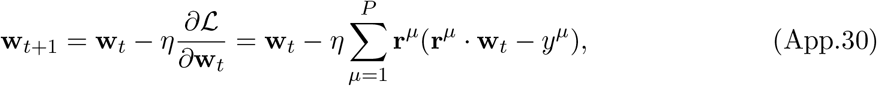

recovers the delta rule that we discussed in the main text [7, 8]. Letting the empirical response matrix **R** = [**r**^1^,…,**r**^*P*^] ∈ ℝ^*N*×*P*^ have a SVD 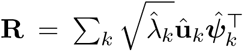, and expanding the weights 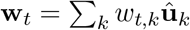 and labels 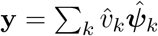 in their respective SVD bases, we find

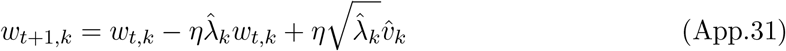

For all directions with 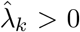, the dynamics converge to the unique fixed point 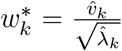, while for all modes with 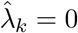, the weights remain at 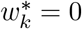. Thus

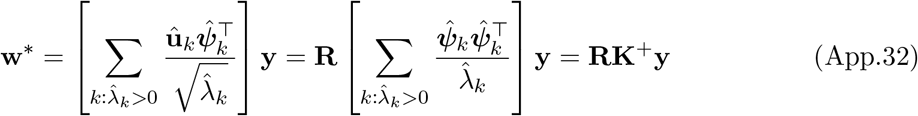

where **K**^+^ is the Moore-Penrose inverse of the kernel matrix *K_μν_* = *K*(***θ***^*μ*^, ***θ***^*ν*^). The predictions of the learned function are given by *f* = **w**\* · **r**(***θ***) which can be expressed as

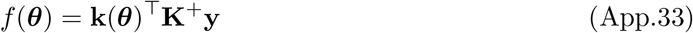

The fact that the solution can be written in terms of a linear combination of 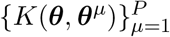 is known as the representer theorem [9, 2]. A similar analysis for nonlinear readouts where *f* (***θ***) = *g* (**w** · **r**(***θ***)) is provided in Appendix 3G.

##### 3.1.1 Weight Decay and Ridge Regression

We can introduce a regularization term in our learning problem which penalizes the size of the readout weights. This leads to a modified learning objective of the form

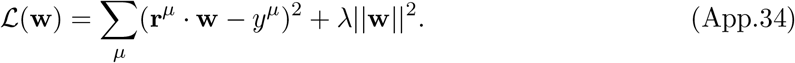

Inclusion of this regularization alters the learning rule through *weight decay* **w**_*t*+1_ = (1 – *η*λ)**w**_*t*_ + *η* ∑_*μ*_ **r**^*μ*^(**r**^*μ*^ · **w**_*t*_ – *y^μ^*), which multiplies the existing weight value by a factor of 1 – *η*λ before adding the data dependent update. This learning problem and gradient descent dynamics have a closed form solution

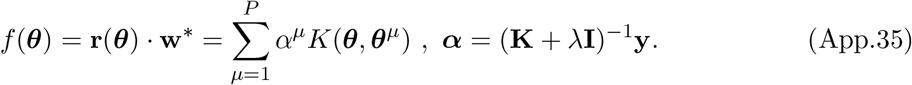

The generalization benefits of explicit regularization through weight decay is known to be related to the noise statistics in the learning problem [10]. We simulate weight decay only in Figure 6C, where we use λ = 0.01 ∑2_*k*_ λ_*k*_ to improve numerical stability at large *P*.

#### 3.2 Computation of Learning Curves

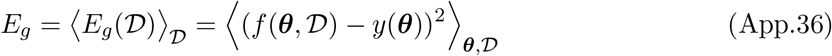

where 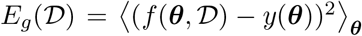 is the generalization error for a certain sample 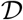 of size *P* and 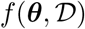 is the kernel regression solution for 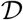 [11, 10]. The typical or average case error *E_g_* is obtained by averaging over all possible datasets of size *P*. This average case generalization error is determined solely by the decomposition of the target function *y*(**x**) along the eigenbasis of the kernel and the eigenspectrum of the kernel. This diagonalization takes the form

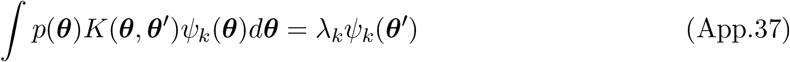

Since the eigenfunctions form a complete set of square integrable functions, we expand both the target function *y*(***θ***) and the learned function *f* (***θ***) in this basis

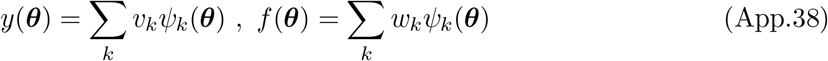

Due to the orthonormality of the kernel eigenfunctions {*ψ_k_*}, the generalization error for any set of coefficients **w** is

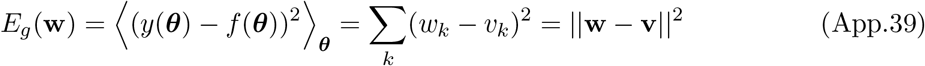

We now introduce training error, or empirical loss, which depends on the disorder in the dataset 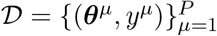

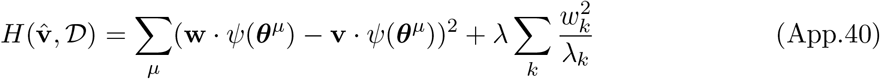

It is straightforward to verify that the optimal 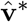 which minimizes 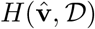 is the kernel regression solution for kernel with eigenvalues {λ_*k*_} when λ → 0. The optimal weights 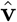 can be identified through the first order condition 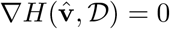 which gives

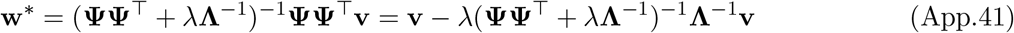

where Ψ_*k,μ*_ = *ψ_k_*(***θ***^*μ*^) are the eigenfunctions evaluated on the training data and *Λ_k,ℓ_* = *δ_k,ℓ_,λ_k_* is a a diagonal matrix containing the kernel eigenvalues. The generalization error for this optimal solution is

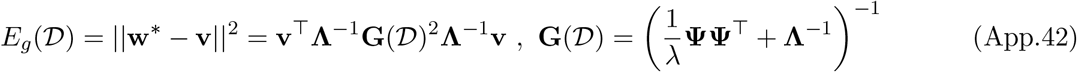

We note that the dependence on the randomly sampled dataset 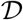 only appears through the matrix 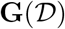. Thus to compute the *typical* generalization error we need to average over this matrix 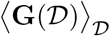. There are multiple strategies to perform such an average and we will study one here based on a partial differential equation which was introduced in [12, 13] and studied further in [11, 10]. In this setting, we denote the average matrix 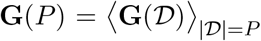 for a dataset of size *P*. We first will derive a recursion relationship using the Sherman Morrison formula for a rank-1 update to an inverse matrix. We imagine adding a new sampled feature vector *ϕ* to a dataset *ψ* with size *P*. The average matrix **G**(*P* + 1) at *P* + 1 samples can be related to **G**(*P*) through the Sherman Morrison rule

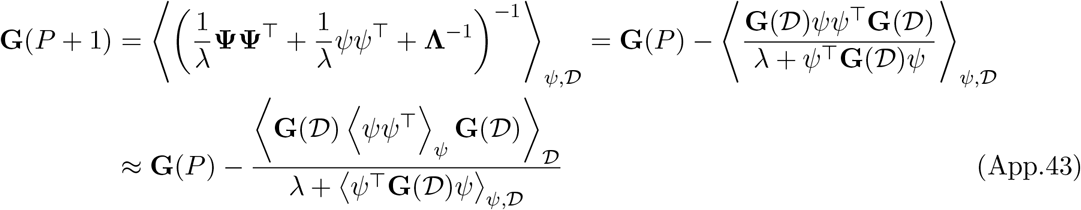

where in the last step we approximated the average of the ratio with the ratio of averages. This operation, is of course, unjustified theoretically, but has been shown to produce accurate learning curves [11, 13]. Since the chosen basis of kernel eigenfunctions are orthonormal, the average over the new sample is trivial 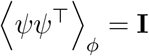. We thus arrive at the following recursion relationship for **G**

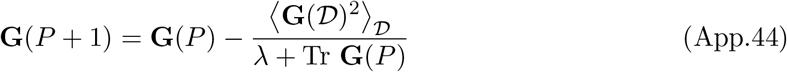

By introducing an additional source *J* so that 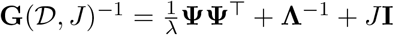, we can relate 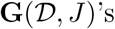 first and second moments through differentiation

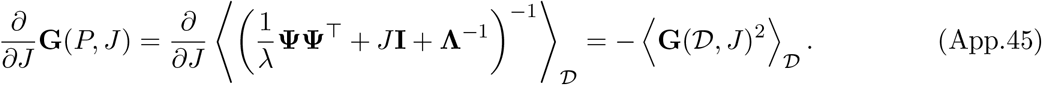

Thus the recursion relation simplifies to

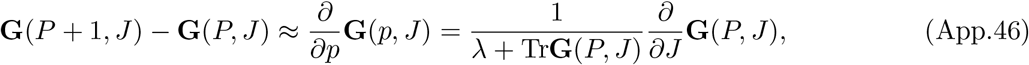

where we approximated the finite difference in *P* as a derivative, treating *P* as a continuous variable. Taking the trace of both sides and defining *κ*(*P, J*) = λ + Tr**G**(*P, J*) we arrive at the following quasilinear PDE

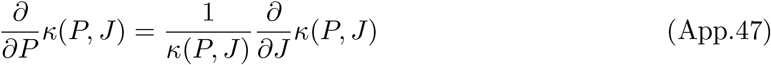

with the initial condition *κ*(0, *J*) = λ + Tr(**Λ**^-1^ + *J***I**)^-1^. Using the method of characteristics, we arrive at the solution 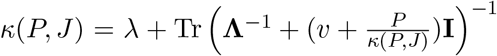. Using this solution to *κ*, we can identify the solution to **G**

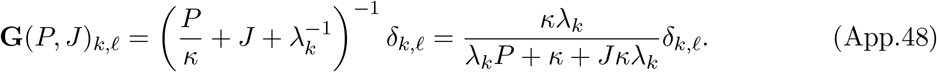

The generalization error, therefore can be written as

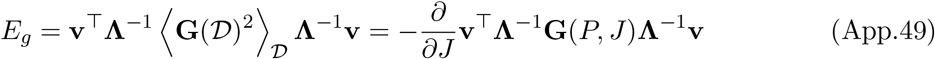

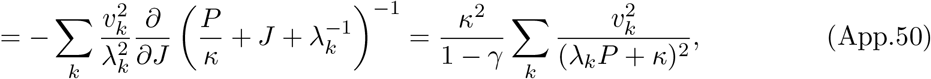

where 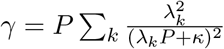, giving the desired result. Note that *κ* depends on *J* implicitly, which is the source of the 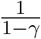 factor. This result was recently reproduced using techniques from statistical mechanics [11, 10].

#### 3.3 Spectral Bias and Code-Task Alignment

Through implicit differentiation it is straightforward to verify that the ordering of the mode errors 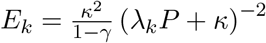 matches the ordering of the eigenvalues [10]. Let λ_*k*_ > λ_ℓ_, then we have

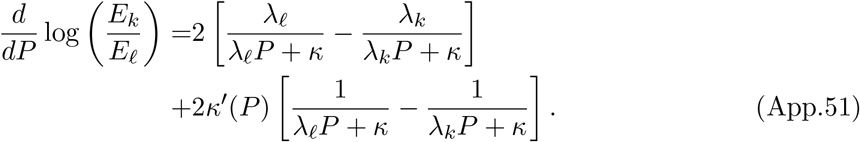

Since λ_ℓ_ < λ_*k*_, the first bracket must be negative and the second bracket must be positive. Further, it is straightforward to compute that 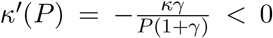. Therefore λ_*k*_ > λ_ℓ_ implies 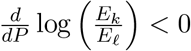 for all *P*. Since 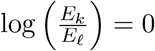 we therefore have that log (*E_k_/E_ℓ_*) < 0 for all *P* and consequently *E_k_* < *E_ℓ_*. Modes with larger eigenvalues *λ_k_* have lower normalized mode errors *E_k_*. This observation can be used to prove that target functions acting on the same data distribution with higher cumulative power distributions *C*(*k*) for all *k* will have lower generalization error normalized by total target power, *E_g_*(*P*)/*E_g_* (0), for all *P*. Proof can be found in [10].

#### 3.4 Asymptotic power law scaling of learning curves

##### Exponential Spectral Decays

First, we will study the setting relevant to the von-Mises kernel where λ_*k*_ ~ *β^k^* and 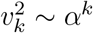 where *α,β* < 1. This exponential behavior accounts for differences in bandwidth between kernels which modulates the base *β* of the exponential scaling of λ_*k*_ with *k*. We will approximate the sum over all mode errors with an integral

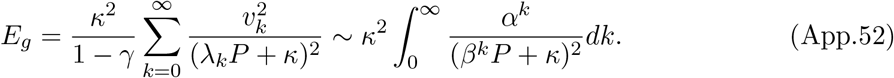

If we include a regularization parameter λ, then *κ* ~ λ as *P* → ∞. With this fact, we can therefore approximate the integral at large *P* by splitting it up into all *k* < *k*\* = ln(*P*/λ)/ln(1/*β*) and *k* > *k*\*.

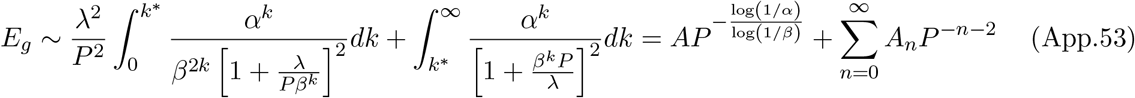

for *P*-independent constants *A* and *A_n_*. Thus, we obtain a power law scaling of the learning curve *E_g_* which is dominated at large *P* by 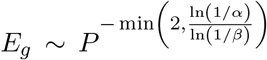. For the von-Mises kernel we can approximate the spectra with λ_*k*_ ~ *σ*^-2*k*^ and 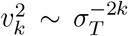 giving rise to a generalization scaling scaling 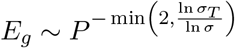.

##### Power Law Spectral Decays

The same arguments can be applied for power law kernels λ_*k*_ ~ *k*^−*b*^ and power law targets 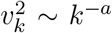, which is of interest due to its connection to nonlinear rectified neural populations. In this setting, the generalization error is

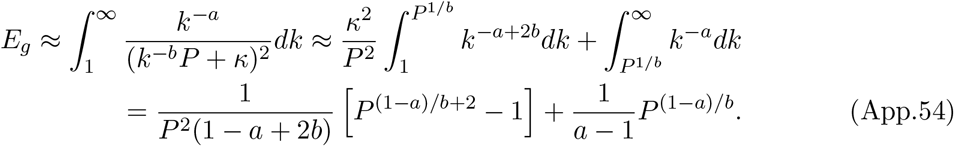

We see that there are two possible power law scalings for *E_g_* with the exponents (*a* – 1)/*b* and 2. At large *P* this formula will be dominated by the term with minimum exponent so *E_g_* ~ *P* – min(*a*–1,2*b*)/*b*.

#### 3.5 Laplace Kernel Generalization

We calculate similar learning curves as we did for the von-Mises kernel but with Laplace kernels to show that our results is not an artifact of the infinite differentiability of the Von Mises kernel. Each of these Laplace kernels has the same asymptotic power law spectrum λ_*k*_ ~ *o*(*k*^-2^), exhibiting a discontinuous first derivative (Figure App.5 A). Despite having the same spectral scaling at large *k*, these kernels can give dramatically different performance in learning tasks, again indicating the influence of the top eigenvalues on generalization at small *P* (Figure App.5). Again, the trend for which kernels perform best at low *P* can be reversed at large P. In this case, all generalization errors scale with *E_g_* ~ *P*^-2^ (Figure App.5B). More generally, our theory shows that if the task power spectrum and kernel eigenspectrum are both falling as power laws with exponents *a* and *b* respectively, then the generalization error asymptotically falls with a power law, *E_g_* ~ *P*^-min(*a*–1,2*b*)/*b*^ (Methods) [11]. This decay is fastest when 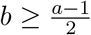 for which *E_g_* ~ *P*^−2^. Therefore, the tail of the kernel’s eigenvalue spectrum determines the large sample size behavior of the generalization error for power law kernels. Small sample size limit is still governed by the bulk of the spectrum.

#### 3.6 Learning with Multiple Output Channels

Our theory is not limited to scalar target functions but rather can be easily extended to multiple output functions *y*_1_,…,*y_C_* from the same data, if for example the task requires computing class membership for *C* categories. In this setting, each data point has the form (***θ***^*μ*^, **y**^*μ*^) with **y**^*μ*^ ∈ ℝ^*C*^. For these *C* classes, the generalization error takes the form

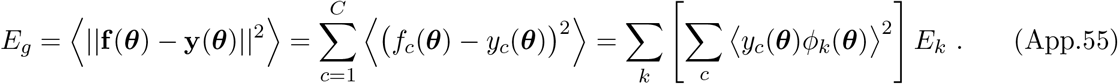

We therefore find that the generalization error in the multi-class setting is the same as the *E_g_* obtained for a single scalar target function with power spectrum 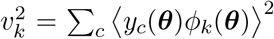 [11, 10]. The relevant cumulative power distribution measures the fraction of total output variance captured by the first *k* eigenfunctions of the population code

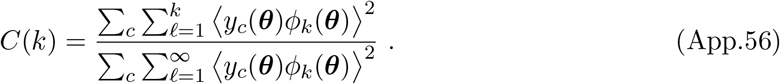

#### 3.7 Convergence of Delta-Rule for Nonlinear Readouts

In this section, we consider gradient descent dynamics on a least squares cost with nonlinear readout function. Let 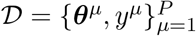 be the dataset with samples ***θ***^*μ*^ and target values *y^μ^*. We introduce a shorthand **r**^*μ*^ = **r**(***θ***^*μ*^) for convenience. We consider output neurons which produce activity *f* (***θ***) = *g*(**w** · **r**(***θ***)) for invertible nonlinear function *g* with non-vanishing gradient. The empirical loss we aim to minimize is a sum of the squared losses of each data point in the training set

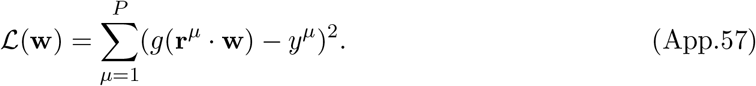

Performing gradient descent updates

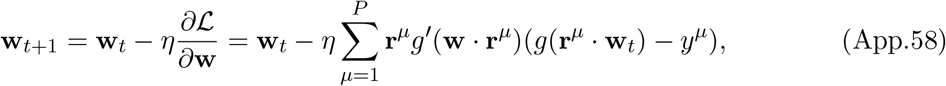

recovers the delta rule that we discussed in the main text for *g*(*z*) = *z* [7, 8]. We define the variables ***z***_*t*_ = **R**^⊤^**w**_*t*_ which satisfy the dynamics

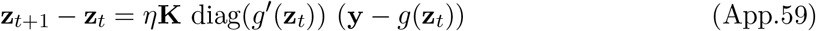

The possible fixed points of this system are the **z** which satisfy **K** diag(*g*′(**z**)) (**y**–*g*(**z**)) = 0. We can make further progress under the condition that **K** is full rank (generally true in overparameterized setting *N* > *P*), the activation function is monotonic *g*′(*z*) > 0 for all *z* and is invertible (some examples include softplus *g*(*z*) = ln(1 + *e^z^*), sigmoids such as *g*(*z*) = tanh(*z*), erf(*z*), etc.). If *y^μ^* are in the range of *g*, then we obtain the simple fixed point condition **z**\* = *g*^-1^(**y**). From this condition, we can infer the learned function. First note that 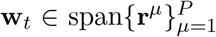 for all *t* so that **w**\* = **R*α***\*. Multiplying both sides of this last expression by **R**^⊤^, we find

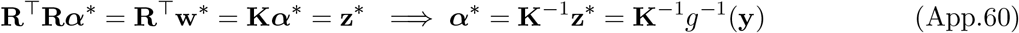

which is legitimate under the assumption that **K** was full rank. Now, the final learned function is merely *f*(***θ***) = *g*(**w**\* · **r**(***θ***)) = *g* **(r**(***θ***)^⊤^**RKV**(**y**)) = *g* **k**(***θ***)^⊤^ **K**^-1^ *g*(**y**)).

#### 3.8 Comparison of our Theory with Recursive Least Squares Adaptive Filter

In the under-parameterized regime where *P* > *N*, the asymptotic error of recursive least squares adaptive filter has been obtained in prior works [14, 15]. Let **w** represent the weights and **r**_*μ*_ represent the *μ*-th presented example. We aim to reproduce a noisy ground truth *y_μ_* = **w**\* · **r**_*μ*_ + *ϵ_μ_*. Analysis of the recursive least squares filter in the under-parameterized regime *P* > *N* + 1 can be performed exactly

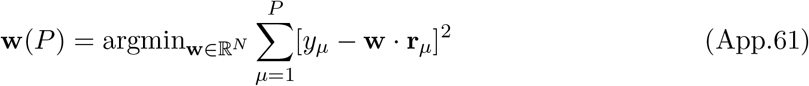

This has a unique minimizer when *P* > *N*. Letting 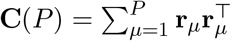

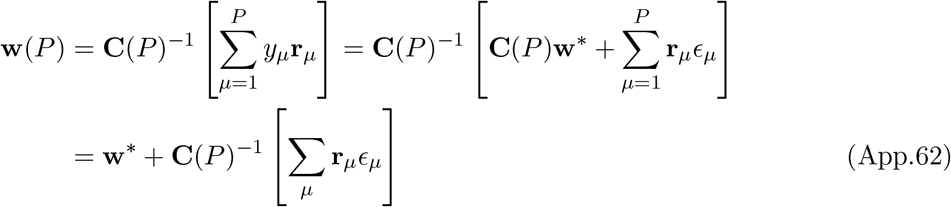

This filter can be computed efficiently in an online manner using the Recursive Least Squares (RLS) algorithm. This recursion is obtained through the formula for the rank-one update to an inverse matrix

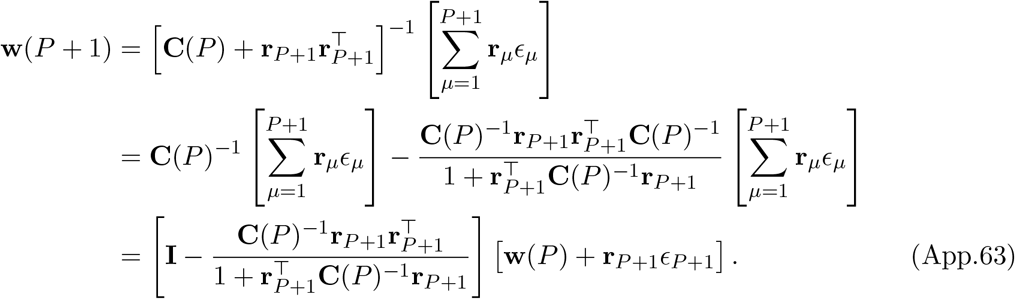

Under the assumption that the noise has covariance 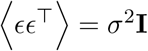 and the neural responses are Gaussian 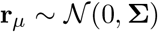, the excess generalization error can be obtained exactly using the formula for the average of an inverse Wishart matrix 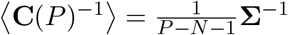 giving

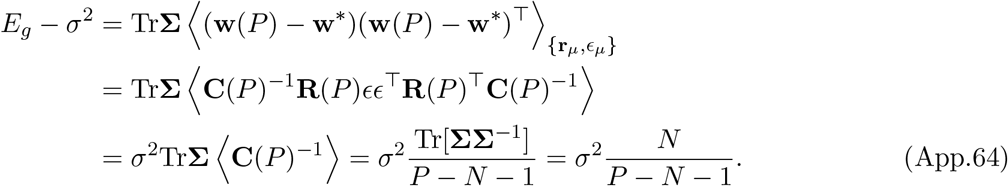

This analysis shows that RLS is consistent in the sense that as *P* → ∞ with *N* held constant the generalization error reach its theoretical minimum *σ*^2^.

Though our main paper focuses on noise free estimation (*σ*^2^ = 0) and the over-parameterized (*N* > *P*) rather than under parameterized case (which is what is discussed above), the above result agrees with what is obtained by our theory in the under-parameterized case since, in the *P* > *N* limit, *κ* = 0 and 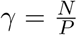. From our previous work, the error in the underparameterized noisy case for large *P* is [10]

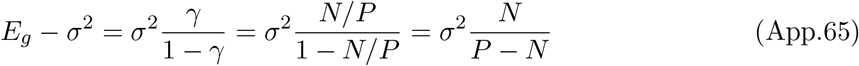

which is agrees with the exact result up to an error which is negligible in the large *N, P* limit. In this over-parameterized setting, the only contribution to the error comes from the explicit noise in the target function *σ*^2^.

We note however, that the above technique does not generalize to the *overparameterized* setting where the system is trying to learn from a small number of examples *P* < *N*. In this case, when *σ* = 0, the delta-rule converges to the minimum norm solution which has the form

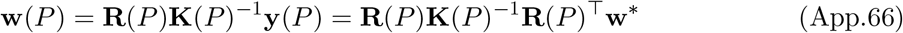

where **K**(*P*) = **R**(*P*)^⊤^**R**(*P*). We see that the learned weight vector **w**(*P*) is a random rank-*P* projection of the optimal weight vector **w**\*. The challenging task is to compute an average over the random data **R** to get generalization error

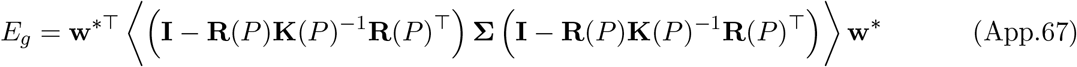

We solve this problem by first making a relaxation to the λ → 0 limit of ridge regression

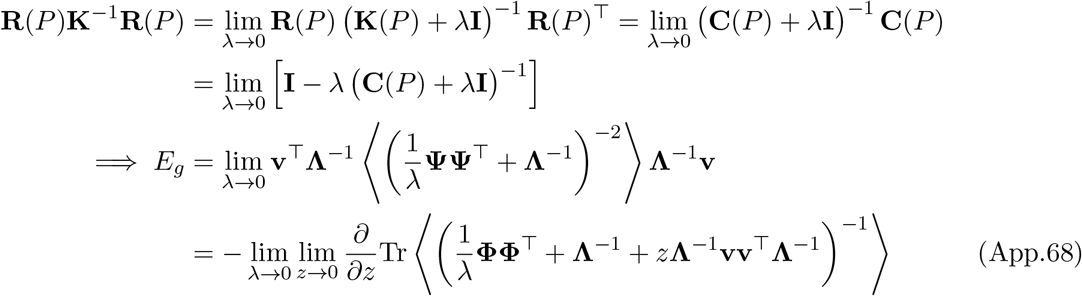

where **v** = **Λ**^1/2^**U**^⊤^**w**\*. This average can be characterized exactly for arbitrary λ, **Λ**, **w**\* only in asymptotic limits where the number of samples *P* and the number of features N are both large. Though in this present work, we resort to the PDE approximation, our theory can also be derived with the replica method from disordered systems [11, 10].

#### 3.9 Comparison of our Theory with Wiener Filter and Equivalent Kernel

We will provide a discussion of how the theory of generalization used in our paper differs from these older results, such as those discussed in Chapter 7 of [2]. Many of the older asymptotic results for kernel regression with a ridge λ can be obtained through an averaging argument (see for example Rasmussen and Williams Chapter 7.1 and 7.2 [2]). In the limit where the number of independent samples is large, we have by the strong law of large numbers, a convergence of the empirical loss to an expectation integral

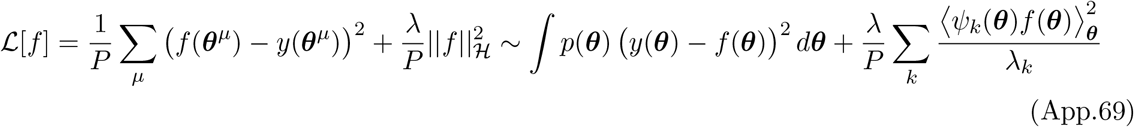

for a complete orthonormal family of eigenfunctions *ψ_k_*. The minimizer to the above functional *f* and its corresponding generalization error *E_g_* have the form

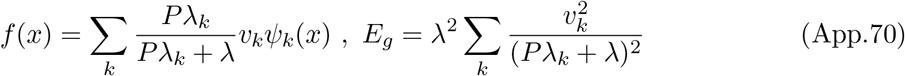

for target coefficients *v_k_* = 〈*y*(*x*)*ψ_k_*(*x*)〉0 This equation (which is equivalent to the Wiener filter when *ψ_k_* are plane waves; see [2] Ch 7.1) is somewhat sensible in that this function approaches *y*(*x*) as *P* → ∞, and agrees with our theory in the limit of large ridge parameter λ → ∞. Further, the large eigenvalue modes are learned at smaller values of *P*: it takes roughly *P* ≈ λ/λ_*k*_ samples to learn mode *k*. Note, however, that this theory is essentially useless in the interpolating limit λ → 0^+^ where it predicts that the learning curve has no dependence on sample size. This occurs because the fluctuations in the loss function from sample to sample (quenched disorder in statistical physics language) have been disregarded. Our analysis does not suffer from this issue and predicts the effects of sampling induced fluctuations in the loss function. Such sample fluctuations cannot typically be disregarded in thermodynamic limits where both dimension and dataset size are taken to infinity simultaneously [16]. A more complete discussion of the error due to sampling noise can be found in [10]. Our theory, in contrast to the Wiener filter described above, gives the following average predictor.

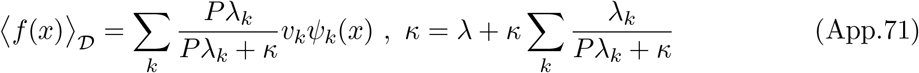

We see that even in the λ → 0 limit, the mean predictor is not equivalent to *y*(*x*) at finite *P*. Rather the sample complexity of each eigenmode *ψ_k_* is set by the evolution of the self-consistent function *κ* which can be nonzero even in the λ → 0 limit. The full generalization error can be decomposed into bias and variance components, where bias component corresponds to the error of the average predictor 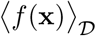.

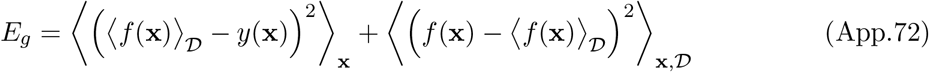

where the bias and variance have the forms

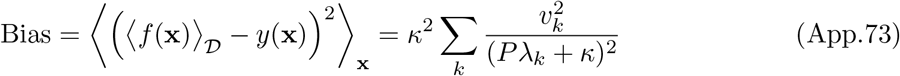

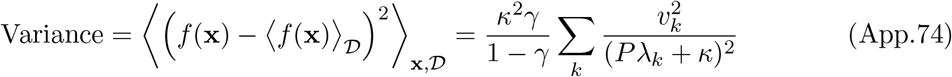

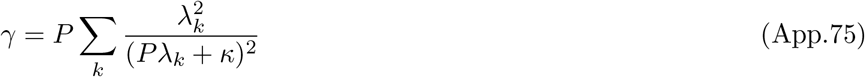

We see that the bias is identical to the generalization error predicted by the equivalent kernel provided that the self-consistent function *κ* is substituted for the ridge parameter λ. This substitution allows for nonvanishing bias in the λ → 0 limit since *κ* > 0 for *P* smaller than the number of nonzero eigenvalues. We find that the sample-to-sample variance of the learned predictor is nonzero 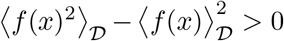 and varies with *P*. This is unlike the Wiener filter theory which treats the variance as 0 for all *P* due to concentration of the training loss.

We note a correspondence between the two theories in the heavily regularized λ → ∞ limit. In this limit, *κ* ~ λ and the bias approaches the risk predicted by the equivalent kernel theory. Further, the variance goes to zero in the λ → ∞ limit since *γ* ~ λ^-2^ → 0. A comparison of the two theories for linear ridge regression on random Gaussian design is provided in Figure App.7.

### 4 Visual Scene Reconstruction Task

#### 4.1 Reconstruction of Natural Scenes from Neural Responses

Using the mouse V1 responses to natural scenes, we attempt to reconstruct original images from the neural codes using different numbers of images. The presented natural scenes are taken from ten classes of imagenet which can be downloaded from https://github.com/MouseLand/stringer-pachitariu-et-al-2018b. Let ***θ***^*μ*^ ∈ ℝ^*D*^ be a *D*-dimensional flattened vector containing the pixel values of the *μ*-th image and let **r**^*μ*^ ∈ ℝ^*N*^ represent the neural response to the *μ*-th image. The goal in the problem is to learn a collection of weights **W** ∈ ℝ^*D*×*N*^ which map neural responses **r**^*μ*^ to images **θ**^*μ*^

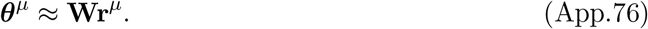

The generalization error *E_g_* again measures the average error on all points, averaged over all possible datasets 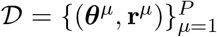 of size *P*. If the optimal weights for dataset 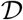 is 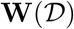 then the generalization error is

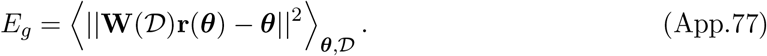

After identifying eigenfunctions *ϕ_k_*(***θ***), we expand the images in this basis ***θ*** = ∑_*k*_ **v**_*k*_*ϕ*_*k*_(***θ***) where **v**_*k*_ ∈ ℝ^*D*^. The generalization error is therefore *E_g_* = ∑_*k*_ |**v**_*k*_|^2^E_*k*_(*P*) and the cumulative power is 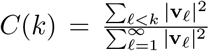. We perform this reconstruction task on many filtered versions of the natural scenes. To construct a filter, we first compute the Fourier transform of the image. Let 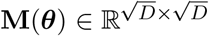 represent the non-flattened image and let 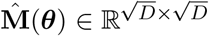 represent the Fourier transform of the image, computed explicitly as

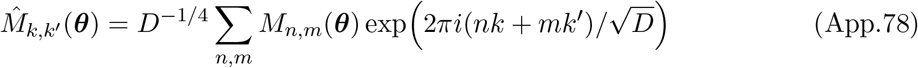

To develop the band-pass filter, we calculate 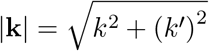 for each of the indices in the matrix. For a band-pass filter with parameters *s_max_, r* we simply zero out the entries in 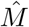 which correspond to states with frequencies outside the appropriate band: for any k, *k′* with 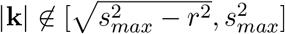 then 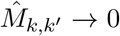. We then perform the inverse Fourier transform on 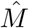 to obtain a filtered version of the original image.

### 5 A Simple Feedforward Model of V1

#### 5.1 Linear Neurons

We consider a simplified but instructive model of the V1 population code as a linear-nonlinear map from photoreceptor responses through Gabor filters and then nonlinearity [17, 18, 19]. Let **x** ∈ ℝ^2^ represent the two-dimensional retinotopic position of photoreceptors. The firing rates of the photoreceptor at position **x** to a static grating stimulus oriented at angle *θ* is

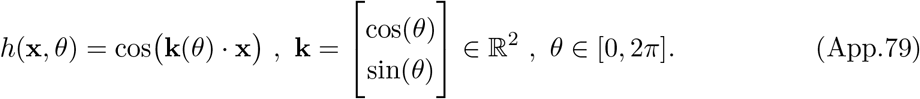

We model each V1 neuron’s receptive field as a Gabor filter of the receptor responses *h*(**x**, *θ*). The *i*-th V1 neuron has preferred wavevector **k**_*i*_, generating the following set of weights between photoreceptors and the *i*-th V1 neuron

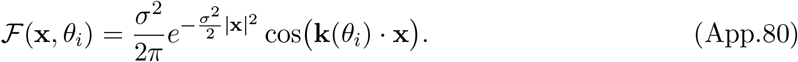

The V1 population code is obtained by filtering the photoreceptor responses. By approximating the resulting sum over all retinal photoreceptors with an integral, we find the response of neuron *i* to grating stimulus with wavenumber **k** is

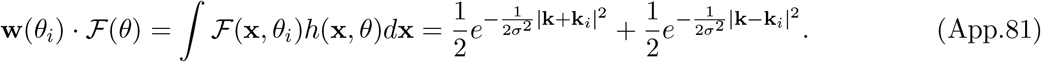

The response of neuron *i* is computed through nonlinear rectification of this input current *r_i_*(*θ*) = *g*(**w**(*θ_i_*) · *h*(*θ*)). For a linear neuron *g*(*z*) = *z*, the kernel has the following form

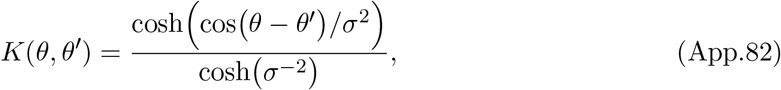

where the kernel is normalized to have maximum value of 1. Note that this normalization of the kernel is completely legitimate since it merely rescales each eigenvalue by a constant and does not change the learning curves.

Since the kernel only depends on the difference between angles *θ* – *θ′,* it is said to posess translation invariance. Such translation invariant kernels admit a Mercer decomposition in terms of Fourier modes *K*(*θ*) = ∑_*n*_ λ_*n*_ cos(*nθ*) since the Fourier modes diagonalize shift invariant integral operators on 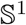. For the linear neuron, the kernel eigenvalues scale like 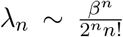, indicating infinite differentiability of the tuning curves. Since *λ_n_* decays rapidly with *n*, we find that this Gabor code has an inductive bias that favors low frequency functions of orientation *θ*.

#### 5.2 Nonlinear Simple Cells

Introducing nonlinear functions *g*(*z*) that map input currents *z* into the V1 population into firing rates, we can obtain a non-linear kernel *K_g_*(*θ*) which has the following definition

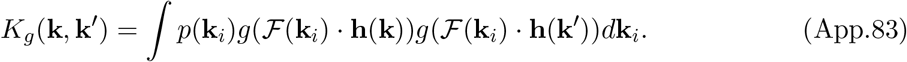

In this setting, it is convenient to restrict **k**_*i*_, **k**, 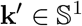 and assume that the preferred wavevectors **k**_*i*_ are uniformly distributed over the circle. In this case, it suffices to identify a decomposition of the composed function *g*(**w**_*i*_ · **h**(*θ*)) in the basis of Chebyshev polynomials *T_n_*(*z*) which satisfy *T_n_* (cos(*θ*)) = cos(*nθ*)

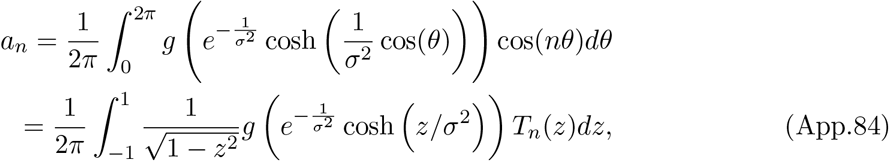

which can be computed efficiently with an appropriate quadrature scheme. Once the coefficients *a_n_* are determined, we can compute the kernel by first letting *θ_i_* to be the angle between **k** and **k**_*i*_ and letting *θ* be the angle between **k** and **k**′

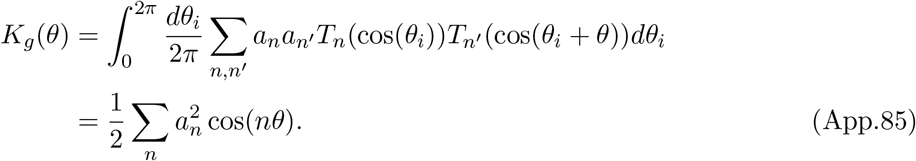

Thus the kernel eigenvalues are 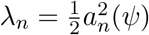.

##### Asymptotic scaling of spectra

Activation functions that encourage sparsity have slower eigenvalue decays. If the nonlinear activation function has the form *g_q,t_*(*z*) = max{0,*z* ‒ *a*}^*q*^, then the spectrum decays like *λ_n_* ~ *n*^-2*q*-2^. A simple argument justifies this scaling: if the function *g*(*e*^-*σ*2^ cosh(*σ*^2^*z*)) is only *q* – 1 times differentiable then *a_n_n^q^* ~ *n*^-1^ since ∑_*n*_ *a_n_n^q^* must diverge. Therefore 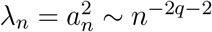. Note that this scaling is independent of the threshold.

#### 5.3 Phase Variation, Complex Cells and Invariance

We can consider a slightly more complicated model where Gabors and stimuli have phase shifts

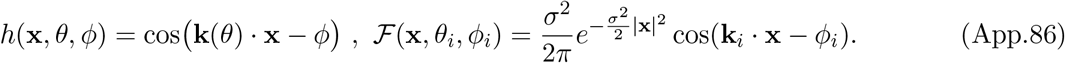

The simple cells are generated by nonlinearity

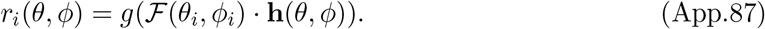

The input currents into the simple V1 cells can be computed exactly

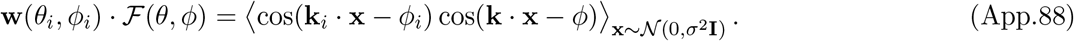

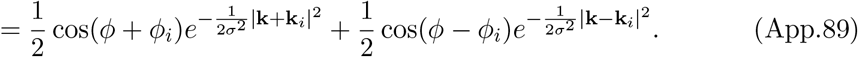

When |**k**| = |**k**_*i*_| = 1, the simple cell tuning curves *r_i_* = *g*(**w**_*i*_ · **h**) only depend on cos(*θ* – *θ_i_*) and *ϕ*, allowing a Fourier decomposition

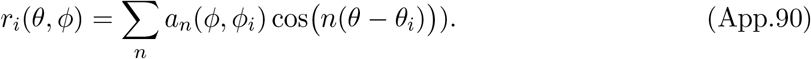

The simple cell kernel *K_s_,* therefore decomposes into Fourier modes over *θ*

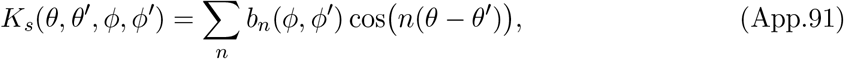

where 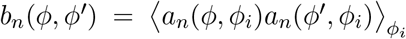. It therefore suffices to solve the infinite sequence of integral eigenvalue problems over *ϕ*

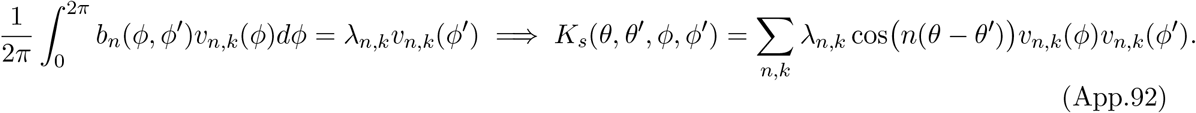

With this choice it is straightforward to verify that the kernel eigenfunctions are *v_n,k_*(*θ, *ϕ**) = *e*^*inθ*^*v_n,k_*(*ϕ*) with corresponding eigenvalue λ_*n,k*_ Since *b_n_* is not translation invariant in *ϕ* – *ϕ′,* the eigenfunctions *v_n,k_* are not necessarily Fourier modes. These eigenvalue problems for *b_n_* must be solved numerically when using arbitrary nonlinearity *ψ*. The top eigenfunctions of the simple cell kernel depend heavily on the phase of the two grating stimuli *ϕ*. Thus, a pure orientation discrimination task which is independent of phase requires a large number of samples to learn with the simple cell population.

#### 5.4 Complex Cells Populations are Phase Invariant

V1 also contains complex cells which possess invariance to the phase *ϕ* of the stimulus. Again using Gabor filters

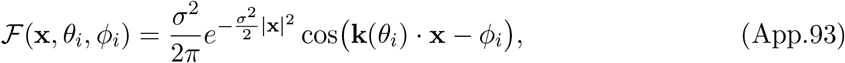

we model the complex cell responses with a quadratic nonlinearity and sum over two squared filters which are phase shifted by *π*/2

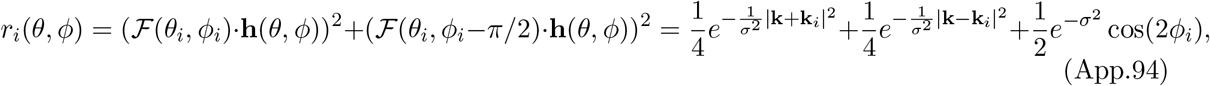

which we see is independent of the phase *ϕ* of the grating stimuluso Integrating over the set of possible Gabor filters (**k**_*i*_, *ϕ_i_*) again gives the following kernel for the complex cells

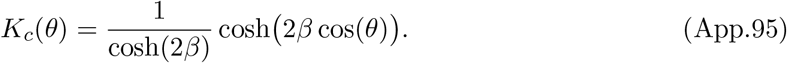

Remarkably, this kernel is independent of the phase *ϕ* of the grating stimuluso Thus, complex cell populations possess good inductive bias for vision tasks where the target function only depends on the orientation of the stimulus rather than it’s phaseo In reality, V1 is a mixture of simple and complex cellso Let *s* ∈ [0,1] represent the relative proportion of neurons which are simple cells and (1 – *s*) the relative proportion of complex cellso The kernel for the mixed V1 population is given by a simple convex combination of the simple and complex cell kernels

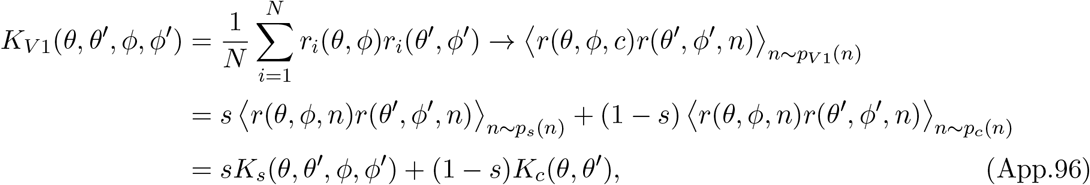

where *n* denotes neuron type (simple vs complex, tuning etc) and *P*_*V*1_(*n*),*p_s_*(*n*),*p_c_*(*n*) are probability distributions over the V1 neuron identities, the simple cell identities and the complex cell identities respectively. Increasing *s* increases the phase dependence of the code by giving greater weight to the simple cell population. Decreasing *s* gives weight to the complex cell population, encouraging phase invariance of readouts.

#### 5.5 Visualization of Feedforward Gabor V1 Model and Induced Kernels

Examples of the induced kernels for the Gabor-bank V1 model are provided in Figure App.4. We show how choice of rectifying nonlinearity *g*(*z*) and sparsifying threshold *a* influence the kernel and their spectra. Learning curves for simple orientation tasks are provided.

### 6 Time Dependent Neural Codes

#### 6.1 RNN Model and Decomposition

In this setting, the population code **r**({***θ***(*t*)},*t*) is a function of an input stimulus sequence ***θ***(*t*) and time *t*. In general the neural code **r** at time *t* can depend on the entire history of the stimulus input ***θ***(*t*′) for *t*′ ≤ *t*, as is the case for recurrent neural networks. We denote dependence of a function *f* on ***θ***(*t*) in this causal manner with the notation *f*({***θ***},*t*). In a learning task, a set of readout weights **w** are chosen so that a downstream linear readout *f*({***θ***},*t*) = **w** · **r**({***θ***},*t*) approximates a target sequence *y*({***θ***},*t*) which maps input stimulus sequences to output scalar sequences. The quantity of interest is the generalization *E_g_*, which in this case is an average over both input sequences and time, *E_g_* = 〈(*y*({***θ***},*t*) – *f*({***θ***},*t*)〉^2^)_***θ***(*t*)*t*_. The average is computed over a distribution of input stimulus sequences *p*(***θ***(*t*)). To train the readout, **w**, the network is given a sample of *P* stimulus sequences ***θ***^*μ*^(*t*), *μ* = 1,…, *P*. For the *μ*-th training input sequence, the target system *y* is evaluated at a set of discrete time points 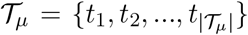 giving a collection of target values 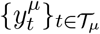 and a total dataset of size 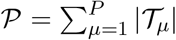. The *average case generalization* computes a further average of the generalization error *E_g_* over randomly sampled datasets of size 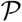.

Learning is again achieved through iterated weight updates with delta-rule form, but now have contributions from both sequence index and time 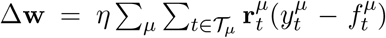. As before, optimization of the readout weights is equivalent to kernel regression with a kernel that computes inner products of neural population vectors at different times *t,t*′ for different input sequences 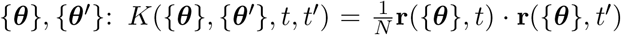. This kernel depends on details of the time varying population code including its recurrent intrinsic dynamics as well as its encoding of the time-varying input stimuli. The optimization problem and delta rule described above converge to the kernel regression solution for kernel gram matrix 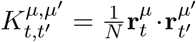 [20, 21, 22]. The learned function has the form 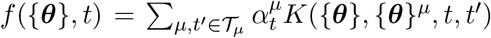, where ***α*** = **K**^+^**y** for kernel gram matrix 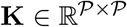 which is computed for the entire set of training sequences, and the vector 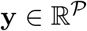 is the vector containing the desired target outputs for each sequence. Assuming a probability distribution over sequences ***θ***(*t*), the kernel can be diagonalized with orthonormal eigenfunctions *ψ_k_*({***θ***},*t*). Our theory carries over from the static case: kernels whose top eigenfunctions have high alignment with the target dynamical system *y*({***θ***}, *t*) will achieve the best average case generalization performance.

## Appendix Figures

**Figure App.1:**
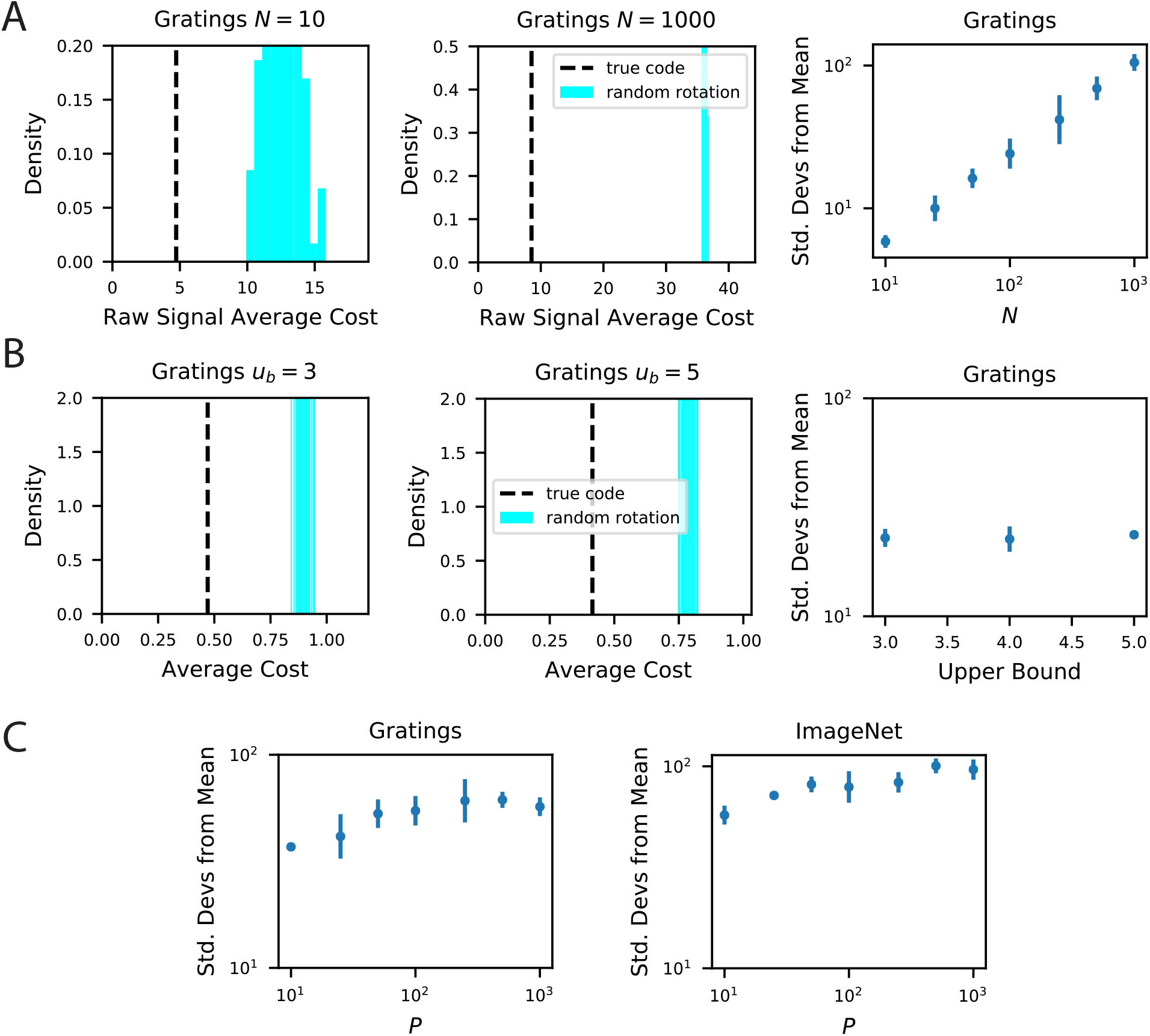
Our metabolic efficiency finding is robust to different pre-processing techniques and upper bounds on neural firing. **A** We show the same result as the main text except we use raw (non *z*-scored) estimate of responses for each stimulus. **B** Our result is robust to imposition of firing rate upper bounds *u_b_* on each neuron. This result uses the *z*-scored responses to be consistent with the rest of the paper. The biological code achieves a maximum *z*-score values in the range [3.2, 4.7], which motivated the range of our tested upper bound values {3, 4, 5}. **C** Our finding is robust to the number of sampled stimuli *P* as we show in an experiment where rotations in *N* = 500 dimensional subspace.

**Figure App.2:**
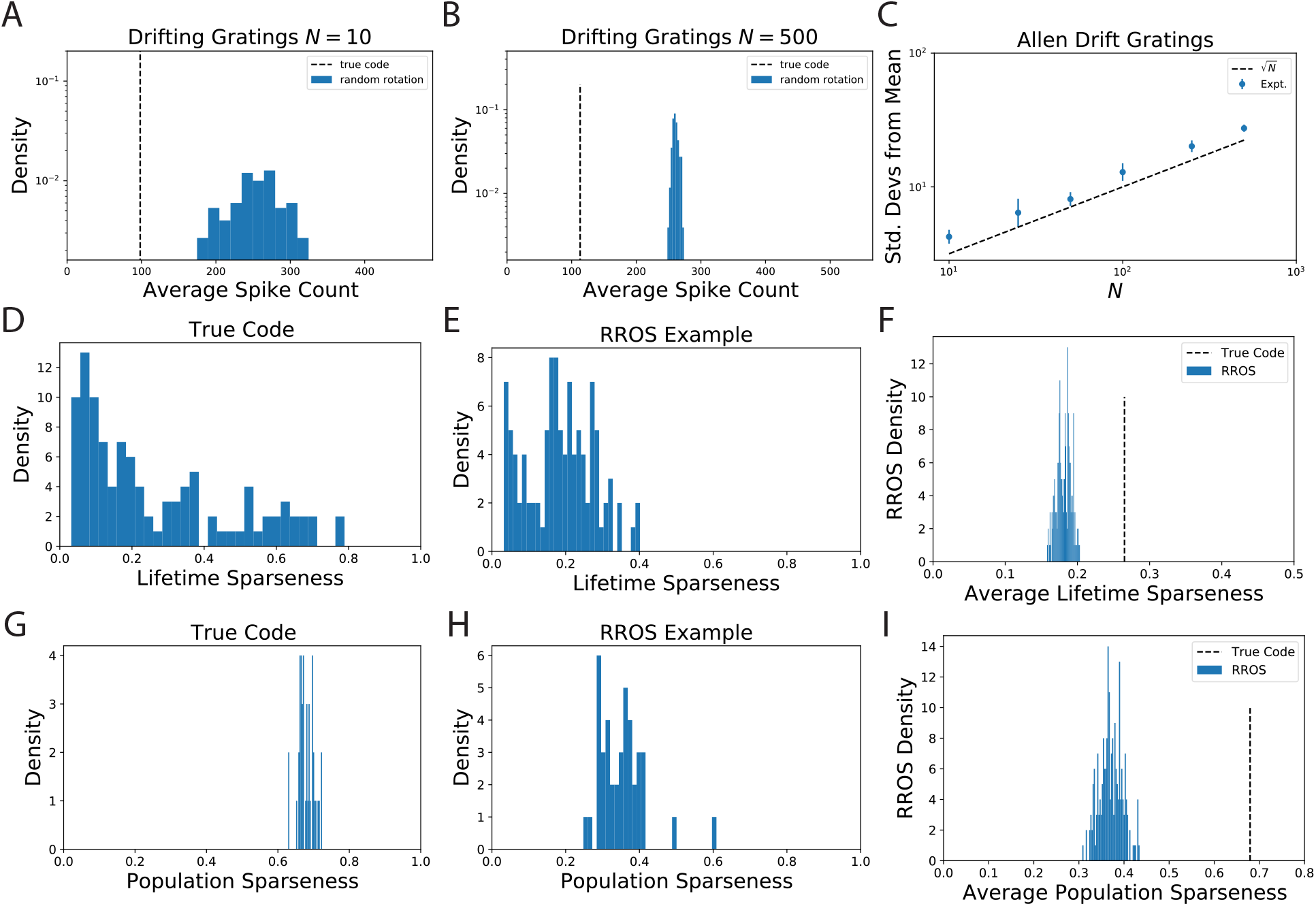
The observation that randomly oriented codes with the same kernel require higher spike counts than the original code is reproduced from electrophysiological recordings of Mouse visual cortex (VISp and VISal) from the AIBO. **A** Distribution of average spike count for a random selection of *N* = 10 neurons. **B** The same for *N* = 500 randomly selected neurons. **C** Since the distribution of average spike count for the randomly rotated codes concentrates with *N* the number of standard deviations the true code is from the mean increases with N. We show that this scaling is approximately like 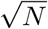 which suggests that the variability in average spike count for randomly rotated codes goes scales with neuron count like 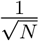. This is intuitive since the average spike count over the rotated code is an empirical average over *N* random variables. **D** The lifetime sparseness of neurons in the true code are spread out over a large range. **E** The lifetime sparseness distribution for an example RROS code does not have the same range. **F** The average (over neurons) lifetime sparseness of RROS codes over random rotations is significantly lower than the average lifetime sparseness of the true code. **G**-**I** The true visual cortex code has much higher population sparseness over each grating stimulus as well

**Figure App.3:**
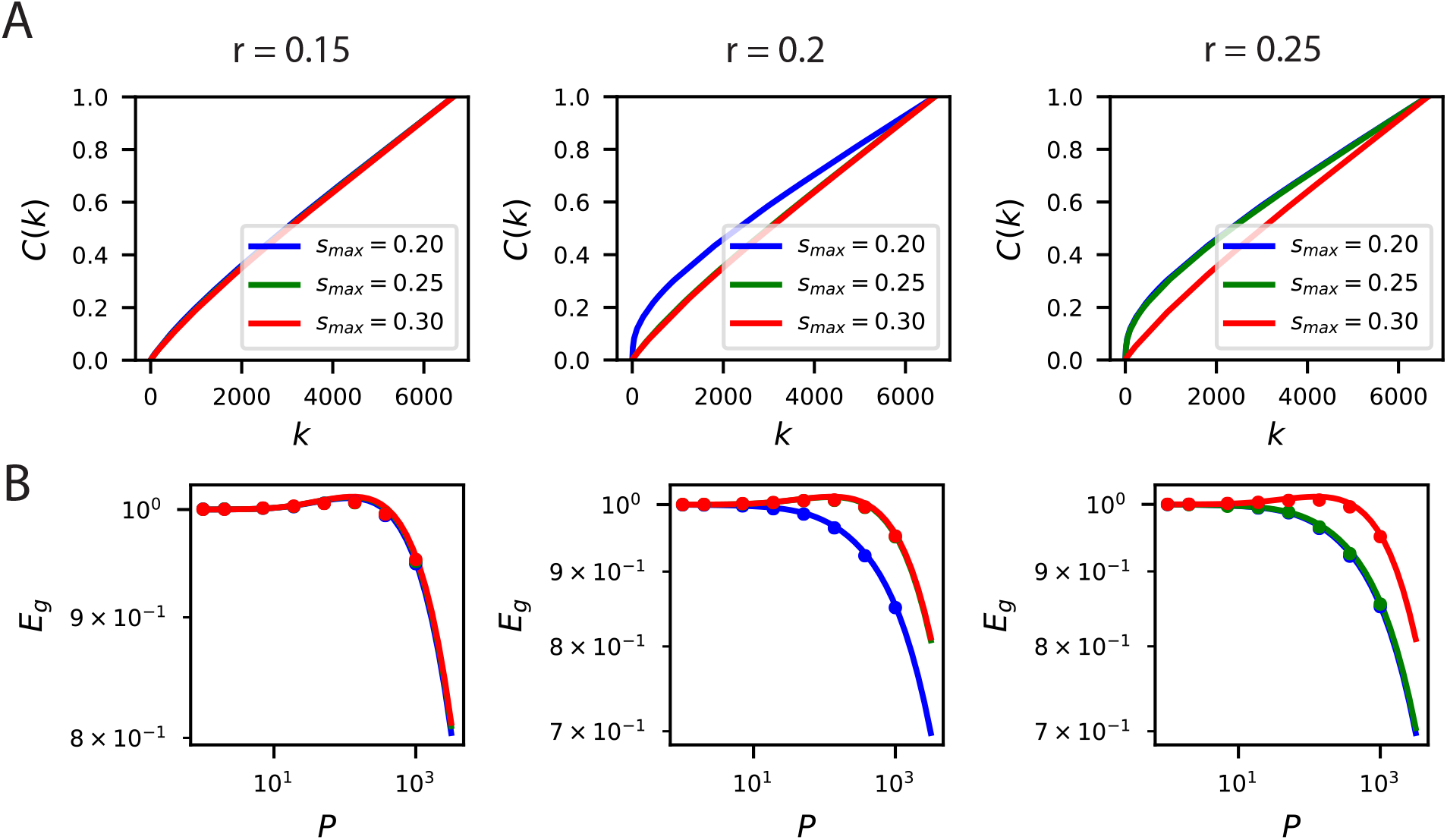
The inductive bias to reconstruct low spatial frequency components of natural scenes from the population responses holds over several band-pass filters of the form 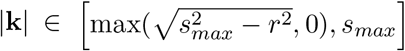. **A** The cumulative power curves for reconstruction of the filtered images with different values of *c* and *s_max_*. Lower values of *s_max_* and higher values of *c* preserve mostly low frequency content in the images and are easier to reconstruct from the neural responses. **B** The learning curves respect the ordering of the cumulative power metric *C*(*k*).

**Figure App.4:**
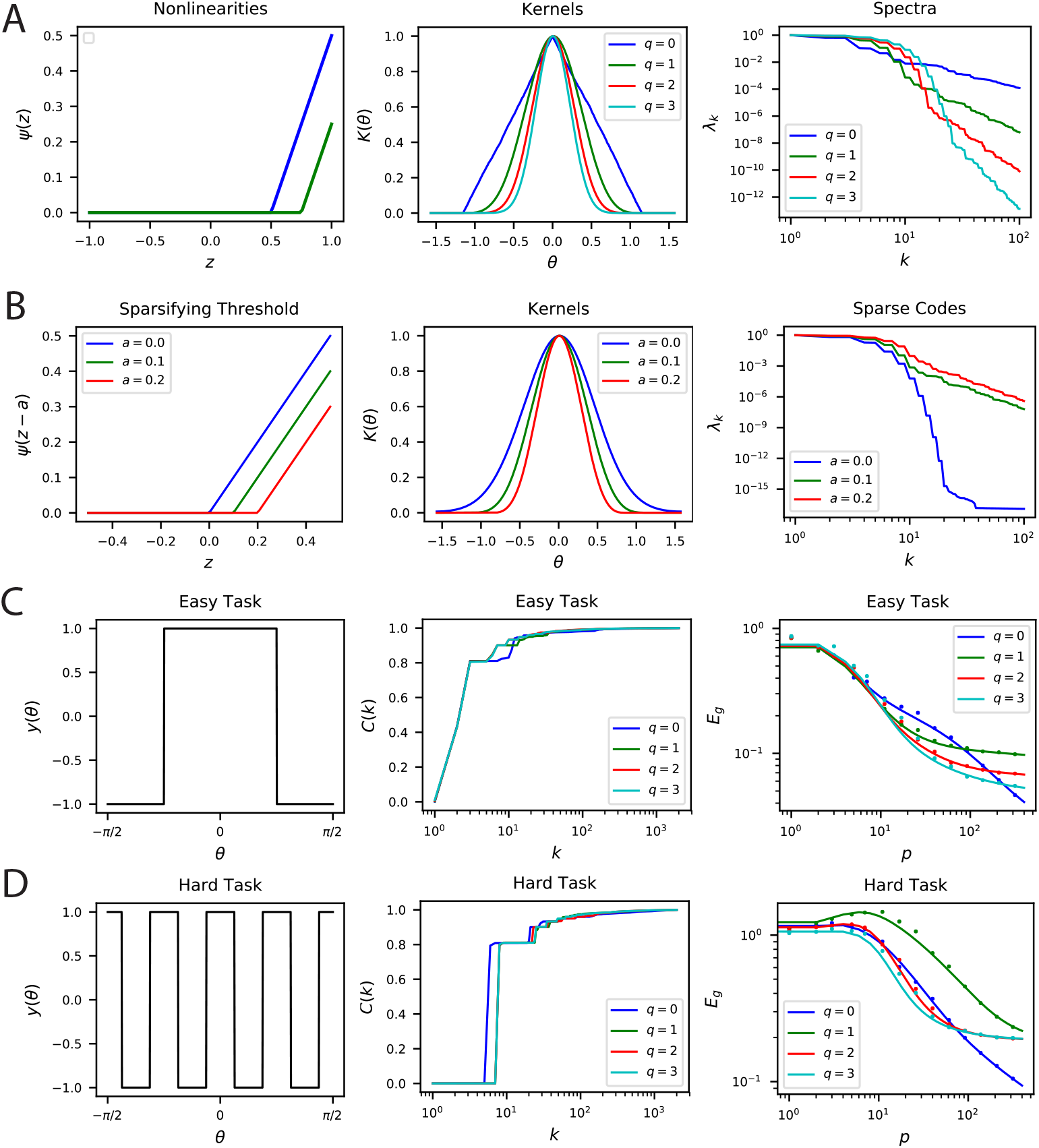
Nonlinear Rectification and proportion of simple and complex cells influences the inductive bias of the population code. **A** The choice of nonlinearity has influence on the kernel and its spectrum. If the nonlinearity is *g*(*z*) = max(0,*z^q^*), then λ_*k*_ ~ *k*^-2*n*-2^. **B** The sparsity can be increased by shifting the nonlinearity *g*(*z*) → *g*(*z* – *a*). Sparser codes have higher dimensionality. Note that *a* = 0 is a special case where the neurons behave in the linear regime for all inputs *θ* since the currents **w** · **h** are positive. Thus, for *a* = 0, the spectrum decays like a Bessel Function *λ_k_* = *I_k_* (*β*). **C-D** Easy and hard orientation discrimination tasks with varying nonlinear polynomial order *q*. At low sample sizes, large *q* performs better, whereas at large *P*, the step function nonlinearity *q* = 0 achieves the best performance.

**Figure App.5:**
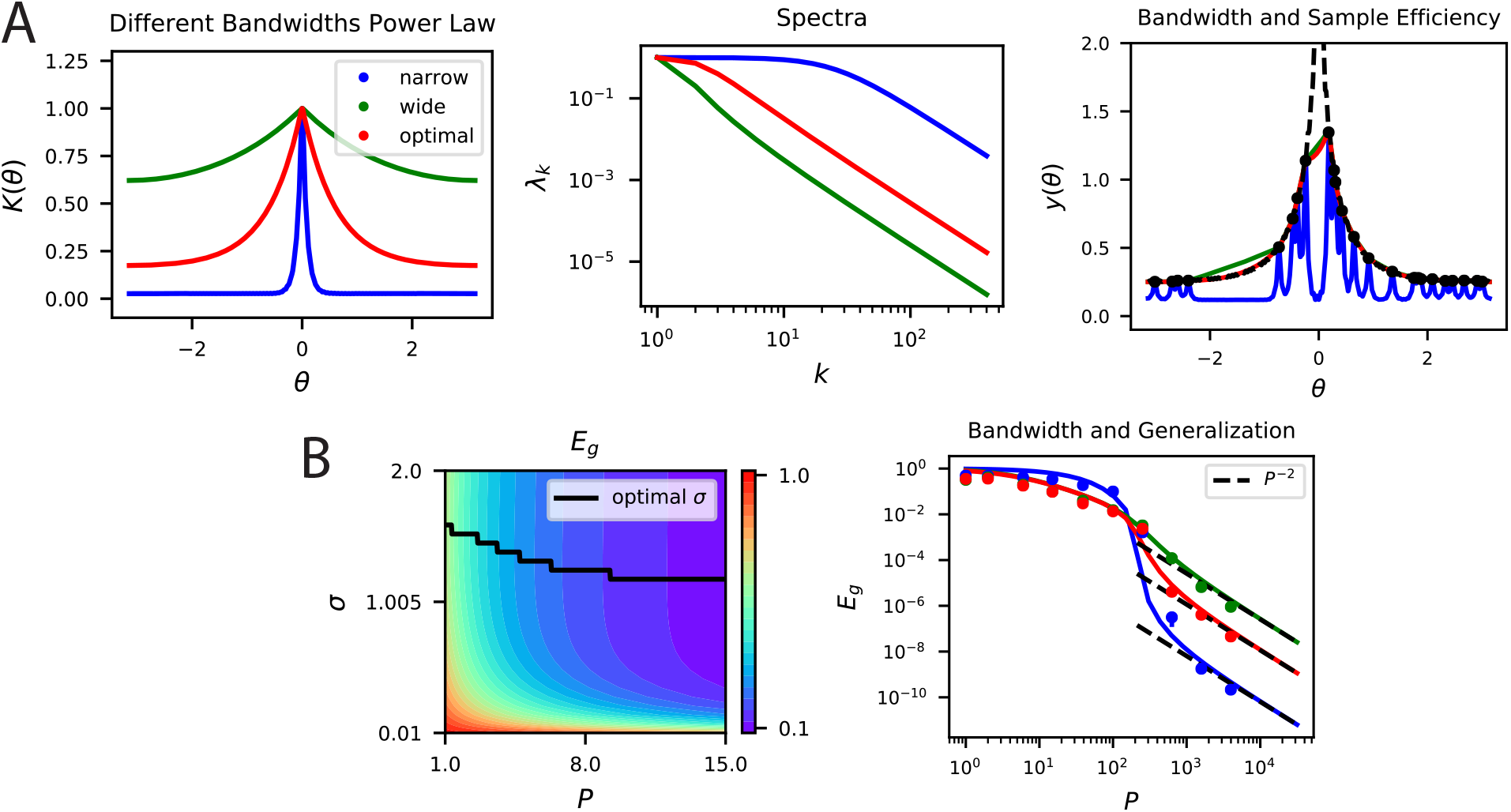
**A, B** Kernel regression experiments are performed with Laplace kernels of varying bandwidth on a non-differentiable target function. The top eigenvalues are modified by changing the bandwidth, but the asymptotic power law scaling is preserved. Generalization at low *P* is shown in the contour plot while the large *P* scaling is provided in the generalization. In A-right and B-right, color code is the same as in the main text.

**Figure App.6:**
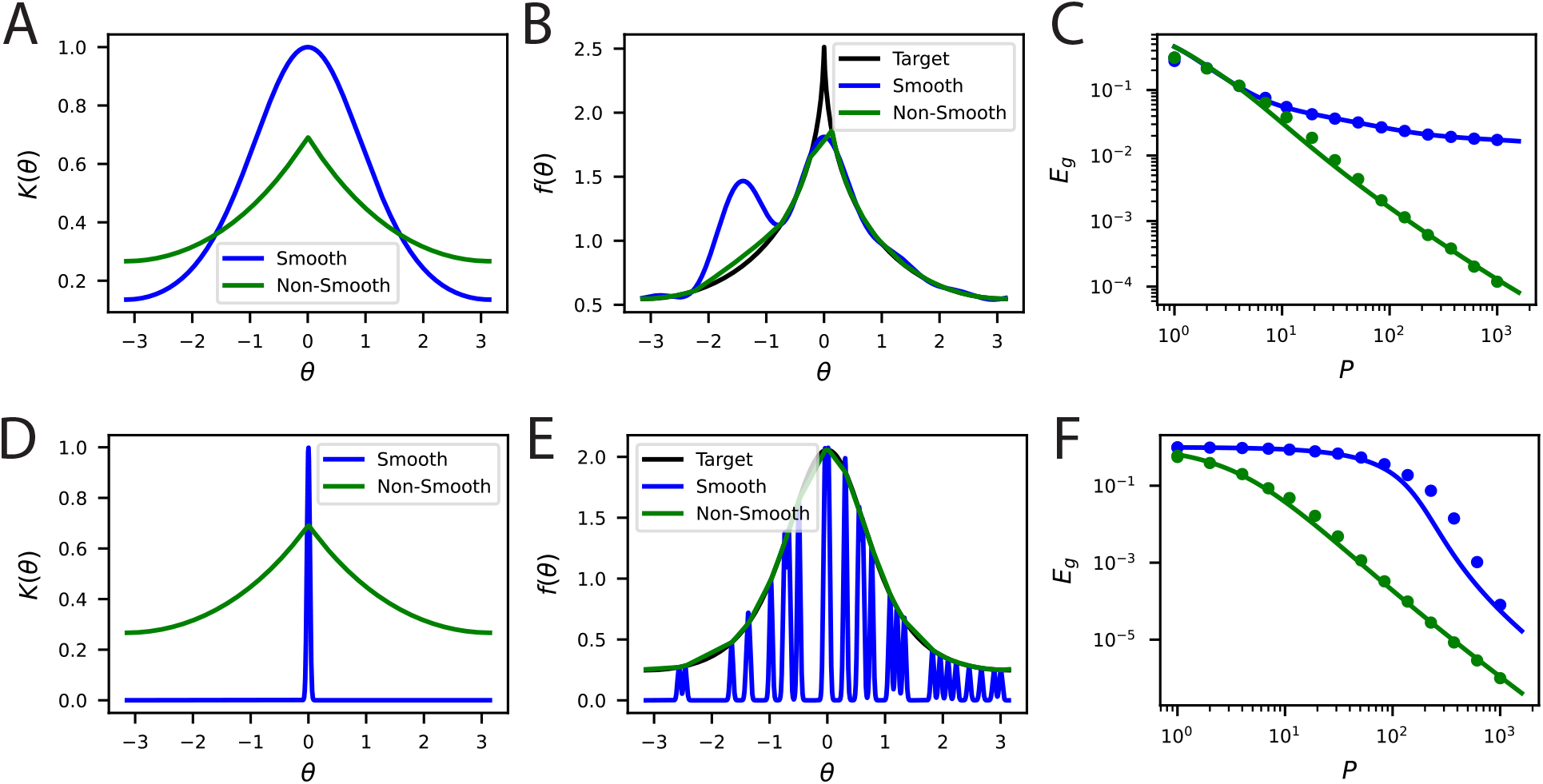
Non-differentiable kernels can generalize better than infinitely differentiable kernels in a variety of contexts. **A** An infinitely differentiable von-Mises kernel (blue) will be compared to a non-differentiable Laplace kernel (green). **B** For a non-differentiable target function *y*(***θ***) with a cusp, the non-smooth kernel can provide a better fit to the target function at *P* = 30 samples. **C** The learning curves for this task. Solid lines are theory, circles are simulations. **D** The lengthscale of the kernel can be more important than local smoothness. We will now compare a narrow von-Mises kernel (still infinitely differentiable) with a non-differentiable wide kernel. **E** The non-smooth kernel generalizes on a smooth task almost perfectly whereas the narrow smooth kernel only locally interpolates. **F** The learning curves for this smooth task.

**Figure App.7:**
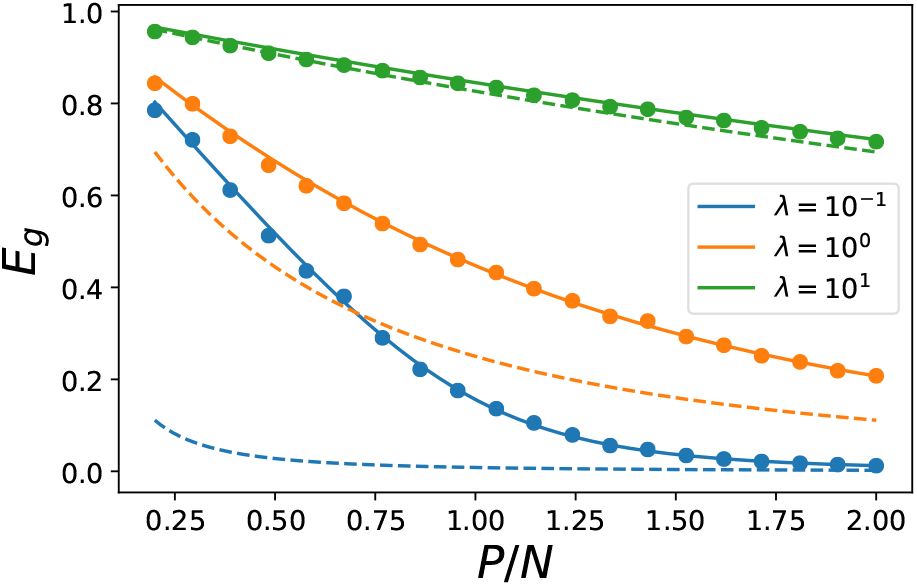
Our theoretical prediction for generalization error *E_g_* (solid lines) on a linear kernel *K*(**x**, *x*′) = **x** · ***x***′ regression problem with uncorrelated Gaussian design 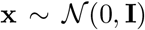 and linear teacher *y* = *β* · **x** in *N* = 500 dimensions. The generalization error as a function of sample size *P* is plotted predicted by the equivalent kernel/Wiener filter (dashed). Each color represents a different regularization level λ. The experimental generalization is shown as dots of the corresponding color and shows the numerical error obtained from solving the kernel regression problem with regularization level λ. We see that the two theories coincide at large λ but are drastically different at low levels of explicit regularization λ.

