## Supplementary Information for "Population codes enable learning from few examples by shaping inductive bias"

#### 1 Neural Code Decomposition

##### 1.1 Singular Value Decomposition of Continuous Population Responses

SVD of population responses is usually evaluated with respect to a discrete and finite set of stimuli. In the main paper, we implicitly assumed that a generalization of SVD to a continuum of stimuli. In this section we provide an explicit construction of this generalized SVD using techniques from functional analysis. Our construction is an example of the quasimatrix SVD defined in [1] and justifies our use of SVD in the main text.

For our construction, we note that Mercer's theorem guarantees the existence of an eigendecomposition of any inner product kernel $K(\boldsymbol{\theta}, \boldsymbol{\theta}')$ in terms of a complete orthonormal set of functions $\{\psi_k\}_{k=1}^{\infty}$ [2]. In particular, there exist a non-negative (but possibly zero) summable eigenvalues $\{\lambda_k\}_{k=1}^{\infty}$ and a corresponding set of orthonormal eigenfunctions such that

$$K(\boldsymbol{\theta}, \boldsymbol{\theta}') = \sum_{k=1}^{\infty} \lambda_k \psi_k(\boldsymbol{\theta}) \psi_k(\boldsymbol{\theta}'). \tag{App.1}$$

For a stimulus distribution $p(\boldsymbol{\theta})$, the set of functions $\{\psi_k\}_{k=1}^{\infty}$ are orthonormal and form a complete basis for square integrable functions $L_2$ which means

$$\begin{aligned} \left\langle \psi_k(\boldsymbol{\theta}) \psi_\ell(\boldsymbol{\theta}) \right\rangle_{\boldsymbol{\theta}} &= \int p(\boldsymbol{\theta}) \psi_k(\boldsymbol{\theta}) \psi_\ell(\boldsymbol{\theta}) d\boldsymbol{\theta} = \delta_{k\ell}, \\ f(\boldsymbol{\theta}) &= \sum_k \left\langle f(\boldsymbol{\theta}') \psi_k(\boldsymbol{\theta}') \right\rangle_{\boldsymbol{\theta}'} \psi_k(\boldsymbol{\theta}) \ , \ \forall f \in L_2. \end{aligned} \tag{App.2}$$

Next, we use this basis to construct the SVD. Each of the tuning curves $r_i \in L_2$ (assumed to be square integrable) can be expressed in this basis with the top $N$ of the functions in the set $\{\psi_k\}_{k=1}^{\infty}$

$$r_i(\boldsymbol{\theta}) = \sum_{k=1}^{N} A_{ik} \psi_k(\boldsymbol{\theta}), \tag{App.3}$$

where we introduced a matrix $\mathbf{A} \in \mathbb{R}^{N \times N}$ of expansion coefficients. Note that $\text{rank}(\mathbf{A}) \leq N$. We compute the singular value decomposition of the finite matrix $\mathbf{A}$

$$\mathbf{A} = \sqrt{N} \sum_{k=1}^{\text{rank}(\mathbf{A})} \sqrt{\lambda_k} \mathbf{u}_k \mathbf{v}_k^\top. \tag{App.4}$$

We note that the signal correlation matrix for this population code can be computed in closed form

$$\mathbf{\Sigma}_s = \frac{1}{N} \mathbf{A} \left\langle \psi(\boldsymbol{\theta}) \psi(\boldsymbol{\theta})^\top \right\rangle_{\boldsymbol{\theta}} \mathbf{A}^\top = \frac{1}{N} \mathbf{A}\mathbf{A}^\top = \sum_{k=1}^{\text{rank}(\mathbf{A})} \lambda_k \mathbf{u}_k \mathbf{u}_k^\top, \tag{App.5}$$

due to the orthonormality of $\{\psi_k\}$. Thus the principal axes $\mathbf{u}_k$ of the neural correlations are the left singular vectors of $\mathbf{A}$. We may similarly express the inner product kernel in terms of the eigenfunctions

$$K(\boldsymbol{\theta}, \boldsymbol{\theta}') = \frac{1}{N} \mathbf{r}(\boldsymbol{\theta}) \cdot \mathbf{r}(\boldsymbol{\theta}') = \frac{1}{N} \psi(\boldsymbol{\theta})^\top \mathbf{A}^\top \mathbf{A} \psi(\boldsymbol{\theta}'). \tag{App.6}$$

The kernel eigenvalue problem demands [2]

$$\int p(\boldsymbol{\theta}) K(\boldsymbol{\theta}, \boldsymbol{\theta}') \psi(\boldsymbol{\theta}) d\boldsymbol{\theta} = \frac{1}{N} \mathbf{A}^\top \mathbf{A} \psi(\boldsymbol{\theta}') = \mathbf{\Lambda} \psi(\boldsymbol{\theta}') \implies \frac{1}{N} \mathbf{A}^\top \mathbf{A} = \mathbf{\Lambda}$$
$$\implies \sum_{k=1}^{\text{rank}(\mathbf{A})} \lambda_k \mathbf{v}_k \mathbf{v}_k^\top = \sum_{k=1}^{\text{rank}(\mathbf{A})} \lambda_k \mathbf{e}_k \mathbf{e}_k^\top. \tag{App.7}$$

The $\mathbf{v}_k$ vectors must be identical to $\pm \mathbf{e}_k$, the Cartesian unit vectors, if the eigenvalues are non-degenerate. From this exercise, we find that the SVD for $\mathbf{A}$ has the form $\mathbf{A} = \sqrt{N} \sum_{k=1}^{\text{rank}(\mathbf{A})} \sqrt{\lambda_k} \mathbf{u}_k \mathbf{e}_k^\top$. With this choice, the population code admits a singular value decomposition

$$\mathbf{r}(\boldsymbol{\theta}) = \mathbf{A} \psi(\boldsymbol{\theta}) = \sqrt{N} \sum_{k=1}^{\text{rank}(\mathbf{A})} \sqrt{\lambda_k} \mathbf{u}_k \psi_k(\boldsymbol{\theta}). \tag{App.8}$$

This singular value decomposition demonstrates the connection between neural manifold structure (principal axes $\mathbf{u}_k$) and function approximation (kernel eigenfunctions $\psi_k$). This singular value decomposition can be verified by computing the inner product kernel and the correlation matrix, utilizing the orthonormality of $\{\mathbf{u}_k\}$ and $\{\psi_k\}$. This exercise has important consequences for the space of learnable functions, which is at most $\text{rank}(\mathbf{A})$ dimensional since linear readouts lie in $\text{span}\{r_i(\boldsymbol{\theta})\}_{i=1}^N$.

#### 1.2 Discrete Stimulus Spaces: Finding Eigenfunctions with Matrix Eigendecomposition

In our discussion so far, our notation suggested that $\boldsymbol{\theta}$ take a continuum of values. Here we want to point that our theory still applies if $\boldsymbol{\theta}$ take a discrete set of values. In this case, we can think of a Dirac measure $p(\boldsymbol{\theta}) = \sum_{i=1}^{\tilde{P}} p_i \delta(\boldsymbol{\theta} - \boldsymbol{\theta}^i)$, where $i$ indexes all the $\tilde{P}$ values $\boldsymbol{\theta}$ can take. With this choice

$$\int p(\boldsymbol{\theta})K(\boldsymbol{\theta},\boldsymbol{\theta}')\psi_k(\boldsymbol{\theta})d\boldsymbol{\theta} = \sum_{i=1}^{\tilde{P}} p_i K(\boldsymbol{\theta}^i,\boldsymbol{\theta}')\psi_k(\boldsymbol{\theta}^i) = \lambda_k\psi_k(\boldsymbol{\theta}'). \tag{App.9}$$

Demanding this equality for $\boldsymbol{\theta}' = \boldsymbol{\theta}^i,\ i = 1, ..., \tilde{P}$ generates a matrix eigenvalue problem

$$\mathbf{KB\Psi} = \mathbf{\Psi\Lambda}, \tag{App.10}$$

where $\mathbf{B}_{ij} = \delta_{ij}p_i$. The eigenfunctions over the stimuli are identified as the columns of $\mathbf{\Psi}$ while the eigenvalues are the diagonal elements of $\mathbf{\Lambda}_{k\ell} = \lambda_k\delta_{k\ell}$.

*Experimental considerations:* In an experimental setting, a finite number of stimuli are presented and the SVD is calculated over this finite set regardless of the support of $p(\boldsymbol{\theta})$. This raises the question of the interpretation of this SVD and its relation to the inductive bias theory we presented. Here we provide two interpretations.

In the first interpretation, we think of the empirical SVD as providing an estimate of the SVD over the full distribution $p(\boldsymbol{\theta})$. To formalize this notion, we can introduce a Monte-Carlo estimate of the integral eigenvalue problem

$$\int p(\boldsymbol{\theta})K(\boldsymbol{\theta},\boldsymbol{\theta}')\psi_k(\boldsymbol{\theta})d\boldsymbol{\theta} \approx \frac{1}{\tilde{P}}\sum_{\mu=1}^{\tilde{P}} K(\boldsymbol{\theta}^\mu,\boldsymbol{\theta}')\psi_k(\boldsymbol{\theta}^\mu) = \lambda_k\psi_k(\boldsymbol{\theta}'). \tag{App.11}$$

For this interpretation to work, the experimenter must sample the stimuli from $p(\boldsymbol{\theta})$, which could be the natural stimulus distribution. Measuring responses to a larger number of stimuli gives a more accurate approximation of the integral above, which will provide a better estimate of generalization performance on the true distribution $p(\boldsymbol{\theta})$.

In the second interpretation, we construct an empirical measure on $\tilde{P}$ experimental stimulus values $\hat{p}(\boldsymbol{\theta}) = \frac{1}{\tilde{P}}\sum_{\mu=1}^{\tilde{P}}\delta(\boldsymbol{\theta}-\boldsymbol{\theta}^\mu)$, and consider learning and generalization over this distribution. This allows the application of our theory to an experimental setting where $\hat{p}(\boldsymbol{\theta})$ is designed by an experimenter. For example, the experimenter could procure a complicated set of $\tilde{P}$ videos, to which an associated function $y(\theta)$ must be learned. After showing these videos to the animal and measuring neural responses, the experimenter could compute, with our theory, generalization error for a uniform distribution over this full set of $\tilde{P}$ videos. Our theory would predict generalization over this distribution after providing supervisory feedback for only a strict subset of $P < \tilde{P}$ videos. Under this interpretation, the relationship between the integral eigenvalue problem and matrix eigenvalue problem is exact rather than approximate

$$\int \hat{p}(\boldsymbol{\theta})K(\boldsymbol{\theta},\boldsymbol{\theta}')\psi_k(\boldsymbol{\theta})d\boldsymbol{\theta} = \frac{1}{\tilde{P}}\sum_{\mu=1}^{\tilde{P}} K(\boldsymbol{\theta}^\mu,\boldsymbol{\theta}')\psi_k(\boldsymbol{\theta}^\mu) = \lambda_k\psi_k(\boldsymbol{\theta}'). \tag{App.12}$$

Demanding either of (App.11) or (App.12) equalities for $\boldsymbol{\theta}' = \boldsymbol{\theta}^\nu,\ \nu = 1, ..., P$ generates a matrix eigenvalue problem

$$\mathbf{K\Psi} = P\mathbf{\Psi\Lambda}. \quad \text{(App.13)}$$

The eigenfunctions restricted to $\{\boldsymbol{\theta}^{\mu}\}$ are identified as the columns of $\mathbf{\Psi}$ while the eigenvalues are the diagonal elements of $\mathbf{\Lambda}_{k\ell} = \lambda_k \delta_{k\ell}$. For the case where $N$ and $P$ are finite, the spectrum obtained through eigendecomposition of the kernel $\mathbf{K}$ is the same as would be obtained through the finite $N$ signal correlation matrix $\mathbf{\Sigma}_s$, since they are inner and outer products of trial averaged population response matrices $\mathbf{R}$.

#### 1.3 Translation Invariant Kernels

For the special case where the data distribution $p(\boldsymbol{\theta}) = \frac{1}{V}$ is uniform over volume $V$ and the kernel is translation invariant $K(\boldsymbol{\theta}, \boldsymbol{\theta}') = \kappa(\boldsymbol{\theta} - \boldsymbol{\theta}')$, the kernel can be diagonalized in the basis of plane waves

$$\int p(\boldsymbol{\theta}) K(\boldsymbol{\theta}, \boldsymbol{\theta}') \psi_{\mathbf{k}}(\boldsymbol{\theta}) d\boldsymbol{\theta} = \frac{1}{V} \int \kappa(\boldsymbol{\theta} - \boldsymbol{\theta}') e^{i\mathbf{k}\cdot\boldsymbol{\theta}} d\boldsymbol{\theta} = \frac{1}{V} \hat{\kappa}(\mathbf{k}) e^{i\mathbf{k}\cdot\boldsymbol{\theta}'} \quad \text{(App.14)}$$

The eigenvalues are the Fourier components of the Kernel $\lambda_{\mathbf{k}} = \frac{1}{V}\hat{\kappa}(\mathbf{k}) = \frac{1}{V}\int d\boldsymbol{\theta} e^{i\mathbf{k}\cdot\boldsymbol{\theta}} \kappa(\boldsymbol{\theta})$ while the eigenfunctions are plane waves $\psi_{\mathbf{k}}(\boldsymbol{\theta}) = e^{i\mathbf{k}\cdot\boldsymbol{\theta}}$. The set of admissible momenta $\mathcal{S}_{\mathbf{k}} = \{\mathbf{k}_0, \pm\mathbf{k}_1, \pm\mathbf{k}_2, ...\}$ are determined by the boundary conditions. The diagonalized representation of the kernel is therefore

$$K(\boldsymbol{\theta}, \boldsymbol{\theta}') = \sum_{\mathbf{k}\in\mathcal{S}_{\mathbf{k}}} \lambda_{\mathbf{k}} e^{i\mathbf{k}\cdot(\boldsymbol{\theta}-\boldsymbol{\theta}')} \quad \text{(App.15)}$$

For example, if the space is the torus $\mathbb{T}^n = S^1 \times S^1 \times ... \times S^1$, then the space of admissable momenta are the points on the integer lattice $\mathcal{S}_{\mathbf{k}} = \mathbb{Z}^n = \{\mathbf{k} \in \mathbb{R}^n | k_i \in \mathbb{Z}\ \forall i = 1, ..., n\}$. Reality and symmetry of the kernel demand that $\text{Im}(\lambda_{\mathbf{k}}) = 0$ and $\lambda_{-\mathbf{k}} = \lambda_{\mathbf{k}} \geq 0$. Most of the models in this paper consider $\theta \sim \text{Unif}\left(S^1\right)$, where the kernel has the following Fourier/Mercer decomposition

$$\begin{aligned} K(\theta - \theta') &= \sum_{k=-\infty}^{\infty} \lambda_k e^{ik(\theta-\theta')} = 2\sum_{k=0}^{\infty} \lambda_k \cos\big(k(\theta - \theta')\big) \\ &= \sum_{k=0}^{\infty} \lambda_k \left[\sqrt{2}\cos(k\theta)\sqrt{2}\cos\left(k\theta'\right) + \sqrt{2}\sin(k\theta)\sqrt{2}\sin\left(k\theta'\right)\right] \end{aligned} \quad \text{(App.16)}$$

where we invoked the simple trigonometric identity $\cos(a-b) = \cos(a)\cos(b) + \sin(a)\sin(b)$. By recognizing that $\{\sqrt{2}\cos(k\theta), \sqrt{2}\sin(k\theta)\}_{k=0}^{\infty}$ form a complete orthonormal set of functions with respect to $\text{Unif}\left(S^1\right)$, we have identified this as the collection of kernel eigenfunctions.

#### 1.4 Invariant Kernels Possess Invariant Eigenfunctions

Suppose the kernel $K(\boldsymbol{\theta}, \boldsymbol{\theta}')$ is invariant to some set of transformations $t \in \mathcal{T}$, by which we mean that

$$K(t\boldsymbol{\theta}, \boldsymbol{\theta}') = K(\boldsymbol{\theta}, t\boldsymbol{\theta}') = K(\boldsymbol{\theta}, \boldsymbol{\theta}')\ ,\ \forall t \in \mathcal{T} \quad \text{(App.17)}$$

We will now show that any eigenfunction of such a kernel with nonzero eigenvalue must be an invariant function. Let $\psi_k(\boldsymbol{\theta})$ be an eigenfunction with eigenvalue $\lambda_k > 0$, then

$$\psi_k(t\boldsymbol{\theta}) = \frac{1}{\lambda_k}\int p(\boldsymbol{\theta}')K(\boldsymbol{\theta}', t\boldsymbol{\theta})d\boldsymbol{\theta}' = \frac{1}{\lambda_k}\int p(\boldsymbol{\theta}')K(\boldsymbol{\theta}', \boldsymbol{\theta})d\boldsymbol{\theta}' = \psi_k(\boldsymbol{\theta}) \tag{App.18}$$

This establishes that all functions which depend on $\mathcal{T}$ transformations must necessarily lie in the null-space of $K$.

### 2 Alternative Neural Codes with Same Kernel

#### 2.1 Orthogonal Transformations are Sufficient for Linear Kernel-Preserving Transformations

We will now show that for any linear transformation $\tilde{\mathbf{r}} = \mathbf{A}\mathbf{r}$ which preserves the inner product kernel $K(\boldsymbol{\theta}, \boldsymbol{\theta}')$, there exists an orthogonal matrix $\mathbf{Q}$ such that $\tilde{\mathbf{r}} = \mathbf{Q}\mathbf{r}$.

*Proof.* Let $\tilde{\mathbf{r}}(\boldsymbol{\theta}) = \mathbf{A}\mathbf{r}(\boldsymbol{\theta})$ for all stimuli $\boldsymbol{\theta}$. To preserve the kernel, we must have

$$K(\boldsymbol{\theta}, \boldsymbol{\theta}') = \tilde{\mathbf{r}}(\boldsymbol{\theta}) \cdot \tilde{\mathbf{r}}(\boldsymbol{\theta}') = \mathbf{r}(\boldsymbol{\theta}) \cdot \mathbf{r}(\boldsymbol{\theta}') \implies \mathbf{r}(\boldsymbol{\theta})\mathbf{A}^\top\mathbf{A}\mathbf{r}(\boldsymbol{\theta}') = \mathbf{r}(\boldsymbol{\theta}) \cdot \mathbf{r}(\boldsymbol{\theta}'). \tag{App.19}$$

Taking projections against each of the orthonormal eigenfunctions $\psi_\ell(\theta)$ (see Appendix 1A), we define vectors $\mathbf{u}_k$ as $\sqrt{\lambda_k}\mathbf{u}_k = \left\langle \mathbf{r}(\boldsymbol{\theta})\psi_k(\boldsymbol{\theta})\right\rangle_{\boldsymbol{\theta}}$, allowing us to express the SVD of the population code $\mathbf{r}(\boldsymbol{\theta}) = \sum_k \sqrt{\lambda_k}\mathbf{u}_k\psi_k(\boldsymbol{\theta})$. These vectors $\{\mathbf{u}_k\}$ are orthonormal $\mathbf{u}_k \cdot \mathbf{u}_\ell = \delta_{k\ell}$ since, by the definition of the kernel eigenfunctions $\psi_k$,

$$\begin{aligned}
\sqrt{\lambda_k\lambda_\ell}\mathbf{u}_k \cdot \mathbf{u}_\ell &= \left\langle \mathbf{r}(\boldsymbol{\theta}) \cdot \mathbf{r}(\boldsymbol{\theta}')\psi_k(\boldsymbol{\theta})\psi_\ell(\boldsymbol{\theta}')\right\rangle_{\boldsymbol{\theta},\boldsymbol{\theta}'} \\
&= \left\langle \psi_k(\boldsymbol{\theta}) \left\langle K(\boldsymbol{\theta}, \boldsymbol{\theta}')\psi_\ell(\boldsymbol{\theta}')\right\rangle_{\boldsymbol{\theta}'}\right\rangle_{\boldsymbol{\theta}} \\
&= \lambda_\ell \left\langle \psi_k(\boldsymbol{\theta})\psi_\ell(\boldsymbol{\theta})\right\rangle_{\boldsymbol{\theta}} = \lambda_k\delta_{k,\ell}.
\end{aligned} \tag{App.20}$$

Since $\mathbf{r}(\boldsymbol{\theta})$ and $\tilde{\mathbf{r}}(\boldsymbol{\theta})$ have the same inner product kernel, they must posess the same kernel eigenfunctions $\psi_k$ and kernel eigenvalues $\lambda_k$, which are identified through the eigenvalue problem $\int p(\boldsymbol{\theta})K(\boldsymbol{\theta}, \boldsymbol{\theta}')\psi_k(\boldsymbol{\theta})d\boldsymbol{\theta} = \lambda_k\psi_k(\boldsymbol{\theta})$. We therefore have the following two singular value decompositions for $\mathbf{r}$ and $\tilde{\mathbf{r}}$

$$\mathbf{r}(\boldsymbol{\theta}) = \sum_{k=1}^{N} \sqrt{\lambda_k}\mathbf{u}_k\psi_k(\boldsymbol{\theta}) \ , \ \tilde{\mathbf{r}}(\boldsymbol{\theta}) = \sum_{k=1}^{N} \sqrt{\lambda_k}\tilde{\mathbf{u}}_k\psi_k(\boldsymbol{\theta}). \tag{App.21}$$

where $\{\mathbf{u}_k\}_{k=1}^N$ and $\{\tilde{\mathbf{u}}_k\}_{k=1}^N$ are both complete sets of orthonormal vectors (the sums above run over possible zero eigenvalues). Taking the equation $\tilde{\mathbf{r}}(\boldsymbol{\theta}) = \mathbf{A}\mathbf{r}(\boldsymbol{\theta})$, we multiply both sides of the equation by $\psi_k(\boldsymbol{\theta})$ and average over $\boldsymbol{\theta}$ giving

$$\left\langle \tilde{\mathbf{r}}(\boldsymbol{\theta})\psi_k(\boldsymbol{\theta}) \right\rangle = \sqrt{\lambda_k}\tilde{\mathbf{u}}_k = \mathbf{A}\left\langle \mathbf{r}(\boldsymbol{\theta})\psi_k(\boldsymbol{\theta}) \right\rangle_{\boldsymbol{\theta}} = \sqrt{\lambda_k}\mathbf{A}\mathbf{u}_k \tag{App.22}$$

For an eigenmode $k$ with positive eigenvalue $\lambda_k > 0$, this implies $\tilde{\mathbf{u}}_k = \mathbf{A}\mathbf{u}_k$, while there is no corresponding constraint for the null modes with $\lambda_k = 0$. However, the action of $\mathbf{A}$ on the nullspace of the code has no influence on $\tilde{\mathbf{r}}$ so there is no loss in generality to restrict consideration to transformations $\mathbf{A}$ which satisfy $\tilde{\mathbf{u}}_k = \mathbf{A}\mathbf{u}_k$ for all $k \in [N]$ (rather than just the $\lambda_k > 0$ modes). This choice gives $\mathbf{A} = \sum_{k=1}^{N} \tilde{\mathbf{u}}_k \mathbf{u}_k^\top \in O(N)$. Thus, the space of codes $\tilde{\mathbf{r}}(\boldsymbol{\theta})$ with equivalent kernels to $\mathbf{r}(\boldsymbol{\theta}) \cdot \mathbf{r}(\boldsymbol{\theta}')$ generated through linear transformations is equivalent to all possible orthogonal transformations of the original code $\{\mathbf{Q}\mathbf{r}(\boldsymbol{\theta}) : \mathbf{Q} \in O(N)\}$. □

#### 2.2 Necessary Conditions for Optimally Sparse Codes

Next we argue why optimally sparse codes should be lifetime and population selective. We consider the following optimization problem: find a non-negative neural responses $\mathbf{S} \in \mathbb{R}^{N\times P}$ and baseline vector $\boldsymbol{\delta} \in \mathbb{R}^N$ so that baseline subtracted responses $\mathbf{R} = \mathbf{S} - \boldsymbol{\delta}\mathbf{1}^\top$ realize a desired inner product kernel $\mathbf{K} \in \mathbb{R}^{P\times P}$ and have minimal total firing. This is equivalent to finding the most metabolically efficient code among the space of codes with equivalent inductive bias. Mathematically, we formulate this problem as

$$\min_{\mathbf{S}\in\mathbb{R}^{N\times P},\, \boldsymbol{\delta}\in\mathbb{R}^N} \sum_{i\mu} S_{i\mu}, \quad \text{s.t.} \ \left(\mathbf{S} - \boldsymbol{\delta}\mathbf{1}^\top\right)^\top \left(\mathbf{S} - \boldsymbol{\delta}\mathbf{1}^\top\right) = \mathbf{K}, \quad S_{i\mu} \geq 0 \ \ \forall i \in [N], \mu \in [P]. \tag{App.23}$$

To enforce the constraints for the definition of the kernel and the non-negativity of the responses, we introduce the following Lagrangian

$$\mathcal{L}(\mathbf{S}, \boldsymbol{\delta}, \mathbf{A}, \mathbf{V}) = \mathbf{1}^\top\mathbf{S}\mathbf{1} - \operatorname{Tr}\left(\left[(\mathbf{S} - \boldsymbol{\delta}\mathbf{1}^\top)^\top(\mathbf{S} - \boldsymbol{\delta}\mathbf{1}^\top) - \mathbf{K}\right]\mathbf{A}\right) - \operatorname{Tr}\mathbf{V}^\top\mathbf{S} \tag{App.24}$$

where $\mathbf{1}$ is the vector containing all ones, the Lagrange multiplier matrix $\mathbf{A}$ enforces the definition of the kernel and the KKT multiplier matrix $\mathbf{V}$ enforces the non-negativity constraints for each element of $\mathbf{S}$. The KKT conditions require that any local optimum of the objective would have to satisfy the following equations [3]

$$\begin{aligned}
\frac{\partial \mathcal{L}}{\partial \mathbf{S}} &= \mathbf{1}\mathbf{1}^\top - (\mathbf{S} - \boldsymbol{\delta}\mathbf{1}^\top)\mathbf{A} - \mathbf{V} = \mathbf{0} \\
\frac{\partial \mathcal{L}}{\partial \boldsymbol{\delta}} &= -(\mathbf{S} - \boldsymbol{\delta}\mathbf{1}^\top)\mathbf{A}\mathbf{1} = \mathbf{0} \\
\frac{\partial \mathcal{L}}{\partial \mathbf{A}} &= (\mathbf{S} - \boldsymbol{\delta}\mathbf{1}^\top)^\top(\mathbf{S} - \boldsymbol{\delta}\mathbf{1}^\top) - \mathbf{K} = \mathbf{0} \\
\mathbf{V} \odot \mathbf{S} &= \mathbf{0},
\end{aligned} \tag{App.25}$$

where $\odot$ denotes the element-wise Hadamard product. Using the complementary slackness condition $\mathbf{S} \odot \mathbf{V} = \mathbf{0}$, and the first optimality condition $\frac{\partial \mathcal{L}}{\partial \mathbf{S}} = \mathbf{0}$, we have

$$\mathbf{S} = \mathbf{S} \odot (\mathbf{S} - \boldsymbol{\delta}\mathbf{1}^\top)\mathbf{A} \tag{App.26}$$

Therefore, for any neuron-stimulus pair $(i, \mu)$, either $S_{i\mu} = 0$ or $\sum_{\nu \in [P]}(S_{i\nu} - \delta_i)A_{\nu\mu} = 1$. Further, under the condition that $\mathbf{K}$ is full rank, we conclude that for any stimulus $\mu$, $\sum_{\nu \in [P]} A_{\mu\nu} = 0$ from the equation $\frac{\partial \mathcal{L}}{\partial \boldsymbol{\delta}} = \mathbf{0}$. Let $\mathcal{I}_i = \{\mu \in [P] : S_{i\mu} > 0\}$ represent the set of stimuli for which neuron $i$ fires. We will call this the *receptive field set* for neuron $i$. Let $\mathbf{B}_{(i)} \in \mathbb{R}^{P \times P}$ have entries

$$[\mathbf{B}_{(i)}]_{\mu\nu} = \begin{cases} [\mathbf{A}^+_{(i)}]_{\mu\nu} & \mu, \nu \in \mathcal{I}_i \\ 0 & \mu \notin \mathcal{I}_i \text{ or } \nu \notin \mathcal{I}_i \end{cases} \tag{App.27}$$

where the matrix $\mathbf{A}_{(i)}$ is the $|\mathcal{I}_i| \times |\mathcal{I}_i|$ minor of $\mathbf{A}$ obtained by taking all rows and columns with indices $\mu, \nu \in \mathcal{I}_i$, and $\mathbf{A}^+$ denotes pseudo-inverse of $\mathbf{A}$. Then the $i$-th neuron's tuning curve is a function of the index set $\mathcal{I}_i$ the baseline $\delta_i$ and the neuron-independent $P \times P$ matrix $\mathbf{A}$: $\mathbf{s}(\mathcal{I}_i, \delta_i, \mathbf{A}) = \mathbf{B}_{(i)} [\delta_i \mathbf{A} + \mathbf{I}] \mathbf{1} \in \mathbb{R}^P$. The non-negativity constraint for neuron $i$'s tuning curve implies that $S_{i\mu} = \sum_{\nu \in \mathcal{I}_i} B_{(i),\mu\nu}[\delta_i \sum_{\gamma \in [P]} A_{\nu\gamma} + 1] > 0$ for all $\mu \in \mathcal{I}_i$. To satisfy the definition of the kernel, we have the following constraint on the matrix $\mathbf{A}$, the index sets $\mathcal{I}_i$ and baselines $\delta_i$

$$\mathbf{K} = \sum_{i=1}^{N} (\mathbf{s}(\mathcal{I}_i, \delta_i, \mathbf{A}) - \delta_i \mathbf{1})(\mathbf{s}(\mathcal{I}_i, \delta_i, \mathbf{A}) - \delta_i \mathbf{1})^\top \tag{App.28}$$

This equation implictly defines the index sets $\mathcal{I}_i$ the baselines $\delta_i$ and the KKT matrix $\mathbf{A}$. We see that, in order to fit an arbitrary kernel, the receptive field sets $\{\mathcal{I}_i\}$ and baselines $\delta_i$ for each neuron must be sufficiently diverse since otherwise only a low rank kernel matrix can be achieved from the optimally sparse code. As a concrete example, suppose that $\mathcal{I}_i = \mathcal{I}$ so that $\mathbf{B}_{(i)} = \mathbf{B}$ and $\delta_i = \delta$ for all $i$. For example, this could occur if each neuron fired for every possible stimulus. In this case, the kernel would be rank one: $\mathbf{K} = N(\mathbf{s}(\mathcal{I}, \delta, \mathbf{A}) - \delta\mathbf{1})(\mathbf{s}(\mathcal{I}, \delta, \mathbf{A}) - \delta\mathbf{1})^\top$. In order to achieve a higher rank code there must be sufficient diversity of the receptive fields $\mathcal{I}_i$. Thus the only way for optimally sparse codes to realize high rank kernels $\mathbf{K}$ is to have neurons to have different receptive field sets $\mathcal{I}_i$. The necessary optimality conditions thus reveal a preference for sparse neural tuning curves to have high *lifetime sparseness*; to achieve diverse index sets $\mathcal{I}_i$, any given neuron will fire only for a unique subset of the possible stimuli.

### 3 Theory of Generalization

#### 3.1 Convergence of the delta-rule

Gradient descent training of readout weights $\mathbf{w}$ on a finite sample of size $P$ converges to the kernel regression solution [4, 5, 6]. Let $\mathcal{D} = \{\boldsymbol{\theta}^\mu, y^\mu\}_{\mu=1}^P$ be the dataset with samples $\boldsymbol{\theta}^\mu$ and target values $y^\mu$. We introduce a shorthand $\mathbf{r}^\mu = \mathbf{r}(\boldsymbol{\theta}^\mu)$ for convenience. The empirical loss we aim to minimize is a sum of the squared losses of each data point in the training set

$$\mathcal{L}(\mathbf{w}) = \frac{1}{2}\sum_{\mu=1}^{P}(\mathbf{r}^\mu \cdot \mathbf{w} - y^\mu)^2. \tag{App.29}$$

Performing gradient descent updates

$$\mathbf{w}_{t+1} = \mathbf{w}_t - \eta\frac{\partial \mathcal{L}}{\partial \mathbf{w}_t} = \mathbf{w}_t - \eta\sum_{\mu=1}^{P}\mathbf{r}^\mu(\mathbf{r}^\mu \cdot \mathbf{w}_t - y^\mu), \tag{App.30}$$

recovers the delta rule that we discussed in the main text [7, 8]. Letting the empirical response matrix $\mathbf{R} = [\mathbf{r}^1, ..., \mathbf{r}^P] \in \mathbb{R}^{N\times P}$ have a SVD $\mathbf{R} = \sum_k \sqrt{\hat{\lambda}_k}\hat{\mathbf{u}}_k\hat{\boldsymbol{\psi}}_k^\top$, and expanding the weights $\mathbf{w}_t = \sum_k w_{t,k}\hat{\mathbf{u}}_k$ and labels $\mathbf{y} = \sum_k \hat{v}_k\hat{\boldsymbol{\psi}}_k$ in their respective SVD bases, we find

$$w_{t+1,k} = w_{t,k} - \eta\hat{\lambda}_k w_{t,k} + \eta\sqrt{\hat{\lambda}_k}\hat{v}_k \tag{App.31}$$

For all directions with $\hat{\lambda}_k > 0$, the dynamics converge to the unique fixed point $w_k^* = \frac{\hat{v}_k}{\sqrt{\hat{\lambda}_k}}$, while for all modes with $\hat{\lambda}_k = 0$, the weights remain at $w_k^* = 0$. Thus

$$\mathbf{w}^* = \left[\sum_{k:\hat{\lambda}_k>0}\frac{\hat{\mathbf{u}}_k\hat{\boldsymbol{\psi}}_k^\top}{\sqrt{\hat{\lambda}_k}}\right]\mathbf{y} = \mathbf{R}\left[\sum_{k:\hat{\lambda}_k>0}\frac{\hat{\boldsymbol{\psi}}_k\hat{\boldsymbol{\psi}}_k^\top}{\hat{\lambda}_k}\right]\mathbf{y} = \mathbf{R}\mathbf{K}^+\mathbf{y} \tag{App.32}$$

where $\mathbf{K}^+$ is the Moore-Penrose inverse of the kernel matrix $K_{\mu,\nu} = K(\boldsymbol{\theta}^\mu, \boldsymbol{\theta}^\nu)$. The predictions of the learned function are given by $f = \mathbf{w}^* \cdot \mathbf{r}(\boldsymbol{\theta})$ which can be expressed as

$$f(\boldsymbol{\theta}) = \mathbf{k}(\boldsymbol{\theta})^\top\mathbf{K}^+\mathbf{y} \tag{App.33}$$

The fact that the solution can be written in terms of a linear combination of $\{K(\boldsymbol{\theta}, \boldsymbol{\theta}^\mu)\}_{\mu=1}^P$ is known as the representer theorem [9, 2]. A similar analysis for nonlinear readouts where $f(\boldsymbol{\theta}) = g\left(\mathbf{w}\cdot\mathbf{r}(\boldsymbol{\theta})\right)$ is provided in Appendix 3G.

##### 3.1.1 Weight Decay and Ridge Regression

We can introduce a regularization term in our learning problem which penalizes the size of the readout weights. This leads to a modified learning objective of the form

$$\mathcal{L}(\mathbf{w}) = \sum_{\mu} (\mathbf{r}^{\mu} \cdot \mathbf{w} - y^{\mu})^2 + \lambda ||\mathbf{w}||^2. \tag{App.34}$$

Inclusion of this regularization alters the learning rule through *weight decay* $\mathbf{w}_{t+1} = (1-\eta\lambda)\mathbf{w}_t + \eta \sum_{\mu} \mathbf{r}^{\mu}(\mathbf{r}^{\mu} \cdot \mathbf{w}_t - y^{\mu})$, which multiplies the existing weight value by a factor of $1-\eta\lambda$ before adding the data dependent update. This learning problem and gradient descent dynamics have a closed form solution

$$f(\boldsymbol{\theta}) = \mathbf{r}(\boldsymbol{\theta}) \cdot \mathbf{w}^* = \sum_{\mu=1}^{P} \alpha^{\mu} K(\boldsymbol{\theta}, \boldsymbol{\theta}^{\mu}) \ , \ \boldsymbol{\alpha} = (\mathbf{K} + \lambda \mathbf{I})^{-1} \mathbf{y}. \tag{App.35}$$

The generalization benefits of explicit regularization through weight decay is known to be related to the noise statistics in the learning problem [10]. We simulate weight decay only in Figure 6C, where we use $\lambda = 0.01 \sum_k \lambda_k$ to improve numerical stability at large $P$.

#### 3.2 Computation of Learning Curves

Recent work has established analytic results that predict the average case generalization error for kernel regression

$$E_g = \left\langle E_g(\mathcal{D}) \right\rangle_{\mathcal{D}} = \left\langle (f(\boldsymbol{\theta}, \mathcal{D}) - y(\boldsymbol{\theta}))^2 \right\rangle_{\boldsymbol{\theta}, \mathcal{D}} \tag{App.36}$$

where $E_g(\mathcal{D}) = \left\langle (f(\boldsymbol{\theta}, \mathcal{D}) - y(\boldsymbol{\theta}))^2 \right\rangle_{\boldsymbol{\theta}}$ is the generalization error for a certain sample $\mathcal{D}$ of size $P$ and $f(\boldsymbol{\theta}, \mathcal{D})$ is the kernel regression solution for $\mathcal{D}$ [11, 10]. The typical or average case error $E_g$ is obtained by averaging over all possible datasets of size $P$. This average case generalization error is determined solely by the decomposition of the target function $y(\mathbf{x})$ along the eigenbasis of the kernel and the eigenspectrum of the kernel. This diagonalization takes the form

$$\int p(\boldsymbol{\theta}) K(\boldsymbol{\theta}, \boldsymbol{\theta}') \psi_k(\boldsymbol{\theta}) d\boldsymbol{\theta} = \lambda_k \psi_k(\boldsymbol{\theta}') \tag{App.37}$$

Since the eigenfunctions form a complete set of square integrable functions, we expand both the target function $y(\boldsymbol{\theta})$ and the learned function $f(\boldsymbol{\theta})$ in this basis

$$y(\boldsymbol{\theta}) = \sum_k v_k \psi_k(\boldsymbol{\theta}) \ , \ f(\boldsymbol{\theta}) = \sum_k w_k \psi_k(\boldsymbol{\theta}) \tag{App.38}$$

Due to the orthonormality of the kernel eigenfunctions $\{\psi_k\}$, the generalization error for any set of coefficients $\mathbf{w}$ is

$$E_g(\mathbf{w}) = \left\langle (y(\boldsymbol{\theta}) - f(\boldsymbol{\theta}))^2 \right\rangle_{\boldsymbol{\theta}} = \sum_k (w_k - v_k)^2 = ||\mathbf{w} - \mathbf{v}||^2 \tag{App.39}$$

We now introduce training error, or empirical loss, which depends on the disorder in the dataset $\mathcal{D} = \{(\boldsymbol{\theta}^\mu, y^\mu)\}_{\mu=1}^P$

$$H(\hat{\mathbf{v}}, \mathcal{D}) = \sum_\mu (\mathbf{w} \cdot \psi(\boldsymbol{\theta}^\mu) - \mathbf{v} \cdot \psi(\boldsymbol{\theta}^\mu))^2 + \lambda \sum_k \frac{w_k^2}{\lambda_k} \tag{App.40}$$

It is straightforward to verify that the optimal $\hat{\mathbf{v}}^*$ which minimizes $H(\hat{\mathbf{v}}, \mathcal{D})$ is the kernel regression solution for kernel with eigenvalues $\{\lambda_k\}$ when $\lambda \to 0$. The optimal weights $\hat{\mathbf{v}}$ can be identified through the first order condition $\nabla H(\hat{\mathbf{v}}, \mathcal{D}) = 0$ which gives

$$\mathbf{w}^* = (\boldsymbol{\Psi}\boldsymbol{\Psi}^\top + \lambda \boldsymbol{\Lambda}^{-1})^{-1} \boldsymbol{\Psi}\boldsymbol{\Psi}^\top \mathbf{v} = \mathbf{v} - \lambda (\boldsymbol{\Psi}\boldsymbol{\Psi}^\top + \lambda \boldsymbol{\Lambda}^{-1})^{-1} \boldsymbol{\Lambda}^{-1} \mathbf{v} \tag{App.41}$$

where $\Psi_{k,\mu} = \psi_k(\boldsymbol{\theta}^\mu)$ are the eigenfunctions evaluated on the training data and $\Lambda_{k,\ell} = \delta_{k,\ell}\lambda_k$ is a a diagonal matrix containing the kernel eigenvalues. The generalization error for this optimal solution is

$$E_g(\mathcal{D}) = ||\mathbf{w}^* - \mathbf{v}||^2 = \mathbf{v}^\top \boldsymbol{\Lambda}^{-1} \mathbf{G}(\mathcal{D})^2 \boldsymbol{\Lambda}^{-1} \mathbf{v} \ , \ \mathbf{G}(\mathcal{D}) = \left( \frac{1}{\lambda} \boldsymbol{\Psi}\boldsymbol{\Psi}^\top + \boldsymbol{\Lambda}^{-1} \right)^{-1} \tag{App.42}$$

We note that the dependence on the randomly sampled dataset $\mathcal{D}$ only appears through the matrix $\mathbf{G}(\mathcal{D})$. Thus to compute the *typical* generalization error we need to average over this matrix $\left\langle \mathbf{G}(\mathcal{D}) \right\rangle_{\mathcal{D}}$. There are multiple strategies to perform such an average and we will study one here based on a partial differential equation which was introduced in [12, 13] and studied further in [11, 10]. In this setting, we denote the average matrix $\mathbf{G}(P) = \left\langle \mathbf{G}(\mathcal{D}) \right\rangle_{|\mathcal{D}|=P}$ for a dataset of size $P$. We first will derive a recursion relationship using the Sherman Morrison formula for a rank-1 update to an inverse matrix. We imagine adding a new sampled feature vector $\phi$ to a dataset $\psi$ with size $P$. The average matrix $\mathbf{G}(P+1)$ at $P+1$ samples can be related to $\mathbf{G}(P)$ through the Sherman Morrison rule

$$\begin{aligned} \mathbf{G}(P+1) = \left\langle \left( \frac{1}{\lambda} \boldsymbol{\Psi}\boldsymbol{\Psi}^\top + \frac{1}{\lambda} \psi\psi^\top + \boldsymbol{\Lambda}^{-1} \right)^{-1} \right\rangle_{\psi, \mathcal{D}} &= \mathbf{G}(P) - \left\langle \frac{\mathbf{G}(\mathcal{D})\psi\psi^\top \mathbf{G}(\mathcal{D})}{\lambda + \psi^\top \mathbf{G}(\mathcal{D})\psi} \right\rangle_{\psi, \mathcal{D}} \\ &\approx \mathbf{G}(P) - \frac{\left\langle \mathbf{G}(\mathcal{D}) \left\langle \psi\psi^\top \right\rangle_\psi \mathbf{G}(\mathcal{D}) \right\rangle_{\mathcal{D}}}{\lambda + \left\langle \psi^\top \mathbf{G}(\mathcal{D})\psi \right\rangle_{\psi, \mathcal{D}}} \end{aligned} \tag{App.43}$$

where in the last step we approximated the average of the ratio with the ratio of averages. This operation, is of course, unjustified theoretically, but has been shown to produce accurate learning curves [11, 13]. Since the chosen basis of kernel eigenfunctions are orthonormal, the average over the new sample is trivial $\left\langle \psi\psi^\top \right\rangle_\phi = \mathbf{I}$. We thus arrive at the following recursion relationship for $\mathbf{G}$

$$\mathbf{G}(P+1) = \mathbf{G}(P) - \frac{\left\langle \mathbf{G}(\mathcal{D})^2 \right\rangle_{\mathcal{D}}}{\lambda + \mathrm{Tr}\ \mathbf{G}(P)} \tag{App.44}$$

By introducing an additional source $J$ so that $\mathbf{G}(\mathcal{D}, J)^{-1} = \frac{1}{\lambda} \boldsymbol{\Psi}\boldsymbol{\Psi}^\top + \boldsymbol{\Lambda}^{-1} + J\mathbf{I}$, we can relate

$\mathbf{G}(\mathcal{D}, J)$'s first and second moments through differentiation

$$\frac{\partial}{\partial J}\mathbf{G}(P,J) = \frac{\partial}{\partial J}\left\langle \left(\frac{1}{\lambda}\mathbf{\Psi}\mathbf{\Psi}^\top + J\mathbf{I} + \mathbf{\Lambda}^{-1}\right)^{-1}\right\rangle_{\mathcal{D}} = -\left\langle \mathbf{G}(\mathcal{D},J)^2\right\rangle_{\mathcal{D}}. \tag{App.45}$$

Thus the recursion relation simplifies to

$$\mathbf{G}(P+1,J) - \mathbf{G}(P,J) \approx \frac{\partial}{\partial p}\mathbf{G}(p,J) = \frac{1}{\lambda + \mathrm{Tr}\mathbf{G}(P,J)}\frac{\partial}{\partial J}\mathbf{G}(P,J), \tag{App.46}$$

where we approximated the finite difference in $P$ as a derivative, treating $P$ as a continuous variable. Taking the trace of both sides and defining $\kappa(P,J) = \lambda + \mathrm{Tr}\mathbf{G}(P,J)$ we arrive at the following quasilinear PDE

$$\frac{\partial}{\partial P}\kappa(P,J) = \frac{1}{\kappa(P,J)}\frac{\partial}{\partial J}\kappa(P,J) \tag{App.47}$$

with the initial condition $\kappa(0,J) = \lambda + \mathrm{Tr}(\mathbf{\Lambda}^{-1} + J\mathbf{I})^{-1}$. Using the method of characteristics, we arrive at the solution $\kappa(P,J) = \lambda + \mathrm{Tr}\left(\mathbf{\Lambda}^{-1} + (v + \frac{P}{\kappa(P,J)})\mathbf{I}\right)^{-1}$. Using this solution to $\kappa$, we can identify the solution to $\mathbf{G}$

$$\mathbf{G}(P,J)_{k,\ell} = \left(\frac{P}{\kappa} + J + \lambda_k^{-1}\right)^{-1}\delta_{k,\ell} = \frac{\kappa\lambda_k}{\lambda_k P + \kappa + J\kappa\lambda_k}\delta_{k,\ell}. \tag{App.48}$$

The generalization error, therefore can be written as

$$E_g = \mathbf{v}^\top\mathbf{\Lambda}^{-1}\left\langle \mathbf{G}(\mathcal{D})^2\right\rangle_{\mathcal{D}}\mathbf{\Lambda}^{-1}\mathbf{v} = -\frac{\partial}{\partial J}\mathbf{v}^\top\mathbf{\Lambda}^{-1}\mathbf{G}(P,J)\mathbf{\Lambda}^{-1}\mathbf{v} \tag{App.49}$$

$$= -\sum_k \frac{v_k^2}{\lambda_k^2}\frac{\partial}{\partial J}\left(\frac{P}{\kappa} + J + \lambda_k^{-1}\right)^{-1} = \frac{\kappa^2}{1-\gamma}\sum_k \frac{v_k^2}{(\lambda_k P + \kappa)^2}, \tag{App.50}$$

where $\gamma = P\sum_k \frac{\lambda_k^2}{(\lambda_k P+\kappa)^2}$, giving the desired result. Note that $\kappa$ depends on $J$ implicitly, which is the source of the $\frac{1}{1-\gamma}$ factor. This result was recently reproduced using techniques from statistical mechanics [11, 10].

##### 3.3 Spectral Bias and Code-Task Alignment

Through implicit differentiation it is straightforward to verify that the ordering of the mode errors $E_k = \frac{\kappa^2}{1-\gamma}(\lambda_k P + \kappa)^{-2}$ matches the ordering of the eigenvalues [10]. Let $\lambda_k > \lambda_\ell$, then we have

$$\begin{aligned}\frac{d}{dP}\log\left(\frac{E_k}{E_\ell}\right) =& 2\left[\frac{\lambda_\ell}{\lambda_\ell P + \kappa} - \frac{\lambda_k}{\lambda_k P + \kappa}\right] \\ &+ 2\kappa'(P)\left[\frac{1}{\lambda_\ell P + \kappa} - \frac{1}{\lambda_k P + \kappa}\right].\end{aligned} \tag{App.51}$$

Since $\lambda_\ell < \lambda_k$, the first bracket must be negative and the second bracket must be positive. Further, it is straightforward to compute that $\kappa'(P) = -\frac{\kappa\gamma}{P(1+\gamma)} < 0$. Therefore $\lambda_k > \lambda_\ell$ implies

$\frac{d}{dP} \log\left(\frac{E_k}{E_\ell}\right) < 0$ for all $P$. Since $\log\left(\frac{E_k}{E_\ell}\right) = 0$ at $P = 0$ we therefore have that $\log\big(E_k/E_\ell\big) < 0$ for all $P$ and consequently $E_k < E_\ell$. Modes with larger eigenvalues $\lambda_k$ have lower normalized mode errors $E_k$. This observation can be used to prove that target functions acting on the same data distribution with higher cumulative power distributions $C(k)$ for all $k$ will have lower generalization error normalized by total target power, $E_g(P)/E_g(0)$, for all $P$. Proof can be found in [10].

#### 3.4 Asymptotic power law scaling of learning curves

*Exponential Spectral Decays*: First, we will study the setting relevant to the von-Mises kernel where $\lambda_k \sim \beta^k$ and $v_k^2 \sim \alpha^k$ where $\alpha, \beta < 1$. This exponential behavior accounts for differences in bandwidth between kernels which modulates the base $\beta$ of the exponential scaling of $\lambda_k$ with $k$. We will approximate the sum over all mode errors with an integral

$$E_g = \frac{\kappa^2}{1-\gamma} \sum_{k=0}^{\infty} \frac{v_k^2}{(\lambda_k P + \kappa)^2} \sim \kappa^2 \int_0^\infty \frac{\alpha^k}{(\beta^k P + \kappa)^2} dk. \tag{App.52}$$

If we include a regularization parameter $\lambda$, then $\kappa \sim \lambda$ as $P \to \infty$. With this fact, we can therefore approximate the integral at large $P$ by splitting it up into all $k < k^* = \ln\big(P/\lambda\big)/\ln\big(1/\beta\big)$ and $k > k^*$.

$$E_g \sim \frac{\lambda^2}{P^2} \int_0^{k^*} \frac{\alpha^k}{\beta^{2k}\left[1 + \frac{\lambda}{P\beta^k}\right]^2} dk + \int_{k^*}^\infty \frac{\alpha^k}{\left[1 + \frac{\beta^k P}{\lambda}\right]^2} dk = A P^{-\frac{\log(1/\alpha)}{\log(1/\beta)}} + \sum_{n=0}^\infty A_n P^{-n-2} \tag{App.53}$$

for $P$-independent constants $A$ and $A_n$. Thus, we obtain a power law scaling of the learning curve $E_g$ which is dominated at large $P$ by $E_g \sim P^{-\min\left(2, \frac{\ln(1/\alpha)}{\ln(1/\beta)}\right)}$. For the von-Mises kernel we can approximate the spectra with $\lambda_k \sim \sigma^{-2k}$ and $v_k^2 \sim \sigma_T^{-2k}$ giving rise to a generalization scaling scaling $E_g \sim P^{-\min\left(2, \frac{\ln \sigma_T}{\ln \sigma}\right)}$.

*Power Law Spectral Decays*: The same arguments can be applied for power law kernels $\lambda_k \sim k^{-b}$ and power law targets $v_k^2 \sim k^{-a}$, which is of interest due to its connection to nonlinear rectified neural populations. In this setting, the generalization error is

$$\begin{aligned} E_g &\approx \int_1^\infty \frac{k^{-a}}{(k^{-b}P + \kappa)^2} dk \approx \frac{\kappa^2}{P^2} \int_1^{P^{1/b}} k^{-a+2b} dk + \int_{P^{1/b}}^\infty k^{-a} dk \\ &= \frac{1}{P^2(1-a+2b)} \left[P^{(1-a)/b+2} - 1\right] + \frac{1}{a-1} P^{(1-a)/b}. \end{aligned} \tag{App.54}$$

We see that there are two possible power law scalings for $E_g$ with the exponents $(a-1)/b$ and 2. At large $P$ this formula will be dominated by the term with minimum exponent so $E_g \sim P^{-\min(a-1, 2b)/b}$.

##### 3.5 Laplace Kernel Generalization

We calculate similar learning curves as we did for the von-Mises kernel but with Laplace kernels to show that our results is not an artifact of the infinite differentiability of the Von Mises kernel. Each of these Laplace kernels has the same asymptotic power law spectrum $\lambda_k \sim o(k^{-2})$, exhibiting a discontinuous first derivative (Figure App.5 A). Despite having the same spectral scaling at large $k$, these kernels can give dramatically different performance in learning tasks, again indicating the influence of the top eigenvalues on generalization at small $P$ (Figure App.5). Again, the trend for which kernels perform best at low $P$ can be reversed at large $P$. In this case, all generalization errors scale with $E_g \sim P^{-2}$ (Figure App.5B). More generally, our theory shows that if the task power spectrum and kernel eigenspectrum are both falling as power laws with exponents $a$ and $b$ respectively, then the generalization error asymptotically falls with a power law, $E_g \sim P^{-\min(a-1,2b)/b}$ (Methods) [11]. This decay is fastest when $b \geq \frac{a-1}{2}$ for which $E_g \sim P^{-2}$. Therefore, the tail of the kernel's eigenvalue spectrum determines the large sample size behavior of the generalization error for power law kernels. Small sample size limit is still governed by the bulk of the spectrum.

##### 3.6 Learning with Multiple Output Channels

Our theory is not limited to scalar target functions but rather can be easily extended to multiple output functions $y_1, ..., y_C$ from the same data, if for example the task requires computing class membership for $C$ categories. In this setting, each data point has the form $(\boldsymbol{\theta}^\mu, \mathbf{y}^\mu)$ with $\mathbf{y}^\mu \in \mathbb{R}^C$. For these $C$ classes, the generalization error takes the form

$$E_g = \left\langle ||\mathbf{f}(\boldsymbol{\theta}) - \mathbf{y}(\boldsymbol{\theta})||^2 \right\rangle = \sum_{c=1}^{C} \left\langle \left(f_c(\boldsymbol{\theta}) - y_c(\boldsymbol{\theta})\right)^2 \right\rangle = \sum_k \left[ \sum_c \left\langle y_c(\boldsymbol{\theta}) \phi_k(\boldsymbol{\theta}) \right\rangle^2 \right] E_k \ . \tag{App.55}$$

We therefore find that the generalization error in the multi-class setting is the same as the $E_g$ obtained for a single scalar target function with power spectrum $v_k^2 = \sum_c \left\langle y_c(\boldsymbol{\theta}) \phi_k(\boldsymbol{\theta}) \right\rangle^2$ [11, 10]. The relevant cumulative power distribution measures the fraction of total output variance captured by the first $k$ eigenfunctions of the population code

$$C(k) = \frac{\sum_c \sum_{\ell=1}^{k} \left\langle y_c(\boldsymbol{\theta}) \phi_k(\boldsymbol{\theta}) \right\rangle^2}{\sum_c \sum_{\ell=1}^{\infty} \left\langle y_c(\boldsymbol{\theta}) \phi_k(\boldsymbol{\theta}) \right\rangle^2} \ . \tag{App.56}$$

##### 3.7 Convergence of Delta-Rule for Nonlinear Readouts

In this section, we consider gradient descent dynamics on a least squares cost with nonlinear readout function. Let $\mathcal{D} = \{\boldsymbol{\theta}^\mu, y^\mu\}_{\mu=1}^P$ be the dataset with samples $\boldsymbol{\theta}^\mu$ and target values $y^\mu$. We introduce a shorthand $\mathbf{r}^\mu = \mathbf{r}(\boldsymbol{\theta}^\mu)$ for convenience. We consider output neurons which produce activity $f(\boldsymbol{\theta}) = g(\mathbf{w} \cdot \mathbf{r}(\boldsymbol{\theta}))$ for invertible nonlinear function $g$ with non-vanishing gradient. The empirical loss we aim to minimize is a sum of the squared losses of each data point in the training set

$$\mathcal{L}(\mathbf{w}) = \sum_{\mu=1}^{P} (g(\mathbf{r}^{\mu} \cdot \mathbf{w}) - y^{\mu})^2. \tag{App.57}$$

Performing gradient descent updates

$$\mathbf{w}_{t+1} = \mathbf{w}_t - \eta \frac{\partial \mathcal{L}}{\partial \mathbf{w}} = \mathbf{w}_t - \eta \sum_{\mu=1}^{P} \mathbf{r}^{\mu} g'(\mathbf{w} \cdot \mathbf{r}^{\mu})(g(\mathbf{r}^{\mu} \cdot \mathbf{w}_t) - y^{\mu}), \tag{App.58}$$

recovers the delta rule that we discussed in the main text for $g(z) = z$ [7, 8]. We define the variables $\mathbf{z}_t = \mathbf{R}^{\top}\mathbf{w}_t$ which satisfy the dynamics

$$\mathbf{z}_{t+1} - \mathbf{z}_t = \eta \mathbf{K} \; \mathrm{diag}(g'(\mathbf{z}_t)) \; (\mathbf{y} - g(\mathbf{z}_t)) \tag{App.59}$$

The possible fixed points of this system are the $\mathbf{z}$ which satisfy $\mathbf{K} \; \mathrm{diag}(g'(\mathbf{z})) \; (\mathbf{y} - g(\mathbf{z})) = 0$. We can make further progress under the condition that $\mathbf{K}$ is full rank (generally true in overparameterized setting $N > P$), the activation function is monotonic $g'(z) > 0$ for all $z$ and is invertible (some examples include softplus $g(z) = \ln(1 + e^z)$, sigmoids such as $g(z) = \tanh(z), \mathrm{erf}(z)$, etc.). If $y^{\mu}$ are in the range of $g$, then we obtain the simple fixed point condition $\mathbf{z}^* = g^{-1}(\mathbf{y})$. From this condition, we can infer the learned function. First note that $\mathbf{w}_t \in \mathrm{span}\{\mathbf{r}^{\mu}\}_{\mu=1}^{P}$ for all $t$ so that $\mathbf{w}^* = \mathbf{R}\boldsymbol{\alpha}^*$. Multiplying both sides of this last expression by $\mathbf{R}^{\top}$, we find

$$\mathbf{R}^{\top}\mathbf{R}\boldsymbol{\alpha}^* = \mathbf{R}^{\top}\mathbf{w}^* = \mathbf{K}\boldsymbol{\alpha}^* = \mathbf{z}^* \implies \boldsymbol{\alpha}^* = \mathbf{K}^{-1}\mathbf{z}^* = \mathbf{K}^{-1}g^{-1}(\mathbf{y}) \tag{App.60}$$

which is legitimate under the assumption that $\mathbf{K}$ was full rank. Now, the final learned function is merely $f(\boldsymbol{\theta}) = g(\mathbf{w}^* \cdot \mathbf{r}(\boldsymbol{\theta})) = g\left(\mathbf{r}(\boldsymbol{\theta})^{\top}\mathbf{R}\mathbf{K}^{-1}g^{-1}(\mathbf{y})\right) = g\left(\mathbf{k}(\boldsymbol{\theta})^{\top}\mathbf{K}^{-1}g^{-1}(\mathbf{y})\right)$.

##### 3.8 Comparison of our Theory with Recursive Least Squares Adaptive Filter

In the under-parameterized regime where $P > N$, the asymptotic error of recursive least squares adaptive filter has been obtained in prior works [14, 15]. Let $\mathbf{w}$ represent the weights and $\mathbf{r}_{\mu}$ represent the $\mu$-th presented example. We aim to reproduce a noisy ground truth $y_{\mu} = \mathbf{w}^* \cdot \mathbf{r}_{\mu} + \epsilon_{\mu}$. Analysis of the recursive least squares filter in the under-parameterized regime $P > N + 1$ can be performed exactly

$$\mathbf{w}(P) = \mathrm{argmin}_{\mathbf{w} \in \mathbb{R}^N} \sum_{\mu=1}^{P} [y_{\mu} - \mathbf{w} \cdot \mathbf{r}_{\mu}]^2 \tag{App.61}$$

This has a unique minimizer when $P > N$. Letting $\mathbf{C}(P) = \sum_{\mu=1}^{P} \mathbf{r}_{\mu}\mathbf{r}_{\mu}^{\top}$

$$
\begin{aligned}
\mathbf{w}(P) = \mathbf{C}(P)^{-1}\left[\sum_{\mu=1}^{P} y_\mu \mathbf{r}_\mu\right] &= \mathbf{C}(P)^{-1}\left[\mathbf{C}(P)\mathbf{w}^* + \sum_{\mu=1}^{P}\mathbf{r}_\mu \epsilon_\mu\right] \\
&= \mathbf{w}^* + \mathbf{C}(P)^{-1}\left[\sum_{\mu}\mathbf{r}_\mu \epsilon_\mu\right]
\end{aligned} \tag{App.62}
$$

This filter can be computed efficiently in an online manner using the Recursive Least Squares (RLS) algorithm. This recursion is obtained through the formula for the rank-one update to an inverse matrix

$$
\begin{aligned}
\mathbf{w}(P+1) &= \left[\mathbf{C}(P) + \mathbf{r}_{P+1}\mathbf{r}_{P+1}^\top\right]^{-1}\left[\sum_{\mu=1}^{P+1}\mathbf{r}_\mu \epsilon_\mu\right] \\
&= \mathbf{C}(P)^{-1}\left[\sum_{\mu=1}^{P+1}\mathbf{r}_\mu \epsilon_\mu\right] - \frac{\mathbf{C}(P)^{-1}\mathbf{r}_{P+1}\mathbf{r}_{P+1}^\top \mathbf{C}(P)^{-1}}{1+\mathbf{r}_{P+1}^\top \mathbf{C}(P)^{-1}\mathbf{r}_{P+1}}\left[\sum_{\mu=1}^{P+1}\mathbf{r}_\mu \epsilon_\mu\right] \\
&= \left[\mathbf{I} - \frac{\mathbf{C}(P)^{-1}\mathbf{r}_{P+1}\mathbf{r}_{P+1}^\top}{1+\mathbf{r}_{P+1}^\top \mathbf{C}(P)^{-1}\mathbf{r}_{P+1}}\right]\left[\mathbf{w}(P) + \mathbf{r}_{P+1}\epsilon_{P+1}\right].
\end{aligned} \tag{App.63}
$$

Under the assumption that the noise has covariance $\left\langle \epsilon\epsilon^\top \right\rangle = \sigma^2 \mathbf{I}$ and the neural responses are Gaussian $\mathbf{r}_\mu \sim \mathcal{N}(0, \mathbf{\Sigma})$, the excess generalization error can be obtained exactly using the formula for the average of an inverse Wishart matrix $\langle \mathbf{C}(P)^{-1}\rangle = \frac{1}{P-N-1}\mathbf{\Sigma}^{-1}$ giving

$$
\begin{aligned}
E_g - \sigma^2 &= \mathrm{Tr}\mathbf{\Sigma}\left\langle (\mathbf{w}(P)-\mathbf{w}^*)(\mathbf{w}(P)-\mathbf{w}^*)^\top \right\rangle_{\{\mathbf{r}_\mu, \epsilon_\mu\}} \\
&= \mathrm{Tr}\mathbf{\Sigma}\left\langle \mathbf{C}(P)^{-1}\mathbf{R}(P)\epsilon\epsilon^\top \mathbf{R}(P)^\top \mathbf{C}(P)^{-1}\right\rangle \\
&= \sigma^2 \mathrm{Tr}\mathbf{\Sigma}\left\langle \mathbf{C}(P)^{-1}\right\rangle = \sigma^2 \frac{\mathrm{Tr}[\mathbf{\Sigma}\mathbf{\Sigma}^{-1}]}{P-N-1} = \sigma^2 \frac{N}{P-N-1}.
\end{aligned} \tag{App.64}
$$

This analysis shows that RLS is consistent in the sense that as $P \to \infty$ with $N$ held constant the generalization error reach its theoretical minimum $\sigma^2$.

Though our main paper focuses on noise free estimation ($\sigma^2 = 0$) and the over-parameterized ($N > P$) rather than under parameterized case (which is what is discussed above), the above result agrees with what is obtained by our theory in the under-parameterized case since, in the $P > N$ limit, $\kappa = 0$ and $\gamma = \frac{N}{P}$. From our previous work, the error in the underparameterized noisy case for large $P$ is [10]

$$E_g - \sigma^2 = \sigma^2 \frac{\gamma}{1-\gamma} = \sigma^2 \frac{N/P}{1-N/P} = \sigma^2 \frac{N}{P-N} \tag{App.65}$$

which is agrees with the exact result up to an error which is negligible in the large $N, P$ limit. In this over-parameterized setting, the only contribution to the error comes from the explicit noise in the target function $\sigma^2$.

We note however, that the above technique does not generalize to the *overparameterized* setting where the system is trying to learn from a small number of examples $P < N$. In this case, when $\sigma = 0$, the delta-rule converges to the minimum norm solution which has the form

$$\mathbf{w}(P) = \mathbf{R}(P)\mathbf{K}(P)^{-1}\mathbf{y}(P) = \mathbf{R}(P)\mathbf{K}(P)^{-1}\mathbf{R}(P)^\top \mathbf{w}^* \tag{App.66}$$

where $\mathbf{K}(P) = \mathbf{R}(P)^\top \mathbf{R}(P)$. We see that the learned weight vector $\mathbf{w}(P)$ is a random rank-$P$ projection of the optimal weight vector $\mathbf{w}^*$. The challenging task is to compute an average over the random data $\mathbf{R}$ to get generalization error

$$E_g = \mathbf{w}^{*\top} \left\langle \left(\mathbf{I} - \mathbf{R}(P)\mathbf{K}(P)^{-1}\mathbf{R}(P)^\top\right) \boldsymbol{\Sigma} \left(\mathbf{I} - \mathbf{R}(P)\mathbf{K}(P)^{-1}\mathbf{R}(P)^\top\right) \right\rangle \mathbf{w}^* \tag{App.67}$$

We solve this problem by first making a relaxation to the $\lambda \to 0$ limit of ridge regression

$$\begin{aligned}
\mathbf{R}(P)\mathbf{K}^{-1}\mathbf{R}(P) &= \lim_{\lambda \to 0} \mathbf{R}(P) \left(\mathbf{K}(P) + \lambda \mathbf{I}\right)^{-1} \mathbf{R}(P)^\top = \lim_{\lambda \to 0} \left(\mathbf{C}(P) + \lambda \mathbf{I}\right)^{-1} \mathbf{C}(P) \\
&= \lim_{\lambda \to 0} \left[\mathbf{I} - \lambda \left(\mathbf{C}(P) + \lambda \mathbf{I}\right)^{-1}\right] \\
\implies E_g &= \lim_{\lambda \to 0} \mathbf{v}^\top \boldsymbol{\Lambda}^{-1} \left\langle \left(\frac{1}{\lambda} \boldsymbol{\Psi}\boldsymbol{\Psi}^\top + \boldsymbol{\Lambda}^{-1}\right)^{-2} \right\rangle \boldsymbol{\Lambda}^{-1} \mathbf{v} \\
&= - \lim_{\lambda \to 0} \lim_{z \to 0} \frac{\partial}{\partial z} \text{Tr} \left\langle \left(\frac{1}{\lambda} \boldsymbol{\Phi}\boldsymbol{\Phi}^\top + \boldsymbol{\Lambda}^{-1} + z \boldsymbol{\Lambda}^{-1} \mathbf{v}\mathbf{v}^\top \boldsymbol{\Lambda}^{-1}\right)^{-1} \right\rangle
\end{aligned} \tag{App.68}$$

where $\mathbf{v} = \boldsymbol{\Lambda}^{1/2}\mathbf{U}^\top \mathbf{w}^*$. This average can be characterized exactly for arbitrary $\lambda, \boldsymbol{\Lambda}, \mathbf{w}^*$ only in asymptotic limits where the number of samples $P$ and the number of features $N$ are both large. Though in this present work, we resort to the PDE approximation, our theory can also be derived with the replica method from disordered systems [11, 10].

#### 3.9 Comparison of our Theory with Wiener Filter and Equivalent Kernel

We will provide a discussion of how the theory of generalization used in our paper differs from these older results, such as those discussed in Chapter 7 of [2]. Many of the older asymptotic results for kernel regression with a ridge $\lambda$ can be obtained through an averaging argument (see for example Rasmussen and Williams Chapter 7.1 and 7.2 [2]). In the limit where the number of independent samples is large, we have by the strong law of large numbers, a convergence of the empirical loss to an expectation integral

$$\mathcal{L}[f] = \frac{1}{P}\sum_{\mu}\left(f(\boldsymbol{\theta}^{\mu}) - y(\boldsymbol{\theta}^{\mu})\right)^2 + \frac{\lambda}{P}||f||_{\mathcal{H}}^2 \sim \int p(\boldsymbol{\theta})\left(y(\boldsymbol{\theta}) - f(\boldsymbol{\theta})\right)^2 d\boldsymbol{\theta} + \frac{\lambda}{P}\sum_k \frac{\left\langle \psi_k(\boldsymbol{\theta}) f(\boldsymbol{\theta})\right\rangle_{\boldsymbol{\theta}}^2}{\lambda_k} \tag{App.69}$$

for a complete orthonormal family of eigenfunctions $\psi_k$. The minimizer to the above functional $f$ and its corresponding generalization error $E_g$ have the form

$$f(x) = \sum_k \frac{P\lambda_k}{P\lambda_k + \lambda} v_k \psi_k(x)\ ,\ E_g = \lambda^2 \sum_k \frac{v_k^2}{(P\lambda_k + \lambda)^2} \tag{App.70}$$

for target coefficients $v_k = \langle y(x)\psi_k(x)\rangle$. This equation (which is equivalent to the Wiener filter when $\psi_k$ are plane waves; see [2] Ch 7.1) is somewhat sensible in that this function approaches $y(x)$ as $P \to \infty$, and agrees with our theory in the limit of large ridge parameter $\lambda \to \infty$. Further, the large eigenvalue modes are learned at smaller values of $P$: it takes roughly $P \approx \lambda/\lambda_k$ samples to learn mode $k$. Note, however, that this theory is essentially useless in the interpolating limit $\lambda \to 0^+$ where it predicts that the learning curve has no dependence on sample size. This occurs because the fluctuations in the loss function from sample to sample (quenched disorder in statistical physics language) have been disregarded. Our analysis does not suffer from this issue and predicts the effects of sampling induced fluctuations in the loss function. Such sample fluctuations cannot typically be disregarded in thermodynamic limits where both dimension and dataset size are taken to infinity simultaneously [16]. A more complete discussion of the error due to sampling noise can be found in [10]. Our theory, in contrast to the Wiener filter described above, gives the following average predictor.

$$\langle f(x)\rangle_{\mathcal{D}} = \sum_k \frac{P\lambda_k}{P\lambda_k + \kappa} v_k \psi_k(x)\ ,\ \kappa = \lambda + \kappa \sum_k \frac{\lambda_k}{P\lambda_k + \kappa} \tag{App.71}$$

We see that even in the $\lambda \to 0$ limit, the mean predictor is not equivalent to $y(x)$ at finite $P$. Rather the sample complexity of each eigenmode $\psi_k$ is set by the evolution of the self-consistent function $\kappa$ which can be nonzero even in the $\lambda \to 0$ limit. The full generalization error can be decomposed into bias and variance components, where bias component corresponds to the error of the average predictor $\langle f(\mathbf{x})\rangle_{\mathcal{D}}$.

$$E_g = \left\langle \left( \langle f(\mathbf{x}) \rangle_{\mathcal{D}} - y(\mathbf{x}) \right)^2 \right\rangle_{\mathbf{x}} + \left\langle \left( f(\mathbf{x}) - \langle f(\mathbf{x}) \rangle_{\mathcal{D}} \right)^2 \right\rangle_{\mathbf{x},\mathcal{D}} \tag{App.72}$$

where the bias and variance have the forms

$$\text{Bias} = \left\langle \left( \langle f(\mathbf{x}) \rangle_{\mathcal{D}} - y(\mathbf{x}) \right)^2 \right\rangle_{\mathbf{x}} = \kappa^2 \sum_k \frac{v_k^2}{(P\lambda_k + \kappa)^2} \tag{App.73}$$

$$\text{Variance} = \left\langle \left( f(\mathbf{x}) - \langle f(\mathbf{x}) \rangle_{\mathcal{D}} \right)^2 \right\rangle_{\mathbf{x},\mathcal{D}} = \frac{\kappa^2 \gamma}{1-\gamma} \sum_k \frac{v_k^2}{(P\lambda_k + \kappa)^2} \tag{App.74}$$

$$\gamma = P \sum_k \frac{\lambda_k^2}{(P\lambda_k + \kappa)^2} \tag{App.75}$$

We see that the bias is identical to the generalization error predicted by the equivalent kernel provided that the self-consistent function $\kappa$ is substituted for the ridge parameter $\lambda$. This substitution allows for nonvanishing bias in the $\lambda \to 0$ limit since $\kappa > 0$ for $P$ smaller than the number of nonzero eigenvalues. We find that the sample-to-sample variance of the learned predictor is nonzero $\langle f(x)^2 \rangle_{\mathcal{D}} - \langle f(x) \rangle_{\mathcal{D}}^2 > 0$ and varies with $P$. This is unlike the Wiener filter theory which treats the variance as 0 for all $P$ due to concentration of the training loss.

We note a correspondence between the two theories in the heavily regularized $\lambda \to \infty$ limit. In this limit, $\kappa \sim \lambda$ and the bias approaches the risk predicted by the equivalent kernel theory. Further, the variance goes to zero in the $\lambda \to \infty$ limit since $\gamma \sim \lambda^{-2} \to 0$. A comparison of the two theories for linear ridge regression on random Gaussian design is provided in Figure App.7.

### 4 Visual Scene Reconstruction Task

#### 4.1 Reconstruction of Natural Scenes from Neural Responses

Using the mouse V1 responses to natural scenes, we attempt to reconstruct original images from the neural codes using different numbers of images. The presented natural scenes are taken from ten classes of imagenet which can be downloaded from `https://github.com/MouseLand/stringer-pachitariu-et-al-2018b`. Let $\boldsymbol{\theta}^\mu \in \mathbb{R}^D$ be a $D$-dimensional flattened vector containing the pixel values of the $\mu$-th image and let $\mathbf{r}^\mu \in \mathbb{R}^N$ represent the neural response to the $\mu$-th image. The goal in the problem is to learn a collection of weights $\mathbf{W} \in \mathbb{R}^{D \times N}$ which map neural responses $\mathbf{r}^\mu$ to images $\boldsymbol{\theta}^\mu$

$$\boldsymbol{\theta}^\mu \approx \mathbf{W}\mathbf{r}^\mu. \tag{App.76}$$

The generalization error $E_g$ again measures the average error on all points, averaged over all possible datasets $\mathcal{D} = \{(\boldsymbol{\theta}^\mu, \mathbf{r}^\mu)\}_{\mu=1}^P$ of size $P$. If the optimal weights for dataset $\mathcal{D}$ is $\mathbf{W}(\mathcal{D})$ then the generalization error is

$$E_g = \left\langle ||\mathbf{W}(\mathcal{D})\mathbf{r}(\boldsymbol{\theta}) - \boldsymbol{\theta}||^2 \right\rangle_{\boldsymbol{\theta},\mathcal{D}} . \tag{App.77}$$

After identifying eigenfunctions $\phi_k(\boldsymbol{\theta})$, we expand the images in this basis $\boldsymbol{\theta} = \sum_k \mathbf{v}_k \phi_k(\boldsymbol{\theta})$ where $\mathbf{v}_k \in \mathbb{R}^D$. The generalization error is therefore $E_g = \sum_k |\mathbf{v}_k|^2 E_k(P)$ and the cumulative power is $C(k) = \frac{\sum_{\ell<k} |\mathbf{v}_\ell|^2}{\sum_{\ell=1}^{\infty} |\mathbf{v}_\ell|^2}$. We perform this reconstruction task on many filtered versions of the natural scenes. To construct a filter, we first compute the Fourier transform of the image. Let $\mathbf{M}(\boldsymbol{\theta}) \in \mathbb{R}^{\sqrt{D}\times\sqrt{D}}$ represent the non-flattened image and let $\hat{\mathbf{M}}(\boldsymbol{\theta}) \in \mathbb{R}^{\sqrt{D}\times\sqrt{D}}$ represent the Fourier transform of the image, computed explicitly as

$$\hat{M}_{k,k'}(\boldsymbol{\theta}) = D^{-1/4} \sum_{n,m} M_{n,m}(\boldsymbol{\theta}) \exp\Big(2\pi i(nk + mk')/\sqrt{D}\Big) \tag{App.78}$$

To develop the band-pass filter, we calculate $|\mathbf{k}| = \sqrt{k^2 + (k')^2}$ for each of the indices in the matrix. For a band-pass filter with parameters $s_{max}, r$ we simply zero out the entries in $\hat{M}$ which correspond to states with frequencies outside the appropriate band: for any $k, k'$ with $|\mathbf{k}| \notin [\sqrt{s_{max}^2 - r^2}, s_{max}^2]$ then $\hat{M}_{k,k'} \to 0$. We then perform the inverse Fourier transform on $\hat{M}$ to obtain a filtered version of the original image.

### 5 A Simple Feedforward Model of V1

#### 5.1 Linear Neurons

We consider a simplified but instructive model of the V1 population code as a linear-nonlinear map from photoreceptor responses through Gabor filters and then nonlinearity [17, 18, 19]. Let $\mathbf{x} \in \mathbb{R}^2$ represent the two-dimensional retinotopic position of photoreceptors. The firing rates of the photoreceptor at position $\mathbf{x}$ to a static grating stimulus oriented at angle $\theta$ is

$$h(\mathbf{x}, \theta) = \cos\big(\mathbf{k}(\theta) \cdot \mathbf{x}\big) \ , \ \mathbf{k} = \begin{bmatrix} \cos(\theta) \\ \sin(\theta) \end{bmatrix} \in \mathbb{R}^2 \ , \ \theta \in [0, 2\pi]. \tag{App.79}$$

We model each V1 neuron's receptive field as a Gabor filter of the receptor responses $h(\mathbf{x}, \theta)$. The $i$-th V1 neuron has preferred wavevector $\mathbf{k}_i$, generating the following set of weights between photoreceptors and the $i$-th V1 neuron

$$\mathcal{F}(\mathbf{x}, \theta_i) = \frac{\sigma^2}{2\pi} e^{-\frac{\sigma^2}{2}|\mathbf{x}|^2} \cos\big(\mathbf{k}(\theta_i) \cdot \mathbf{x}\big). \tag{App.80}$$

The V1 population code is obtained by filtering the photoreceptor responses. By approximating the resulting sum over all retinal photoreceptors with an integral, we find the response of neuron $i$

to grating stimulus with wavenumber $\mathbf{k}$ is

$$\mathbf{w}(\theta_i) \cdot \mathcal{F}(\theta) = \int \mathcal{F}(\mathbf{x}, \theta_i) h(\mathbf{x}, \theta) d\mathbf{x} = \frac{1}{2} e^{-\frac{1}{2\sigma^2}|\mathbf{k}+\mathbf{k}_i|^2} + \frac{1}{2} e^{-\frac{1}{2\sigma^2}|\mathbf{k}-\mathbf{k}_i|^2}. \tag{App.81}$$

The response of neuron $i$ is computed through nonlinear rectification of this input current $r_i(\theta) = g(\mathbf{w}(\theta_i) \cdot h(\theta))$. For a linear neuron $g(z) = z$, the kernel has the following form

$$K(\theta, \theta') = \frac{\cosh\Big(\cos\big(\theta - \theta'\big)/\sigma^2\Big)}{\cosh\big(\sigma^{-2}\big)}, \tag{App.82}$$

where the kernel is normalized to have maximum value of 1. Note that this normalization of the kernel is completely legitimate since it merely rescales each eigenvalue by a constant and does not change the learning curves.

Since the kernel only depends on the difference between angles $\theta - \theta'$, it is said to posess translation invariance. Such translation invariant kernels admit a Mercer decomposition in terms of Fourier modes $K(\theta) = \sum_n \lambda_n \cos(n\theta)$ since the Fourier modes diagonalize shift invariant integral operators on $\mathbb{S}^1$. For the linear neuron, the kernel eigenvalues scale like $\lambda_n \sim \frac{\beta^n}{2^n n!}$, indicating infinite differentiability of the tuning curves. Since $\lambda_n$ decays rapidly with $n$, we find that this Gabor code has an inductive bias that favors low frequency functions of orientation $\theta$.

#### 5.2 Nonlinear Simple Cells

Introducing nonlinear functions $g(z)$ that map input currents $z$ into the V1 population into firing rates, we can obtain a non-linear kernel $K_g(\theta)$ which has the following definition

$$K_g(\mathbf{k}, \mathbf{k}') = \int p(\mathbf{k}_i) g(\mathcal{F}(\mathbf{k}_i) \cdot \mathbf{h}(\mathbf{k})) g(\mathcal{F}(\mathbf{k}_i) \cdot \mathbf{h}(\mathbf{k}')) d\mathbf{k}_i. \tag{App.83}$$

In this setting, it is convenient to restrict $\mathbf{k}_i, \mathbf{k}, \mathbf{k}' \in \mathbb{S}^1$ and assume that the preferred wavevectors $\mathbf{k}_i$ are uniformly distributed over the circle. In this case, it suffices to identify a decomposition of the composed function $g(\mathbf{w}_i \cdot \mathbf{h}(\theta))$ in the basis of Chebyshev polynomials $T_n(z)$ which satisfy $T_n(\cos(\theta)) = \cos(n\theta)$

$$\begin{aligned} a_n &= \frac{1}{2\pi} \int_0^{2\pi} g\left(e^{-\frac{1}{\sigma^2}} \cosh\left(\frac{1}{\sigma^2}\cos(\theta)\right)\right) \cos(n\theta) d\theta \\ &= \frac{1}{2\pi} \int_{-1}^{1} \frac{1}{\sqrt{1-z^2}} g\left(e^{-\frac{1}{\sigma^2}} \cosh\left(z/\sigma^2\right)\right) T_n(z) dz, \end{aligned} \tag{App.84}$$

which can be computed efficiently with an appropriate quadrature scheme. Once the coefficients $a_n$ are determined, we can compute the kernel by first letting $\theta_i$ to be the angle between $\mathbf{k}$ and $\mathbf{k}_i$

and letting $\theta$ be the angle between $\mathbf{k}$ and $\mathbf{k}'$

$$\begin{aligned} K_g(\theta) &= \int_0^{2\pi} \frac{d\theta_i}{2\pi} \sum_{n,n'} a_n a_{n'} T_n(\cos(\theta_i)) T_{n'}(\cos(\theta_i + \theta)) d\theta_i \\ &= \frac{1}{2} \sum_n a_n^2 \cos(n\theta). \end{aligned} \quad \text{(App.85)}$$

Thus the kernel eigenvalues are $\lambda_n = \frac{1}{2} a_n^2(\psi)$.

*Asymptotic scaling of spectra:* Activation functions that encourage sparsity have slower eigenvalue decays. If the nonlinear activation function has the form $g_{q,t}(z) = \max\{0, z - a\}^q$, then the spectrum decays like $\lambda_n \sim n^{-2q-2}$. A simple argument justifies this scaling: if the function $g(e^{-\sigma^2} \cosh(\sigma^2 z))$ is only $q-1$ times differentiable then $a_n n^q \sim n^{-1}$ since $\sum_n a_n n^q$ must diverge. Therefore $\lambda_n = a_n^2 \sim n^{-2q-2}$. Note that this scaling is independent of the threshold.

#### 5.3 Phase Variation, Complex Cells and Invariance

We can consider a slightly more complicated model where Gabors and stimuli have phase shifts

$$h(\mathbf{x}, \theta, \phi) = \cos\big(\mathbf{k}(\theta) \cdot \mathbf{x} - \phi\big) \ , \ \mathcal{F}(\mathbf{x}, \theta_i, \phi_i) = \frac{\sigma^2}{2\pi} e^{-\frac{\sigma^2}{2}|\mathbf{x}|^2} \cos(\mathbf{k}_i \cdot \mathbf{x} - \phi_i). \quad \text{(App.86)}$$

The simple cells are generated by nonlinearity

$$r_i(\theta, \phi) = g(\mathcal{F}(\theta_i, \phi_i) \cdot \mathbf{h}(\theta, \phi)). \quad \text{(App.87)}$$

The input currents into the simple V1 cells can be computed exactly

$$\mathbf{w}(\theta_i, \phi_i) \cdot \mathcal{F}(\theta, \phi) = \big\langle \cos(\mathbf{k}_i \cdot \mathbf{x} - \phi_i) \cos(\mathbf{k} \cdot \mathbf{x} - \phi) \big\rangle_{\mathbf{x} \sim \mathcal{N}(0, \sigma^2 \mathbf{I})} \cdot \quad \text{(App.88)}$$

$$= \frac{1}{2} \cos(\phi + \phi_i) e^{-\frac{1}{2\sigma^2}|\mathbf{k} + \mathbf{k}_i|^2} + \frac{1}{2} \cos(\phi - \phi_i) e^{-\frac{1}{2\sigma^2}|\mathbf{k} - \mathbf{k}_i|^2}. \quad \text{(App.89)}$$

When $|\mathbf{k}| = |\mathbf{k}_i| = 1$, the simple cell tuning curves $r_i = g(\mathbf{w}_i \cdot \mathbf{h})$ only depend on $\cos(\theta - \theta_i)$ and $\phi$, allowing a Fourier decomposition

$$r_i(\theta, \phi) = \sum_n a_n(\phi, \phi_i) \cos\big(n(\theta - \theta_i)\big)). \quad \text{(App.90)}$$

The simple cell kernel $K_s$, therefore decomposes into Fourier modes over $\theta$

$$K_s(\theta, \theta', \phi, \phi') = \sum_n b_n(\phi, \phi') \cos\big(n(\theta - \theta')\big), \quad \text{(App.91)}$$

where $b_n(\phi, \phi') = \big\langle a_n(\phi, \phi_i) a_n(\phi', \phi_i) \big\rangle_{\phi_i}$. It therefore suffices to solve the infinite sequence of integral eigenvalue problems over $\phi$

$$\frac{1}{2\pi}\int_0^{2\pi} b_n(\phi,\phi')v_{n,k}(\phi)d\phi = \lambda_{n,k}v_{n,k}(\phi') \implies K_s(\theta,\theta',\phi,\phi') = \sum_{n,k}\lambda_{n,k}\cos\big(n(\theta-\theta')\big)v_{n,k}(\phi)v_{n,k}(\phi'). \tag{App.92}$$

With this choice it is straightforward to verify that the kernel eigenfunctions are $v_{n,k}(\theta,\phi) = e^{in\theta}v_{n,k}(\phi)$ with corresponding eigenvalue $\lambda_{n,k}$. Since $b_n$ is not translation invariant in $\phi - \phi'$, the eigenfunctions $v_{n,k}$ are not necessarily Fourier modes. These eigenvalue problems for $b_n$ must be solved numerically when using arbitrary nonlinearity $\psi$. The top eigenfunctions of the simple cell kernel depend heavily on the phase of the two grating stimuli $\phi$. Thus, a pure orientation discrimination task which is independent of phase requires a large number of samples to learn with the simple cell population.

#### 5.4 Complex Cells Populations are Phase Invariant

V1 also contains complex cells which possess invariance to the phase $\phi$ of the stimulus. Again using Gabor filters

$$\mathcal{F}(\mathbf{x},\theta_i,\phi_i) = \frac{\sigma^2}{2\pi}e^{-\frac{\sigma^2}{2}|\mathbf{x}|^2}\cos\big(\mathbf{k}(\theta_i)\cdot\mathbf{x}-\phi_i\big), \tag{App.93}$$

we model the complex cell responses with a quadratic nonlinearity and sum over two squared filters which are phase shifted by $\pi/2$

$$r_i(\theta,\phi) = (\mathcal{F}(\theta_i,\phi_i)\cdot\mathbf{h}(\theta,\phi))^2+(\mathcal{F}(\theta_i,\phi_i-\pi/2)\cdot\mathbf{h}(\theta,\phi))^2 = \frac{1}{4}e^{-\frac{1}{\sigma^2}|\mathbf{k}+\mathbf{k}_i|^2}+\frac{1}{4}e^{-\frac{1}{\sigma^2}|\mathbf{k}-\mathbf{k}_i|^2}+\frac{1}{2}e^{-\sigma^2}\cos(2\phi_i), \tag{App.94}$$

which we see is independent of the phase $\phi$ of the grating stimulus. Integrating over the set of possible Gabor filters $(\mathbf{k}_i,\phi_i)$ again gives the following kernel for the complex cells

$$K_c(\theta) = \frac{1}{\cosh(2\beta)}\cosh\big(2\beta\cos(\theta)\big). \tag{App.95}$$

Remarkably, this kernel is independent of the phase $\phi$ of the grating stimulus. Thus, complex cell populations possess good inductive bias for vision tasks where the target function only depends on the orientation of the stimulus rather than it's phase. In reality, V1 is a mixture of simple and complex cells. Let $s \in [0,1]$ represent the relative proportion of neurons which are simple cells and $(1-s)$ the relative proportion of complex cells. The kernel for the mixed V1 population is given by a simple convex combination of the simple and complex cell kernels

$$\begin{aligned} K_{V1}(\theta,\theta',\phi,\phi') &= \frac{1}{N}\sum_{i=1}^N r_i(\theta,\phi)r_i(\theta',\phi') \to \left\langle r(\theta,\phi,c)r(\theta',\phi',n)\right\rangle_{n\sim p_{V1}(n)} \\ &= s\left\langle r(\theta,\phi,n)r(\theta',\phi',n)\right\rangle_{n\sim p_s(n)} + (1-s)\left\langle r(\theta,\phi,n)r(\theta',\phi',n)\right\rangle_{n\sim p_c(n)} \\ &= sK_s(\theta,\theta',\phi,\phi') + (1-s)K_c(\theta,\theta'), \end{aligned} \tag{App.96}$$

where $n$ denotes neuron type (simple vs complex, tuning etc) and $P_{V1}(n), p_s(n), p_c(n)$ are probability distributions over the V1 neuron identities, the simple cell identities and the complex cell identities respectively. Increasing $s$ increases the phase dependence of the code by giving greater weight to the simple cell population. Decreasing $s$ gives weight to the complex cell population, encouraging phase invariance of readouts.

##### 5.5 Visualization of Feedforward Gabor V1 Model and Induced Kernels

Examples of the induced kernels for the Gabor-bank V1 model are provided in Figure App.4. We show how choice of rectifying nonlinearity $g(z)$ and sparsifying threshold $a$ influence the kernel and their spectra. Learning curves for simple orientation tasks are provided.

#### 6 Time Dependent Neural Codes

##### 6.1 RNN Model and Decomposition

In this setting, the population code $\mathbf{r}(\{\boldsymbol{\theta}(t)\}, t)$ is a function of an input stimulus sequence $\boldsymbol{\theta}(t)$ and time $t$. In general the neural code $\mathbf{r}$ at time $t$ can depend on the entire history of the stimulus input $\boldsymbol{\theta}(t')$ for $t' \leq t$, as is the case for recurrent neural networks. We denote dependence of a function $f$ on $\boldsymbol{\theta}(t)$ in this causal manner with the notation $f(\{\boldsymbol{\theta}\}, t)$. In a learning task, a set of readout weights $\mathbf{w}$ are chosen so that a downstream linear readout $f(\{\boldsymbol{\theta}\}, t) = \mathbf{w} \cdot \mathbf{r}(\{\boldsymbol{\theta}\}, t)$ approximates a target sequence $y(\{\boldsymbol{\theta}\}, t)$ which maps input stimulus sequences to output scalar sequences. The quantity of interest is the generalization $E_g$, which in this case is an average over both input sequences and time, $E_g = \left\langle (y(\{\boldsymbol{\theta}\}, t) - f(\{\boldsymbol{\theta}\}, t))^2 \right\rangle_{\boldsymbol{\theta}(t), t}$. The average is computed over a distribution of input stimulus sequences $p(\boldsymbol{\theta}(t))$. To train the readout, $\mathbf{w}$, the network is given a sample of $P$ stimulus sequences $\boldsymbol{\theta}^\mu(t), \mu = 1, ..., P$. For the $\mu$-th training input sequence, the target system $y$ is evaluated at a set of discrete time points $\mathcal{T}_\mu = \{t_1, t_2, ..., t_{|\mathcal{T}_\mu|}\}$ giving a collection of target values $\{y_t^\mu\}_{t \in \mathcal{T}_\mu}$ and a total dataset of size $\mathcal{P} = \sum_{\mu=1}^P |\mathcal{T}_\mu|$. The *average case generalization* computes a further average of the generalization error $E_g$ over randomly sampled datasets of size $\mathcal{P}$.

Learning is again achieved through iterated weight updates with delta-rule form, but now have contributions from both sequence index and time $\Delta \mathbf{w} = \eta \sum_\mu \sum_{t \in \mathcal{T}_\mu} \mathbf{r}_t^\mu (y_t^\mu - f_t^\mu)$. As before, optimization of the readout weights is equivalent to kernel regression with a kernel that computes inner products of neural population vectors at different times $t, t'$ for different input sequences $\{\boldsymbol{\theta}\}, \{\boldsymbol{\theta}'\}$: $K(\{\boldsymbol{\theta}\}, \{\boldsymbol{\theta}'\}, t, t') = \frac{1}{N} \mathbf{r}(\{\boldsymbol{\theta}\}, t) \cdot \mathbf{r}(\{\boldsymbol{\theta}\}, t')$ . This kernel depends on details of the time varying population code including its recurrent intrinsic dynamics as well as its encoding of the time-varying input stimuli. The optimization problem and delta rule described above converge to the kernel regression solution for kernel gram matrix $K_{t,t'}^{\mu,\mu'} = \frac{1}{N} \mathbf{r}_t^\mu \cdot \mathbf{r}_{t'}^{\mu'}$ [20, 21, 22]. The learned function has the form $f(\{\boldsymbol{\theta}\}, t) = \sum_{\mu, t' \in \mathcal{T}_\mu} \alpha_t^\mu K(\{\boldsymbol{\theta}\}, \{\boldsymbol{\theta}\}^\mu, t, t')$, where $\boldsymbol{\alpha} = \mathbf{K}^+ \mathbf{y}$ for kernel gram matrix $\mathbf{K} \in \mathbb{R}^{\mathcal{P} \times \mathcal{P}}$ which is computed for the entire set of training sequences, and the vector $\mathbf{y} \in \mathbb{R}^{\mathcal{P}}$ is the vector containing the desired target outputs for each sequence. Assuming a probability distribution over sequences $\boldsymbol{\theta}(t)$, the kernel can be diagonalized with orthonormal eigenfunctions $\psi_k(\{\boldsymbol{\theta}\}, t)$. Our theory carries over from the static case: kernels whose top eigenfunctions have high alignment with the target dynamical system $y(\{\boldsymbol{\theta}\}, t)$ will achieve the best average case generalization performance.

### Appendix Figures

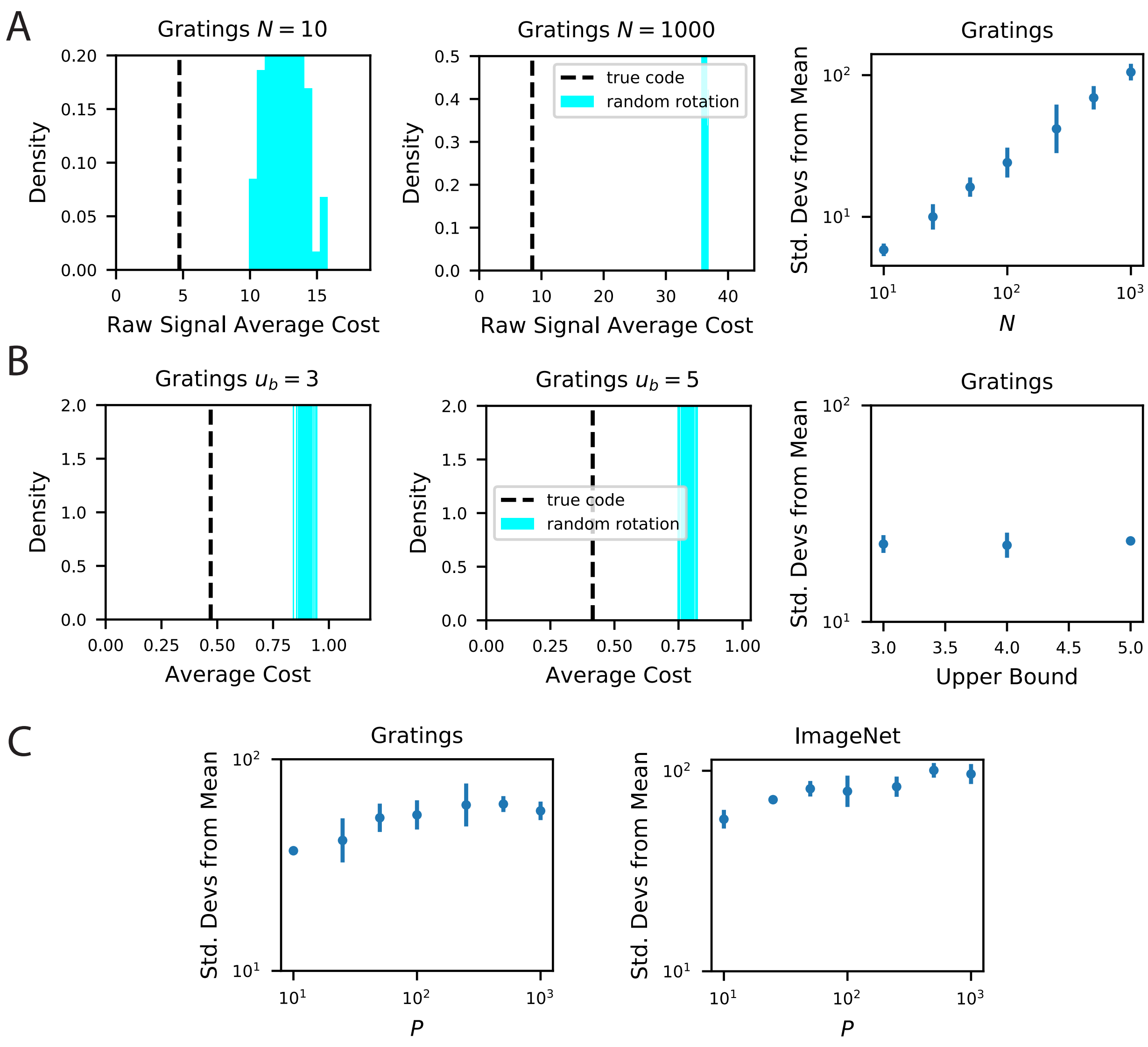


Figure App.1: Our metabolic efficiency finding is robust to different pre-processing techniques and upper bounds on neural firing. **A** We show the same result as the main text except we use raw (non $z$-scored) estimate of responses for each stimulus. **B** Our result is robust to imposition of firing rate upper bounds $u_b$ on each neuron. This result uses the $z$-scored responses to be consistent with the rest of the paper. The biological code achieves a maximum $z$-score values in the range $[3.2, 4.7]$, which motivated the range of our tested upper bound values $\{3, 4, 5\}$. **C** Our finding is robust to the number of sampled stimuli $P$ as we show in an experiment where rotations in $N = 500$ dimensional subspace.

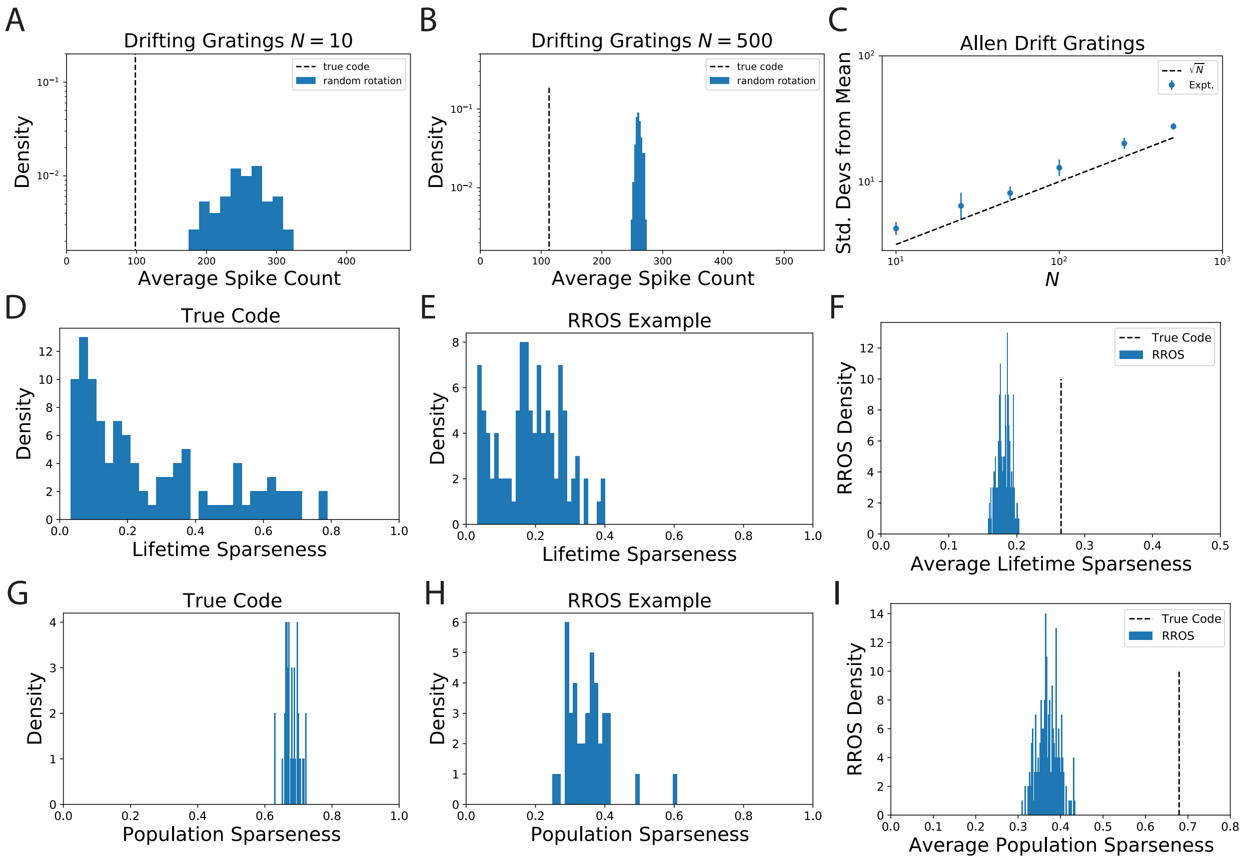


Figure App.2: The observation that randomly oriented codes with the same kernel require higher spike counts than the original code is reproduced from electrophysiological recordings of Mouse visual cortex (VISp and VISal) from the AIBO. **A** Distribution of average spike count for a random selection of $N = 10$ neurons. **B** The same for $N = 500$ randomly selected neurons. **C** Since the distribution of average spike count for the randomly rotated codes concentrates with $N$ the number of standard deviations the true code is from the mean increases with $N$. We show that this scaling is approximately like $\sqrt{N}$ which suggests that the variability in average spike count for randomly rotated codes goes scales with neuron count like $\frac{1}{\sqrt{N}}$. This is intuitive since the average spike count over the rotated code is an empirical average over $N$ random variables. **D** The lifetime sparseness of neurons in the true code are spread out over a large range. **E** The lifetime sparseness distribution for an example RROS code does not have the same range. **F** The average (over neurons) lifetime sparseness of RROS codes over random rotations is significantly lower than the average lifetime sparseness of the true code. **G-I** The true visual cortex code has much higher population sparseness over each grating stimulus as well

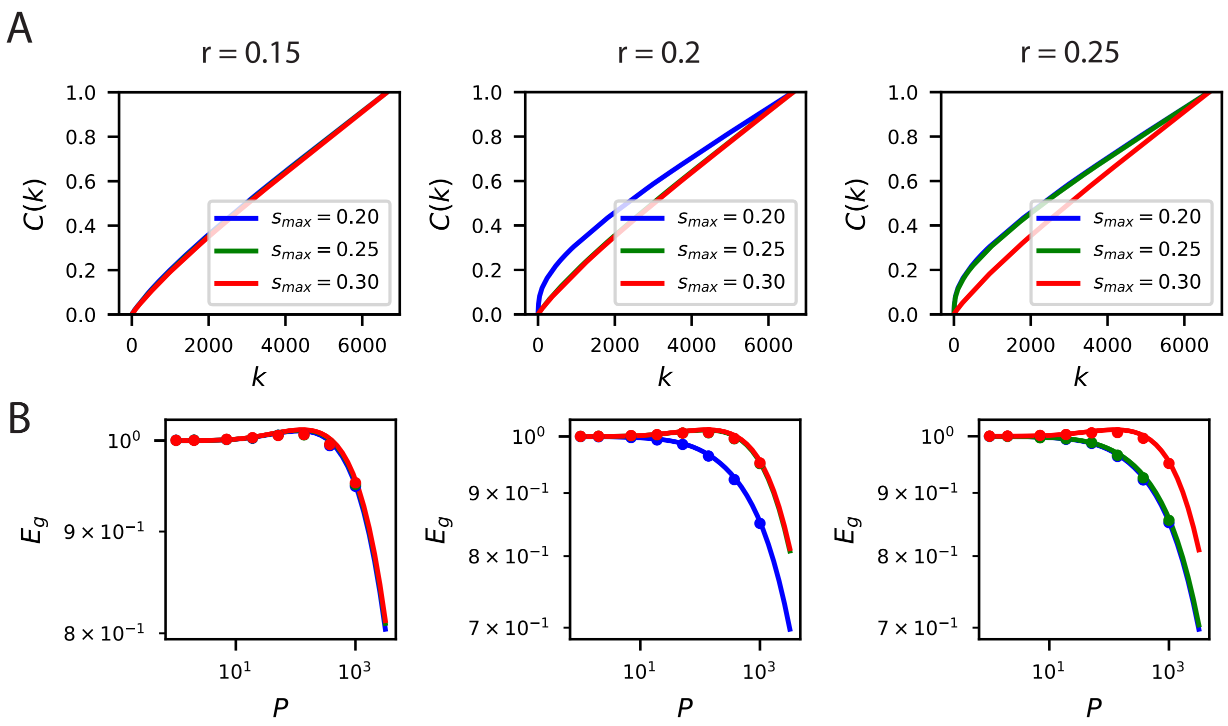


Figure App.3: The inductive bias to reconstruct low spatial frequency components of natural scenes from the population responses holds over several band-pass filters of the form $|\mathbf{k}| \in \left[\max(\sqrt{s_{max}^2 - r^2}, 0), s_{max}\right]$. **A** The cumulative power curves for reconstruction of the filtered images with different values of $c$ and $s_{max}$. Lower values of $s_{max}$ and higher values of $c$ preserve mostly low frequency content in the images and are easier to reconstruct from the neural responses. **B** The learning curves respect the ordering of the cumulative power metric $C(k)$.

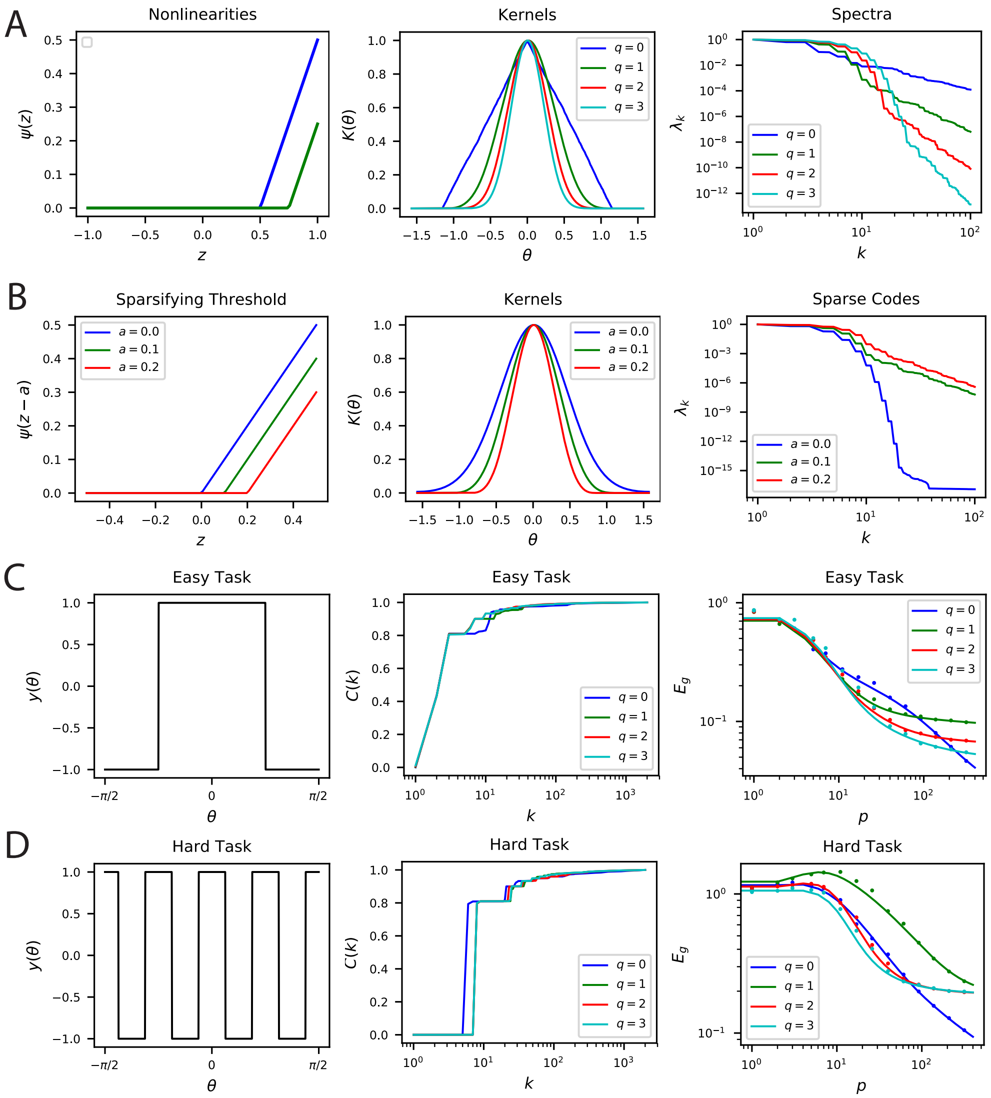


Figure App.4: Nonlinear Rectification and proportion of simple and complex cells influences the inductive bias of the population code. **A** The choice of nonlinearity has influence on the kernel and its spectrum. If the nonlinearity is $g(z) = \max(0, z^q)$, then $\lambda_k \sim k^{-2n-2}$. **B** The sparsity can be increased by shifting the nonlinearity $g(z) \to g(z-a)$. Sparser codes have higher dimensionality. Note that $a = 0$ is a special case where the neurons behave in the linear regime for all inputs $\theta$ since the currents $\mathbf{w} \cdot \mathbf{h}$ are positive. Thus, for $a = 0$, the spectrum decays like a Bessel Function $\lambda_k = I_k(\beta)$. **C-D** Easy and hard orientation discrimination tasks with varying nonlinear polynomial order $q$. At low sample sizes, large $q$ performs better, whereas at large $P$, the step function nonlinearity $q = 0$ achieves the best performance.

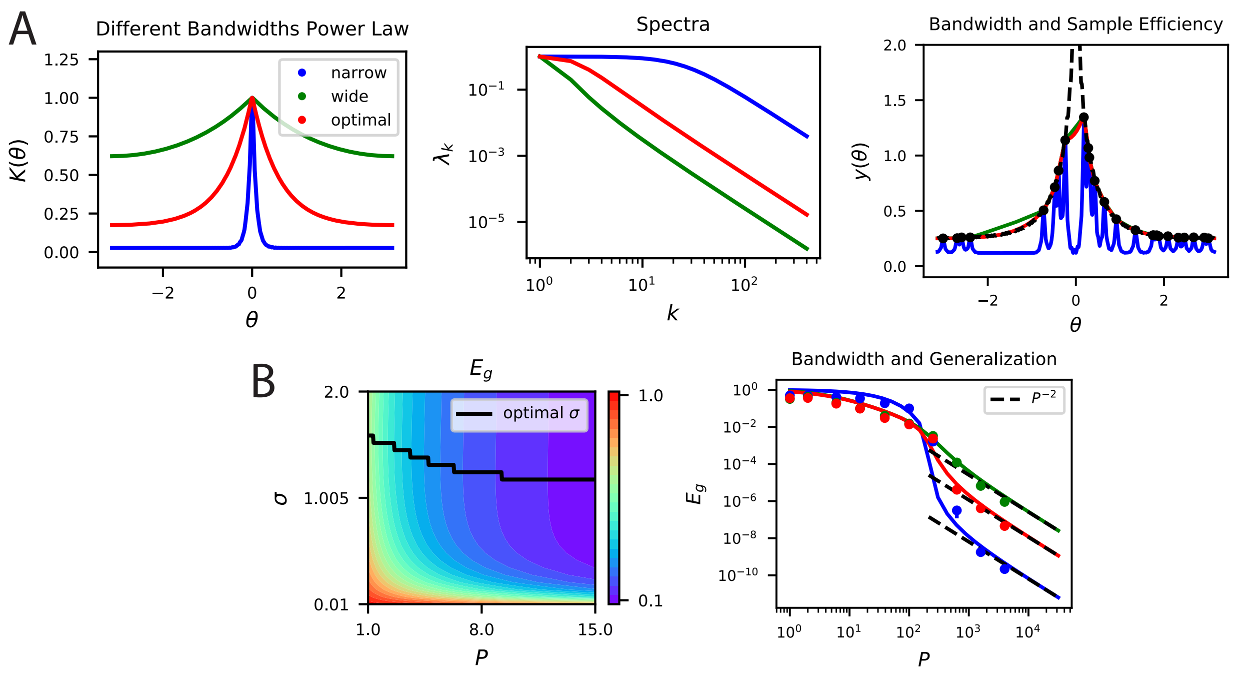


Figure App.5: **A, B** Kernel regression experiments are performed with Laplace kernels of varying bandwidth on a non-differentiable target function. The top eigenvalues are modified by changing the bandwidth, but the asymptotic power law scaling is preserved. Generalization at low $P$ is shown in the contour plot while the large $P$ scaling is provided in the generalization. In A-right and B-right, color code is the same as in the main text.

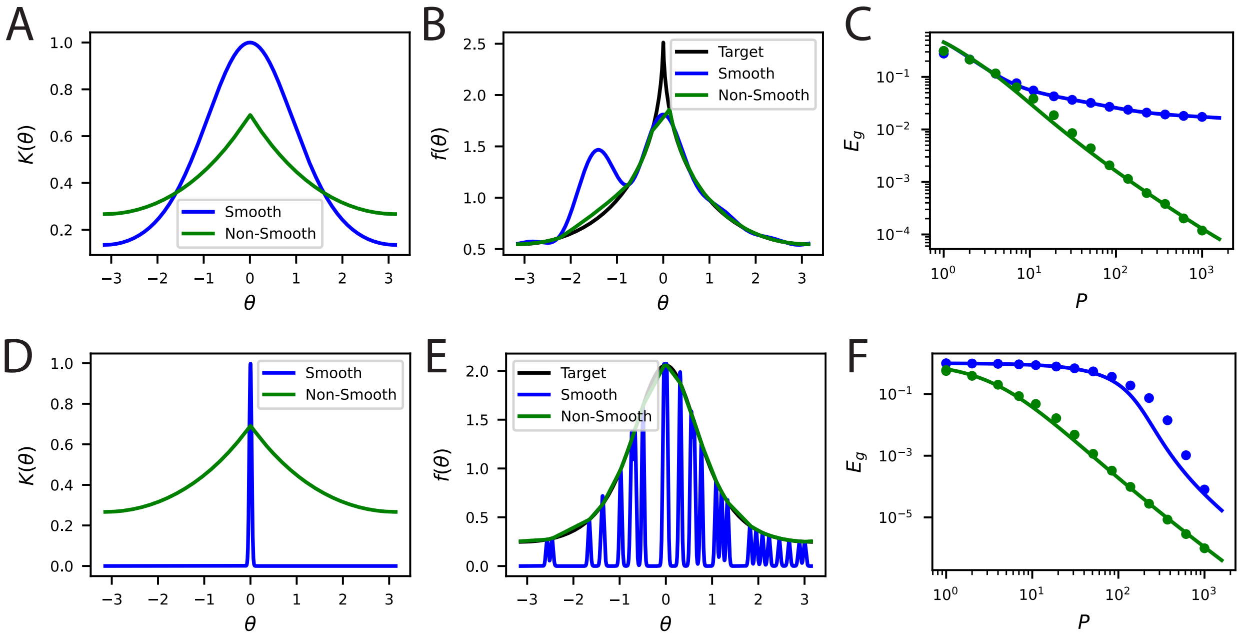


Figure App.6: Non-differentiable kernels can generalize better than infinitely differentiable kernels in a variety of contexts. **A** An infinitely differentiable von-Mises kernel (blue) will be compared to a non-differentiable Laplace kernel (green). **B** For a non-differentiable target function $y(\boldsymbol{\theta})$ with a cusp, the non-smooth kernel can provide a better fit to the target function at $P = 30$ samples. **C** The learning curves for this task. Solid lines are theory, circles are simulations. **D** The lengthscale of the kernel can be more important than local smoothness. We will now compare a narrow von-Mises kernel (still infinitely differentiable) with a non-differentiable wide kernel. **E** The non-smooth kernel generalizes on a smooth task almost perfectly whereas the narrow smooth kernel only locally interpolates. **F** The learning curves for this smooth task.

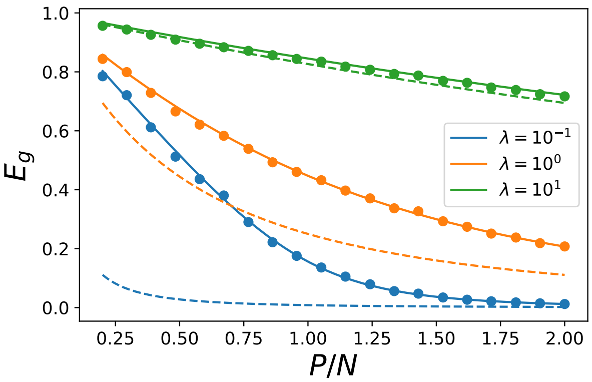


Figure App.7: Our theoretical prediction for generalization error $E_g$ (solid lines) on a linear kernel $K(\mathbf{x}, \mathbf{x}') = \mathbf{x} \cdot \mathbf{x}'$ regression problem with uncorrelated Gaussian design $\mathbf{x} \sim \mathcal{N}(0, \mathbf{I})$ and linear teacher $y = \beta \cdot \mathbf{x}$ in $N = 500$ dimensions. The generalization error as a function of sample size $P$ is plotted predicted by the equivalent kernel/Wiener filter (dashed). Each color represents a different regularization level $\lambda$. The experimental generalization is shown as dots of the corresponding color and shows the numerical error obtained from solving the kernel regression problem with regularization level $\lambda$. We see that the two theories coincide at large $\lambda$ but are drastically different at low levels of explicit regularization $\lambda$.

[5] Trevor Hastie, Andrea Montanari, Saharon Rosset, and Ryan J. Tibshirani. Surprises in high-dimensional ridgeless least squares interpolation, 2020.

[6] Mikhail Belkin, Daniel Hsu, Siyuan Ma, and Soumik Mandal. Reconciling modern machine-learning practice and the classical bias–variance trade-off. *Proceedings of the National Academy of Sciences*, 116(32):15849–15854, 2019.

[7] Bernard Widrow and Marcian E. Hoff. Adaptive switching circuits. In *1960 IRE WESCON Convention Record, Part 4*, pages 96–104, New York, 1960. IRE.

[8] John Hertz, Anders Krough, and Richard Palmer. *Introduction To The Theory Of Neural Computation*, volume 44. 01 1991.

[9] Bernhard Schölkopf, Ralf Herbrich, and Alex J. Smola. A generalized representer theorem. In *Proceedings of the 14th Annual Conference on Computational Learning Theory and and 5th European Conference on Computational Learning Theory*, COLT '01/EuroCOLT '01, page 416–426, Berlin, Heidelberg, 2001. Springer-Verlag.

[10] Abdulkadir Canatar, Blake Bordelon, and Cengiz Pehlevan. Spectral bias and task-model alignment explain generalization in kernel regression and infinitely wide neural networks. *Nature Communications*, in press, 2021.

[11] Blake Bordelon, Abdulkadir Canatar, and Cengiz Pehlevan. Spectrum dependent learning curves in kernel regression and wide neural networks. In Hal Daumé III and Aarti Singh, editors, *Proceedings of the 37th International Conference on Machine Learning*, volume 119 of *Proceedings of Machine Learning Research*, pages 1024–1034. PMLR, 13–18 Jul 2020.

[12] Peter Sollich. Learning curves for gaussian processes. In *Neurips*, 1998.

[13] Peter Sollich. Gaussian process regression with mismatched models. In T. Dietterich, S. Becker, and Z. Ghahramani, editors, *Advances in Neural Information Processing Systems*, volume 14, pages 519–526. MIT Press, 2002.

[14] Thomas Kailath. Correlation detection of signals perturbed by a random channel. *IRE Transactions on Information Theory*, 6(3):361–366, 1960.

[15] Simon Haykin. Adaptive filter theory. prentice-hall, 1991.

[16] Andreas Engel and Christian Van den Broeck. *Statistical mechanics of learning.* Cambridge University Press, 2001.

[17] E. Adelson and J. Bergen. Spatiotemporal energy models for the perception of motion. *Journal of the Optical Society of America. A, Optics and image science*, 2 2:284–99, 1985.

[18] Bruno A. Olshausen and David J. Field. Sparse coding with an overcomplete basis set: A strategy employed by v1? *Vision Research*, 37(23):3311–3325, 1997.

[19] Oleg Rumyantsev, Jérôme Lecoq, Oscar Hernandez, Yanping Zhang, Joan Savall, Radosław Chrapkiewicz, Jane Li, Hongkui Zeng, Surya Ganguli, and Mark Schnitzer. Fundamental bounds on the fidelity of sensory cortical coding. *Nature*, 580:1–6, 04 2020.

[20] Jonathan Dong, Ruben Ohana, Mushegh Rafayelyan, and Florent Krzakala. Reservoir computing meets recurrent kernels and structured transforms, 2020.

[21] Greg Yang. Tensor programs i: Wide feedforward or recurrent neural networks of any architecture are gaussian processes, 2019.

[22] Greg Yang. Tensor programs ii: Neural tangent kernel for any architecture, 2020.
